# Control of intestinal stemness and cell lineage by histone variant H2A.Z isoforms

**DOI:** 10.1101/2023.10.10.561706

**Authors:** Jérémie Rispal, Clémence Rives, Virginie Jouffret, Caroline Leoni, Louise Dubois, Martine Chevillard-Briet, Didier Trouche, Fabrice Escaffit

## Abstract

The histone variant H2A.Z plays important functions in the regulation of gene expression. In mammals, it is encoded by two genes, giving raise to two highly related isoforms named H2A.Z.1 and H2A.Z.2, which can have similar or antagonistic functions depending on the promoter. Knowledge of the physiopathological consequences of such functions emerges, but how the balance between these isoforms regulates tissue homeostasis is not fully understood. Here, we investigated the relative role of H2A.Z isoforms in intestinal epithelial homeostasis. Through genome-wide analysis of H2A.Z genomic localization in differentiating Caco-2 cells, we uncovered an enrichment of H2A.Z isoforms on the bodies of genes which are induced during enterocyte differentiation, stressing the potential importance of H2A.Z isoforms dynamics in this process. Through a combination of *in vitro* and *in vivo* experiments, we further demonstrated the two isoforms cooperate for stem and progenitor cells proliferation, as well as for secretory lineage differentiation. However, we found that they antagonistically regulate enterocyte differentiation, with H2A.Z.1 preventing terminal differentiation and H2A.Z.2 favoring it. Altogether, these data indicate that H2A.Z isoforms are critical regulators of intestine homeostasis and may provide a paradigm of how the balance between two isoforms of the same chromatin structural protein can control physiopathological processes.

## Introduction

Chromatin dynamics is crucial for organ development, including the control of proliferation, cell fate determination and tissue homeostasis, as well as for its plasticity and adaptive abilities to stresses (Rispal *et al*., 2020). The role of chromatin components in the establishment of genetic programs controlling tissue homeostasis is a very partially explored field. Due to their positioning, in closed proximity with DNA, and the regulations which can affect them, histones are central elements in the control of numerous biological processes, such as transcription, replication or DNA damage repair. Integrated in signaling pathways and functionally associated with specific transcription factors, canonical histones and their variants participate in cell fate. For instance, incorporation into chromatin of one of the most conserved histone variant, H2A.Z, has been shown to exerts an important role in various biological events, including the licensing of replication origins (Long *et al*., 2020), the chronobiological regulations (Tong *et al*., 2020) or homeostatic processes, such as in mammalian intestine (Rispal *et al*., 2019; Zhao *et al*., 2019), in worm gonads (Shibata *et al*., 2014) as well as in plant flowering (Carter *et al*., 2018). Moreover, more and more evidences give to H2A.Z a key role in the epithelial-to-mesenchymal transition (EMT) in physiologic embryonic conditions as well as in pathologies (Domaschenz *et al*., 2017; Lin and Wu, 2020). Mechanisms driving the pleiotropic impacts of H2A.Z are complex and remain to be elucidated. H2A.Z histone variant has been shown to regulate the transcription of many genes (Scacchetti and Becker, 2020), activating or repressing subsets of genes, depending on its post-translational modifications which can trigger the binding of key regulators such as BET family (Patel *et al*., 2021), as well as on its interactors (Lamaa *et al*., 2020). The dynamics of its incorporation is a key determinant for those regulations (Murphy *et al*., 2020) participating to the H2A.Z’s “social” network recently described by Hake’s lab (Kreienbaum *et al*., 2022).

Since the intestinal epithelium is one of the most rapidly regenerating tissue, whose complete renewal occurs each 3 to 5 days, it is a widely used model to study homeostasis regulatory processes which have to be tightly coordinated. Several chromatin-related mechanisms and signaling pathways are involved in the control of the intestinal stem cells (ISC) self-renewal at the bottom of the crypts (Barker *et al*., 2007; Rispal *et al*., 2020). They are also important for the generation of progenitor cells and their subsequent commitment, thus giving rise to absorptive or secretory lineages, whose distinct functions are critical for the physiology of the mature organ. For instance, the Wnt signaling activity, in the crypt niche, is essential for the stem cell maintenance, and the Notch pathway is crucial for the commitment into lineages, thanks to the equilibrium between MATH1 and HES1 transcription factors (Sancho *et al*., 2015). Recently, it has also been demonstrated that the impairment of epithelial TNF pathway alters the secretory lineage by increasing the differentiation of progenitors to goblet cells, then causing an excess in mucus production (Reyes *et al*., 2023).

In this context, it has been previously found that H2A.Z histone variant incorporation into chromatin can participate in repression of Notch-target genes (Giaimo *et al*., 2018). In our group, we demonstrated that the H2A.Z.1 isoform, encoded by *H2AFZ* gene, is a major effector of the Wnt pathway in the maintenance of stem cells and progenitors, as well as an important repressor of the differentiation marker expression of the absorptive lineage (Rispal *et al*., 2019).

Moreover, Kazakevych *et al*. (Kazakevych *et al*., 2017) showed that transcriptome modification correlates with the decrease of H2A.Z expression during epithelial cells differentiation. Consistent with this finding, we previously showed that the major isoform H2A.Z.1 is involved in the maintenance of proliferative and undifferentiated state of the crypt cells (Rispal *et al*., 2019).

However, the role of H2A.Z.2, another isoform of H2A.Z which is encoded by the *H2AFV* paralog gene, and which diverges from H2A.Z.1 by only 3 amino acids (Eirin-Lopez *et al*., 2009), is totally unknown in such context. Recently, some studies have shown specific roles for each of these isoforms in the control of gene expression in melanoma cells (Vardabasso *et al*., 2015), neurons (Dunn *et al*., 2017) or neural crest cells (Greenberg *et al*., 2019). Moreover, using genome editing strategy to bypass the lack of isoform-specific antibody, we demonstrated, in a genome-wide study, that the balance between both isoforms determines the regulation of specific gene expression (Lamaa *et al*., 2020). Other isoform-specific roles have been recently characterized, such as the essential transcription-independent contribution of H2A.Z.2 to chromatids cohesion and chromosomes segregation, for example (Sales-Gil *et al*., 2021), even if deciphering the role of each isoform of the same histone variant remains challenging (Cheema *et al*., 2020).

In this study, we analyzed the functional specificities or redundancy of H2A.Z.1 and H2A.Z.2 isoforms in the intestinal epithelial homeostasis. Using genome-wide approaches, we characterized the binding of each isoform on promoters during Caco-2/15 clone differentiation and correlated this binding with gene expression changes. In combination with gain or loss of function experiments, we next found that both isoforms are redundant in promoting the proliferation and in the inhibition of secretory cell lineage, but that they exert opposite effects on the control of the enterocyte gene expression. Our results thus show how isoforms of a same histone variant specifically trigger homeostasis and cell fate.

## Results

### Gene activation during intestinal epithelial differentiation correlates with the enrichment of H2A.Z isoforms on gene bodies

In previous works, we and others demonstrated that H2A.Z histone variant isoforms can specifically regulate different subsets of genes (Lamaa *et al*., 2020; Tang *et al*., 2020; Sales-Gil *et al*., 2021). Moreover, a specific role of each isoform was also demonstrated for craniofacial morphogenesis (Greenberg *et al*., 2019) and brain development (Dunn *et al*., 2017). However, little is known about the relative contribution of each isoform to epithelial differentiation process related to tissue homeostasis.

Caco-2/15 cells is a useful model to study the intestinal enterocyte-type differentiation process, since it spontaneously acquired lineage-specific differentiation features upon confluence (Van Beers *et al*., 1995). To decipher the contribution of each H2A.Z isoform to gene expression pattern modifications that occur during intestinal epithelial differentiation, we first generated two Caco-2/15 cell-based models in which we heterozygously introduced a Flag tag into the 3’ regions of either *H2AFZ* (Caco-2 Z1flag cell line) or *H2AFV* (Caco-2 Z2flag cell line) genes, by CRISPR-Cas9 genome editing, to bypass the lack of isoform-specific antibody. These models allowed us to follow, during the differentiation process, H2A.Z isoforms expression (SuppFig.1A and 1B) and their incorporation into chromatin.

**Figure 1:**
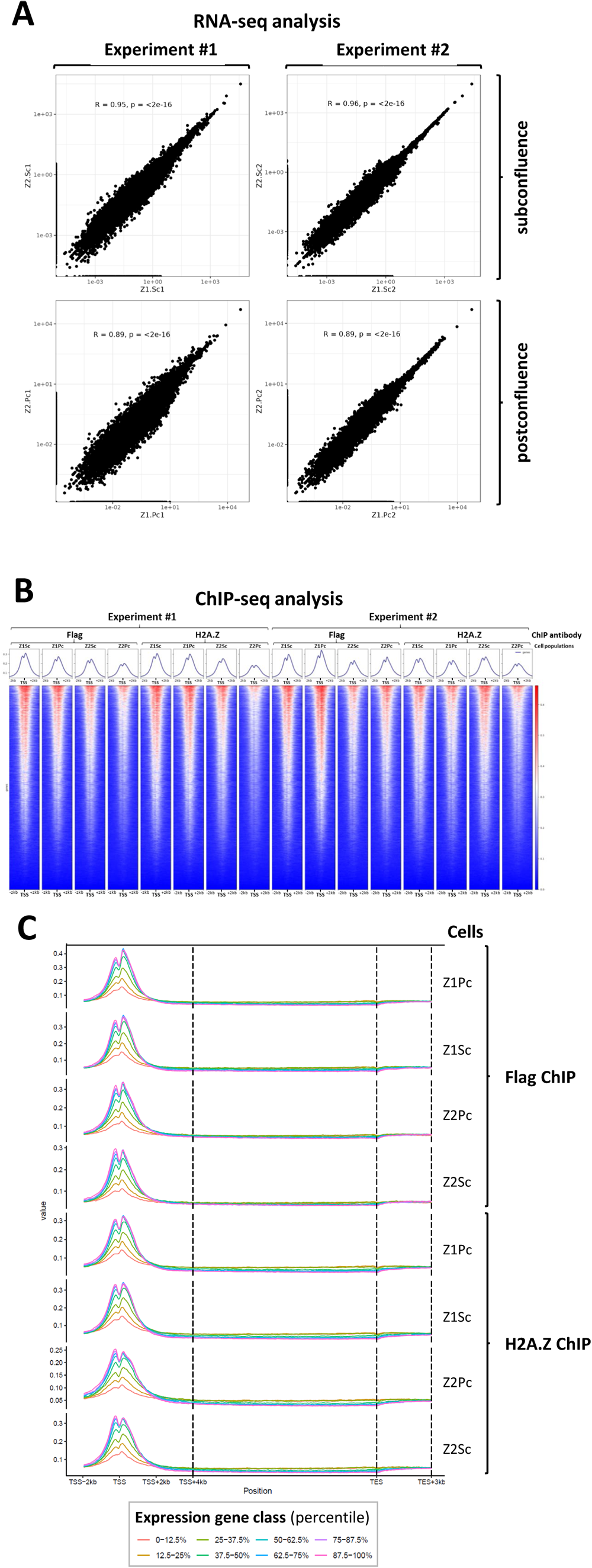
Validation of CRISPR-flagged Caco-2/15 cell lines cells and correlation between isoform recruitment and gene expression. **A:** mRNA from subconfluent or 7-days postconfluent Z1-Flag or Z2-Flag Caco-2/15 cells were subjected to RNA-seq analysis and a DESEQ2 standardisation was done. Correlations of RPKM counts between cell lines, either at subconfluent (upper panels) or postconfluent (lower ones) stages, in two RNA-seq experiments (left and right) are shown. Spearman correlation was done and indicated on graphs. **B:** heatmaps for Transcription Start Site regions (−2kb to +2kb from the TSS) representing the enrichment in H2A.Z.1-Flag, H2A.Z.2-Flag or total H2A.Z for subconfluent (Sc) or postconfluent (Pc) Caco-2/15 cells with Flag-tagged H2A.Z.1 (Z1) or H2A.Z.2 (Z2), as indicated. Data from two ChIP-seq experiments have been analyzed and an heatmap classifying genes by the intensity of total H2A.Z signal around TSS is shown. The corresponding metaprofile of the analyzed TSS region is also presented above each heatmap. **C:** Enrichment in H2A.Z.1-Flag, H2A.Z2-Flag or total H2A.Z all along genes depending on their expression levels. Genes were dispatched in eight classes of equal size and the average metaprofile of ChIP-seq data (from subconfluent (Sc) or postconfluent (Pc) genome-edited Caco-2/15 cells, as indicated) is shown for each class.

Using these models, we first performed RNA-seq experiments in duplicates from cell cultures in subconfluent (Sc) or 7-days post-confluent (Pc) state, allowing us to identify subsets of genes that are differentially expressed during differentiation (see SuppFig.2A and 2B for PCA analysis and volcano plots, respectively). We found a strong correlation in gene expression between the two cell lines, both in subconfluent and post-confluent cells (Fig.1A), indicating that gene expression and differentiation abilities are highly similar in the two cell lines.

Combining RNA-Seq data from the two models, we found that most of significantly regulated genes are repressed in post-confluent cells (13 591 genes), probably due to lower proliferation rate and higher commitment towards a specific lineage, which would together result in the repression of proliferation-linked genes and of genes specific to other lineages. Analyzing the TOP1000 of these repressed genes (SuppFig.2C), we observed, as expected, a strong link of such genes to cell proliferation (cell cycle (such as cyclins/CDK families) or DNA replication (MCM family)) or to mechanisms depending on cell cycle (homologous recombination (for example RAD51C)). Among upregulated genes (1 355 genes), the TOP1000 subset is linked to signaling pathways activation and metabolism of lipids (APO family), amino-acids (SAT family) and carbohydrates (ALDOC), which are expected regulations during enterocyte-like differentiation.

**Figure 2:**
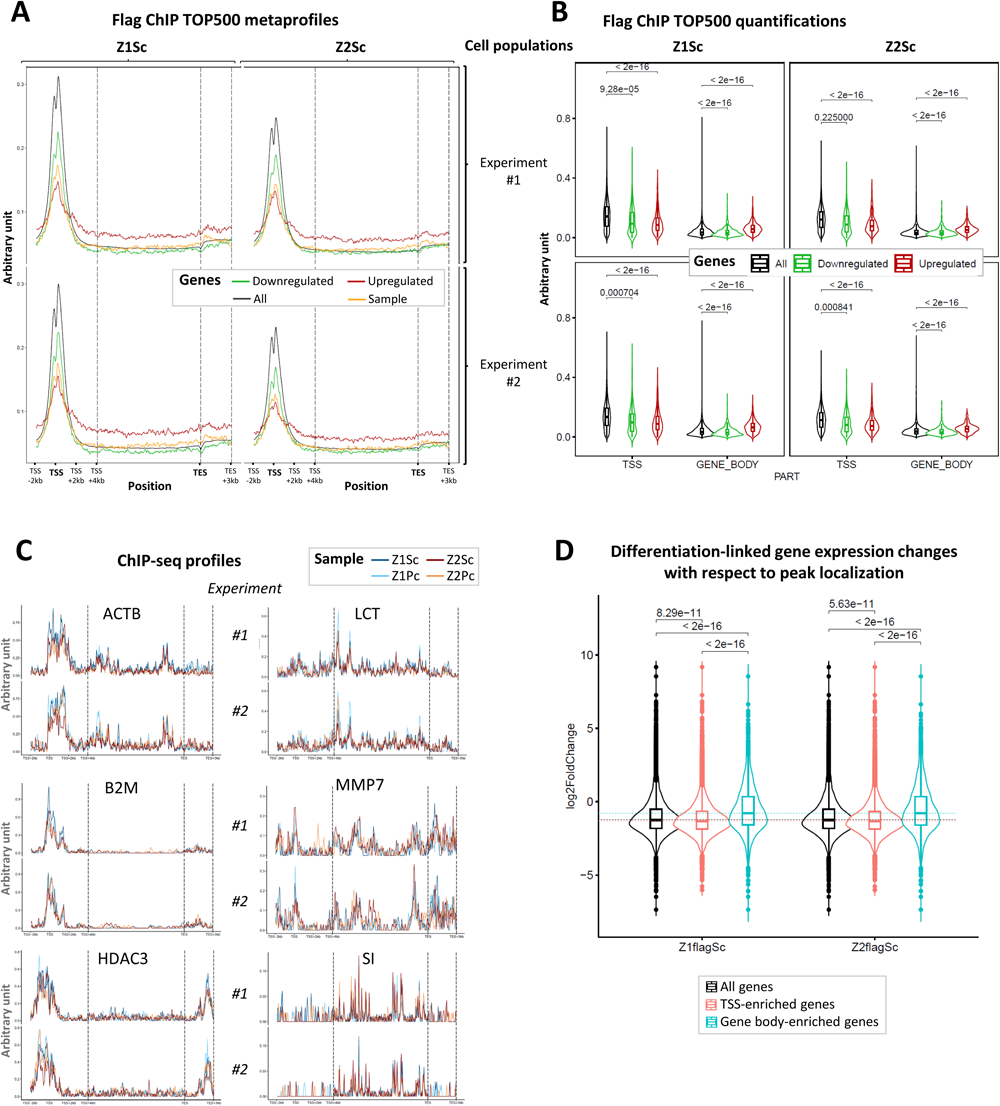
H2A.Z isoforms enrichment in gene bodies is higher on genes activated during differentiation. **A:** Metaprofiles of H2A.Z.1 or H2A.Z.2 ChIP-Seq signals on the most regulated genes during differentiation of subconfluent genome-edited Caco-2/15 cells. The 500 most activated genes during differentiation (TOP500 upregulated) and the 500 most repressed genes (TOP500 down-regulated) were identified from RNA-seq data obtained in both sub-confluent and 7-days post-confluent cells. Metaprofiles for these genes populations as well as for all genes (black) as a control are shown. **B:** The mean signal around TSS and on gene body was calculated for each individual gene. Violin plots showing these values for TOP500 upregulated, downregulated or all gene populations are drawn. Wilcoxon tests using BH correction were applied and p values are indicated on the graphs. **C:** Individual profiles of H2A.Z isoforms ChIP-Seq on the indicated housekeeping (left panels) and enterocyte-specific (right panels) genes. **D:** Impact of peaks abundance on TSS or Gene Body on gene expression during differentiation. Violin box plots represent the genome-wide mean for gene expression, in subconfluent flag-tagged cells, of All genes or genes enriched in peaks within their TSS or their Gene Body. Wilcoxon tests using BH correction were applied and results are indicated on the graphs.

Altogether, these data thus demonstrate that CRISPR-mediated genome editing of H2A.Z-encoding genes did not alter the differentiation abilities of the cells, as also shown in SuppFig.1A with accumulation of CDX2 intestine-specific transcription factor (Hryniuk *et al*., 2012), or Sucrase-Isomaltase (SI) and Lactase-Phlorizin Hydrolase (LPH/LCT) enterocyte marker (Van Beers *et al*., 1995) proteins.

We then performed in these two cell lines ChIP-seq experiments with Flag or total H2A.Z antibodies in order to characterize H2A.Z.1, H2A.Z.2 or total H2A.Z genomic occupancy from subconfluent (Sc) Caco-2/15 cells or from cells induced to differentiate for 72 hours after reaching the confluence (Pc) (see SuppFig.3A for PCA analysis). Thereafter, the cell populations expressing H2A.Z.1-flag are called “Z1Sc” or “Z1Pc” depending on the confluence stage (subconfluent or 7-days post-confluent stage, respectively), whereas that expressing H2A.Z.2-flag are named “Z2Sc” or “Z2Pc”.

We observed that, both in sub-confluent or post-confluent Caco-2/15 cells, H2A.Z.1 and H2A.Z.2 are enriched around gene TSS, in a manner very similar to H2A.Z (Fig.1B). Moreover, metadata analyses at genes show that, as H2A.Z, both H2A.Z isoforms mostly localize around TSS with a double peak, reflecting H2A.Z incorporation at the −1 and + 1 nucleosomes (Figure 1B upper panels). Taken together, these data indicate that epitope tagging of H2A.Z isoforms does not affect their genomic localization.

We next investigated whether genes with different expression patterns could have specific H2A.Z occupancy with respect to H2A.Z isoforms. We first ranked genes according to their expression levels in eight different classes, and plotted for each class the mean profile of H2A.Z isoforms occupancy on genes (Fig.1C). As already shown, total H2A.Z occupancy at TSS largely correlates with transcription (Fig.1C, lower panels). A very similar result was observed for both H2A.Z.1 and H2A.Z.2 (Flag IPs, upper panels), suggesting that there is no global specificity of H2A.Z isoforms binding with respect to gene expression or cell differentiation status.

We also ranked genes with respect to their variations upon differentiation, selecting the Top 500 most activated and most repressed genes, and plotted the mean profile on genes. Using ChIP-seq data from subconfluent cells (Fig.2A), we first observed very similar profiles for H2A.Z.1 and H2A.Z.2. Moreover, genes which are highly regulated, irrespective of whether they are induced or repressed, have less H2A.Z isoforms around their TSS compared to all genes (Fig.2A). Indeed, box plots showing H2A.Z isoforms levels at the TSS of these population show that the difference is significant (Fig.2B). Interestingly, such a low enrichment of H2A.Z at the TSS of regulated genes was already observed in plants (Coleman-Derr and Zilberman, 2012), suggesting that it is a conserved feature of regulated genes. However, and very strikingly, genes which are highly activated upon differentiation have a higher H2A.Z isoforms occupancy on their bodies compared to all genes or repressed genes, as shown on the mean profiles (Fig.2A) and on box plots showing H2A.Z levels in the bodies of these gene populations (Fig.2B). Note that we also observed the same differential enrichment for these gene sets in post-confluent samples (SuppFig.3B). Moreover, randomly chosen genes with similar expression level and gene size than the Top500 upregulated one, harbor a metaprofile similar to all genes (see “Sample” profile in Fig.2A), providing evidence that gene body enrichment observed in the Top500 upregulated population is not due to differences in the absolute expression level. Thus, we conclude from this analysis that genes which are induced during enterocyte differentiation have a higher H2A.Z occupancy at their bodies prior to differentiation induction, independently of their expression level. To our knowledge, this feature has never been described so far.

**Figure 3:**
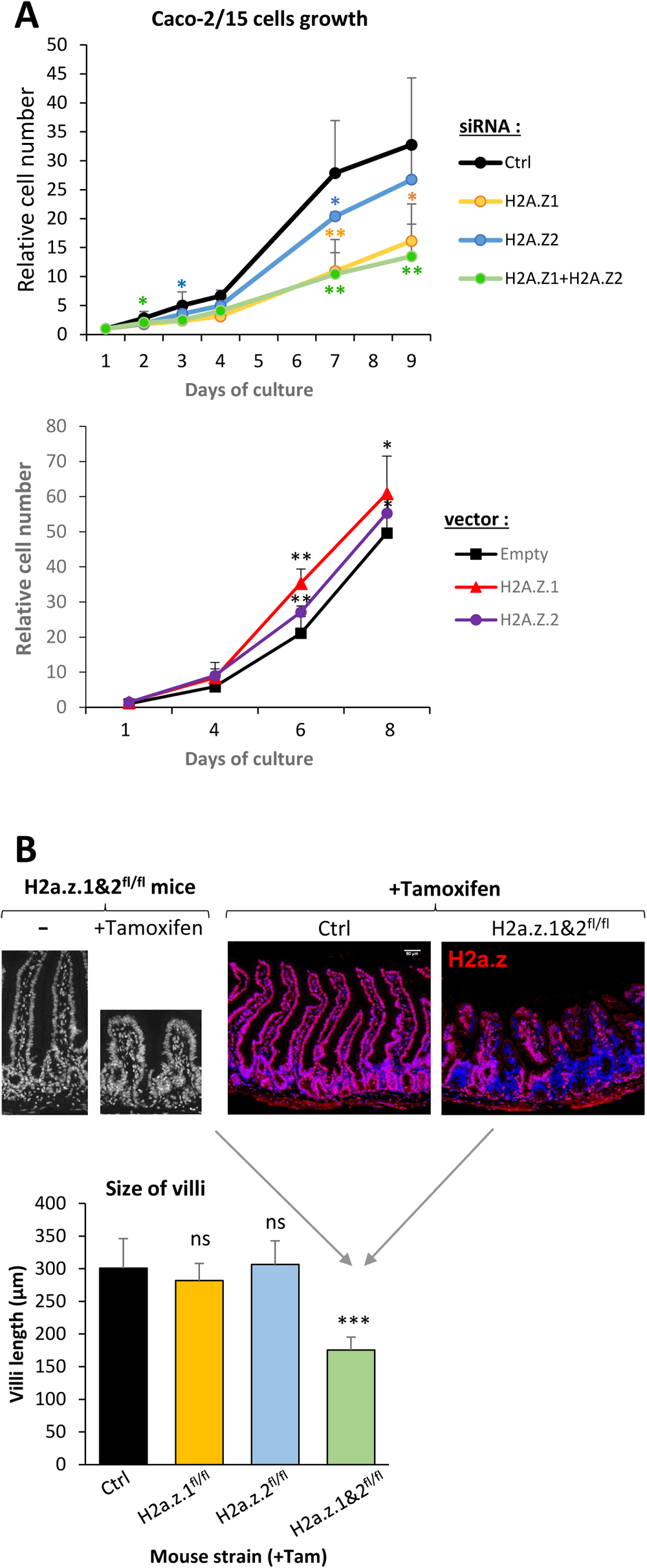
Essential redundant role of H2A.Z isoforms in intestinal epithelial renewal. **A**: Cell growth assay were performed using Caco-2/15 cells transfected with indicated siRNA (upper panel) or expression vector (lower panel). Cell number was assessed at different times after transfection as indicated, and the mean and standard deviation from 5 independent experiments are represented. Statistical analysis was done using the Student’s t-Test (*<0.05; **<0.02). **B**: Representative H2A.Z immunohistofluorescence (right panels, in red) of jejunal sections in control or double mutant *H2a.z.1&2^fl/fl^* mice, ten days after the recombination induced by tamoxifen treatment. Nuclei are stained with DAPI in blue. Scale bar represent 50µm. Left panels are from *H2a.z.1&2^fl/fl^*mice with or without tamoxifen treatment. Nuclei are stained with DAPI (in grey). The graph represents measurements of the size of about 100 villi for each mouse. Genotypes of mice are stated below the diagram. Statistical analysis was done using the Student’s t-Test (***<0.01).

Altogether, these data suggest that H2A.Z.1 and H2A.Z.2 do not have isoform-specific incorporation global pattern with respect to gene expression or gene regulation during differentiation. Indeed, we found that the amounts of H2A.Z.1 and of H2A.Z.2 are highly correlated both at gene TSS and on gene bodies (see Figures 1&2 and SuppFig.3B). They also indicate that the presence of H2A.Z isoforms in gene bodies in subconfluent cells is linked to gene activation during differentiation.

### H2A.Z isoforms peaks are enriched in the bodies of activated genes

Interestingly, we noticed that profiles of H2A.Z isoforms occupancy in subconfluent cells on individual genes show striking differences between examples of enterocytes specific genes and house-keeping genes (Fig.2C): whereas the latter genes (left profiles) harbor classical profiles with peaks of H2A.Z isoforms occupancy around their TSS, the former (right profiles) do not show strong peaks at TSS but harbor many reproducible peaks in their gene bodies.

We thus performed a peak calling analysis to investigate the localization of H2A.Z isoforms at individual genes level. Indeed, the size of gene bodies may have masked localized specific binding of one or the other isoform. Moreover, Figure 2C show that enterocyte specific genes harbor peaks within their gene bodies. Peak calling analysis was performed on H2A.Z.1 and H2A.Z.2 ChIP-seq with high stringency parameters to minimize false peaks detection, and peaks were considered only when they overlapped between the two replicates. Taking into account all conditions (irrespective of the H2A.Z isoform and of the cell confluence state), we found 169 284 peaks, with 24.8% around TSS, 24.6% at gene bodies, 20% overlapping both a TSS and a gene body and 30.6% in intergenic regions (SuppFig.4A).

**Figure 4:**
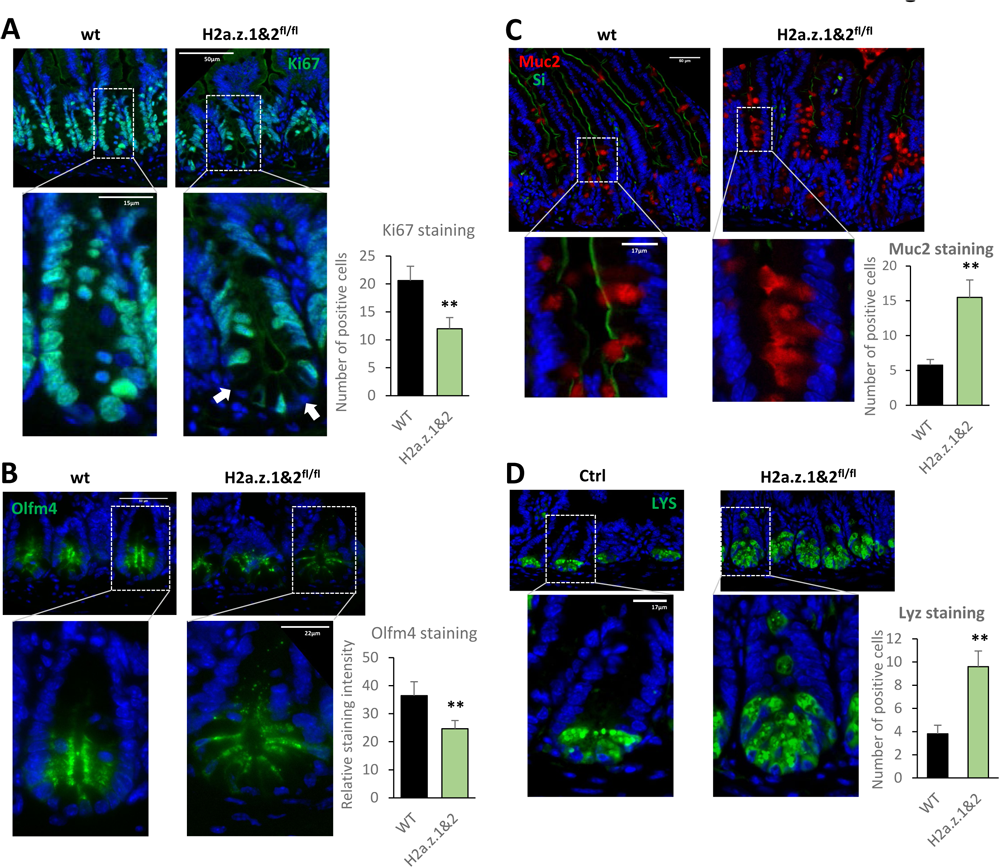
H2A.Z isoforms are essential for proliferation and stemness. **A**: Representative Ki67 immunohistofluorescence on jejunal sections of control or *H2a.z.1&2^fl/fl^* mice, 10 days after starting tamoxifen treatment. Nuclei are stained with DAPI in blue. Arrows indicate Ki67 negative cells in the crypt. Quantification of positive cell number was performed on at least 20 crypt-villus axis per genotype. Statistical analysis was done using the Student’s t-Test (**<0.02). **B**: Same as in A for Olfm4 staining. Quantification was done by measuring global signal intensity from at least 20 crypt-villus axis per genotype. **C**: Same as in A for Muc2 (in red) and Sucrase-isomaltase (in green) staining. **D**: Same as in A for Lys immunostaining.

If we now considered peaks from a genes point of view, 8902 genes (59.9%) have a peak only at their TSS, 2161 (14.5%) only in their body, and 3808 (25.6%) in both (SuppFig.4B). To gain insights into the function and dynamics of these peaks, we computed for each gene and each condition the number of reads present in TSS-associated peaks or gene body-associated peaks.

Interestingly, a significant population of genes (2516 genes for H2A.Z.1 and 2542 genes for H2A.Z.2) have more signal in peaks in their body than at their TSS in subconfluent cells (SuppTables 1 and 2, respectively), contrary to the general view that H2A.Z is generally enriched around TSS. Importantly, the Log2 of fold change during differentiation of these two populations is higher than that of all genes or that of genes which have more H2A.Z isoforms at their TSS than in the gene body (Fig.2D). Thus, these data suggest that H2A.Z isoforms peaks within gene bodies may play a role in gene regulation during differentiation. And indeed, the gene ontology analysis of these two subsets revealed a clear link of such genes with signal transduction, adhesion/polarization and ion transport processes (SuppFig.4C). In addition, the presence of these peaks is probably responsible for the higher mean of H2A.Z isoforms occupancy we observed on the bodies of highly activated genes (Fig.2A).

This result thus suggests that some genes important for intestinal epithelial biology are specifically labelled by H2A.Z isoforms peaks in their bodies, which can participate in processes linked to epithelial differentiation.

Thus, altogether, these findings indicate that the presence and dynamics of H2A.Z isoforms peaks at gene bodies correlates with lower repression or with activation of genes during differentiation process, pointing to the importance of deciphering the function of H2A.Z isoforms in intestine homeostasis.

### H2A.Z isoforms are essential for the intestinal epithelium maintenance in a redundant manner

We thus intended to investigate the function of H2A.Z isoforms in intestine cells proliferation abilities. We previously showed that H2A.Z.1-deficient cells have proliferation defects, in cultured conditions as well as *in vivo* (Rispal *et al*., 2019). This suggests that H2A.Z.2 isoform cannot compensate for H2A.Z.1 deficiency.

We analyzed the impact of the siRNA-mediated decrease of H2A.Z.1 and/or H2A.Z.2 expression on proliferation abilities of Caco-2/15 cells (Fig.3A, upper panel). We observed that the downregulation of each isoform (see SuppFig.6A for siRNA efficiency) impairs the proliferation of this cell model. Interestingly, the decrease of proliferation abilities is more pronounced upon H2A.Z.1 knockdown, without any additional effect of the concomitant reduction of H2A.Z.2. This result correlates with the impact of siRNA-mediated downregulation of H2A.Z1 and H2A.Z2 on the pool of total H2A.Z protein, where H2A.Z.1-targetting siRNA reduces by about 2/3 of the total H2A.Z signal, whereas H2A.Z.2-tagetting one has a limited effect (around 1/3 of reduction) (see H2A.Z panel in SuppFig.1A), probably due to the relative abundance of each isoform in Caco-2/15 cells. Accordingly, transfecting expression vectors encoding for one isoform or the other induced a similar proliferative advantage in cell population (Figure 3A, lower panel. See SuppFig.6B for the quantification of the overexpression). Thus, these results suggested that the two isoforms exert similar roles on cell growth, proportionally to their respective expression levels.

Zhao and collaborators showed that the intestinal epithelial structure is highly affected (i.e. decreased size of villi) in whole epithelium double knock-out mice (*Villin-Cre; H2afz^fl/fl^/H2afv^fl/fl^*), whereas individual deficiencies for one isoform did not induce any abnormality in the size of the structures (Zhao *et al*., 2019). Thus, these results suggest a redundant role *in vivo* of H2A.Z isoforms on intestinal epithelial renewal and maintenance. Thus, to precisely characterize the contribution of each H2A.Z isoform in the renewing of the intestinal epithelium, we used mice models allowing the knock-out for one or both isoforms in an inducible manner, specifically in the intestinal epithelial stem cells (*Lgr5-Cre^ER^*^T2^ *; H2afz^fl/fl^ and/or H2afv^fl/fl^*). This model allowed us to avoid any bystander effect potentially caused by the knock-out of H2AZ in the villus. Ten days after tamoxifen induction, and thus genomic recombination (SuppFig.7A), we observed a strong decrease of the villus length in the double KO mice (Fig.3B). We did not observe any major effect on the structure size upon the knock-out of only H2A.Z.1 ((Rispal *et al*., 2019) and quantifications in Fig.3B) or H2A.Z.2 (Fig.3B). Note that discriminating the specific expression of each isoform is difficult due to the lack of isoform-specific antibodies. In immunohistofluorescence experiments using H2A.Z antibodies, we observed a mosaic pattern of the H2A.Z staining upon H2A.Z.1/2 knockdown (Fig.3B), which is the result of the crypt-villus structures colonization by cells originating from both recombined and wild-type stem cells (typical for such models). We also observed a reduction in H2A.Z staining upon specific knockdown of H2A.Z.1 (SuppFig.7B), but not in H2A.Z.2 knock-down samples, probably due to the differences in the relative abundance of each isoform. These results, in agreement with Zhao *et al*.‘s work, confirm that the H2A.Z isoforms have redundant and essential roles on the abilities of stem cells to ensure tissue maintenance.

To test the impact of isoforms on crypt cells proliferation, we isolated these cells from mice and generated *in vitro* 3D culture of intestinal organoids. This system allowed us to induce or not H2A.Z isoforms knock-out in stem cells from the same animal. We observed that the induction of the genomic recombination using Tat-CRE recombinant protein resulted in a significant decrease in mRNA expression for both H2A.Z isoforms in organoids derived from mice harboring recombination on H2A.Z.1-and H2A.Z.2-encoding genes. Importantly, the growth of the 3D structures in these organoids was also decreased (SuppFig.8A), with an increase in the number of organoids whose growth was less than 2-fold six days after the induction (i.e. 13.3% in Ctrl versus 30.8% in double KO, in green) (SuppFig.8B, lower panel). As *in vivo*, the single KO of H2A.Z.1 had no effect (SuppFig.8B, upper panel). These data confirmed the redundant compensatory effects of both isoforms in the development of complex 3D cultured structures emerging from crypt cells.

Next, we intended to investigate whether this phenotype is due to proliferation defects of the progenitors or to a global decrease of the stemness. First, by analyzing Ki67 staining, we observed a proliferation defect upon the induction of the double KO, in particular at the base of the crypts (Fig.4A) where the number of negative cells appears to be increased. H2A.Z.1 KO alone had no major effect (Rispal *et al*., 2019), in accordance with the lack of effect of this individual KO on the epithelial structure. The same result was observed in *in vitro* models since the double KO in organoids (SuppFig.8C for mRNA expression levels) also leads to the decrease of Ki67 staining (SuppFig.8D). Thus, these results indicate that only the knock-out of the two isoforms together impacts intestinal cell proliferation.

We next analyzed the role of H2A.Z isoforms in stemness maintenance by staining for the stem cell marker Olfm4. We observed a decrease of Olfm4 staining in double KO mice compared to control (Fig.4B). In organoids, the double KO also led to the decrease of the expression of the stem cells markers Lgr5 and Olfm4 mRNA (SuppFig.8E, lower panels), whereas the single deletion of H2A.Z.1 had a much weaker effect (upper panels).

Altogether, these results uncover the major and redundant role of both isoforms of H2A.Z in the renewal of the intestinal epithelium by contributing to normal proliferation and stemness maintenance.

### H2A.Z isoforms act redundantly to control lineage specification

We next analyzed the role of H2A.Z isoforms on lineage differentiation. We observed a huge increase of Muc2^+^ cell number (marker of goblet cells) in double KO mice (Fig.4C), as compared to control or single KOs (SuppFig.9A and (Rispal *et al*., 2019)). Moreover, the number of Paneth cells (Lys^+^) also increases in the double KO mice compared to control (Fig.4D), whereas there was no difference in H2A.Z.1 single KO (Rispal *et al*., 2019). Altogether, these results suggest that both isoforms of H2A.Z are involved, in a redundant manner, in the inhibition of secretory lineage commitment. Note however that we did not observe any major effect of H2A.Z KO on the number of enteroendocrine cells (SuppFig.9B). This result can be explained by a specific enteroendocrine lineage control or by the intestinal region we analyzed (jejunum). Indeed, Kim and Shivdasani have shown that there is a specific regulation of these cells depending of the intestinal region (duodenum vs ileum) (Kim and Shivdasani, 2011).

Altogether, these results suggest a dose-dependent effect of total H2A.Z which determines the commitment of crypt cells towards the secretory fate.

### Antagonistic control of enterocytes differentiation by H2A.Z.1 and H2A.Z.2

In our previous work, we showed the essential role of H2A.Z.1 in intestinal homeostasis by preventing enterocyte terminal differentiation, through the inhibition of binding to DNA of the CDX2 transcription factor (Rispal *et al*., 2019). Thus, we intended to analyze the function of H2A.Z.2. As already shown (Rispal *et al*., 2019), the induction of H2A.Z.1 KO in mice led to a major increase of Sucrase-Isomaltase (SI) (a marker of enterocyte terminal differentiation) staining compared to control in immunohistofluorescence experiments on intestinal tissues (Fig.5A). The deletion of H2A.Z.2-encoding alleles did not induce any alteration of SI staining as compared to control mice. However, the induction of double KO in mice led to the complete disappearance of Sucrase-Isomaltase staining (Fig.5A and also 4C). Therefore, HA.Z.2 KO reversed the effect of the H2A.Z.1 KO with respect to enterocyte differentiation, indicating an antagonistic function of H2A.Z isoforms on this process. Moreover, considering the above-described results on secretory cells differentiation, these data suggest that the double depletion of H2A.Z.1 and H2A.Z.2 leads to a differentiation shift from enterocytes to secretory cells, suggesting that H2A.Z isoforms redundantly control the choice of intestinal lineages.

**Figure 5:**
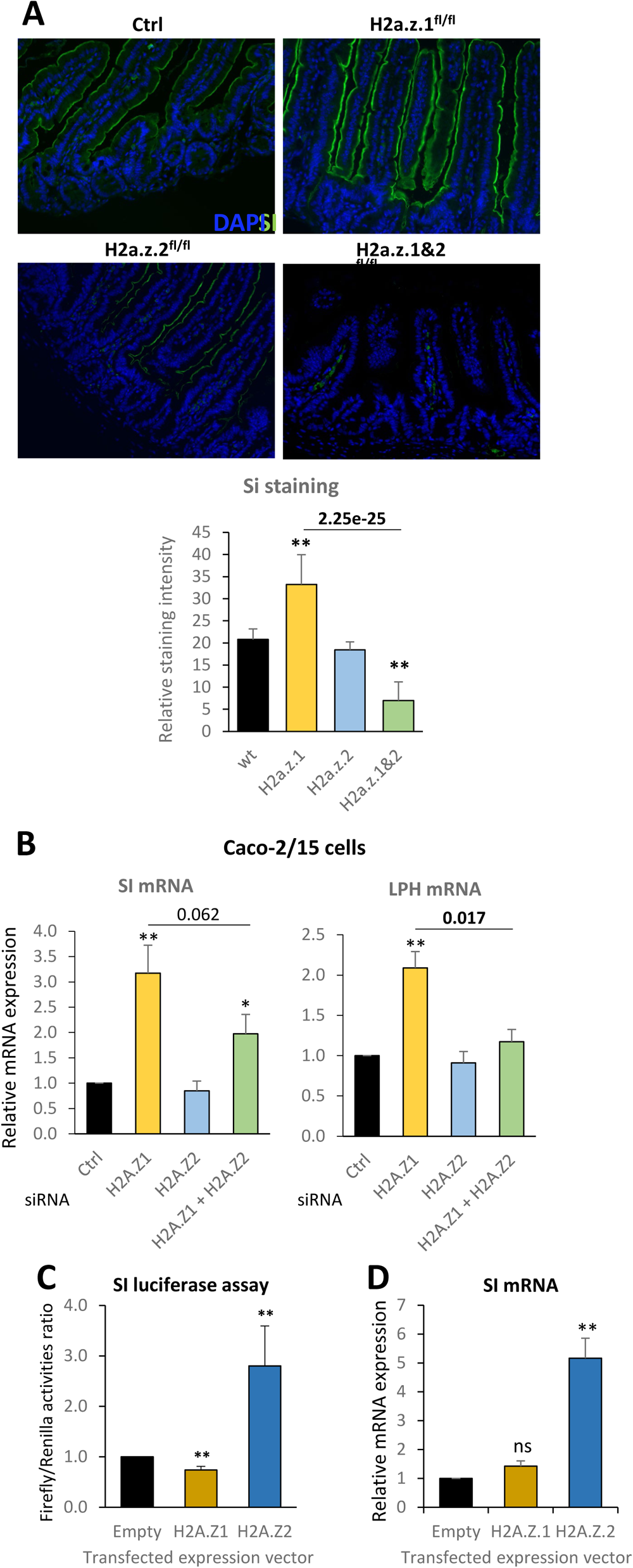
Antagonistic control of enterocytes differentiation by H2A.Z.1 and H2A.Z.2. **A**: Representative Sucrase-Isomaltase (SI) immunohistofluorescence of jejunal sections in mice from the indicated genotypes, 10 days after starting tamoxifen treatment. Nuclei are stained with DAPI in blue. Quantification was done by measuring global signal intensity from at least 20 crypt-villus axis per genotype. Statistical analysis was done using Student’s t-Test relative to wild-type (wt) condition and indicated in the graphic (**: p-value <0.02). The one-way Student’s t-Test p-values for double KO tissues compared to H2A.Z.1-deficient ones are also indicated in the graph. **B**: mRNA expression of SI and Lactase (LPH) in Caco-2/15 cells, 3 days after the transfection of siRNA directed against H2A.Z.1, H2A.Z.2 or both. The mean and standard deviation from 5 independent experiments is represented. Statistical analysis was done using Student’s t-Test relative to control siRNA (Ctrl) condition and indicated in the graphic (p-value: * <0.05; ** <0.02). The one-way Student’s t-Test p-values for double siRNA-transfected cells compared to H2A.Z.1-tagetting one are also indicated in the graph. **C**: Firefly luciferase assays 2 days after transfection of H2A.Z.1 or H2A.Z.2 -expressing vectors. Results were calculated relative to measurements of Renilla luciferase constitutively expressed from a normalizing co-transfected vector. The mean and standard deviation from 3 independent experiments is represented. Statistical analysis was done using Student’s t-Test and the p-values relative to empty vector condition are indicated in the graphic (**<0.02). **D**: Endogenous SI mRNA expression 7 days after transfection of H2A.Z.1 or H2A.Z.2 -expressing vector. Total mRNA was extracted, reverse-transcribed and cDNA were analyzed by RT-qPCR. The mean and standard deviation from 4 independent experiments is represented. Statistical analysis was done using Student’s t-Test and the p-values relative to empty vector condition are indicated in the graphic (**<0.02).

To investigate further the respective functions of H2A.Z isoforms in enterocyte differentiation, we made use of the well characterized colonic Caco-2/15 epithelial cells, which express markers of enterocyte differentiation when they reach confluence. We found that, as in mice, the knock-down (KD) of H2A.Z.1 using specific siRNAs led to an increase of the enterocyte markers SI or LPH (Lactase Phlorizin Hydrolase) mRNA, as already shown (Rispal *et al*., 2019), whereas the KD of H2A.Z.2 had no effect (Fig.5B). Strikingly, the induction observed upon H2A.Z.1 knock-down was partially reversed by concomitant depletion of H2A.Z.2. This shows an antagonistic role of both isoforms on markers expression, the effect of H2A.Z.1 depletion being, at least in part, dependent on H2A.Z.2, therefore mimicking what we observed in vivo. We next tested the effect of H2A.Z isoforms overexpression. Using a reporter vector in which the luciferase gene is controlled by the SI promoter, we found that overexpression of H2A.Z.2 led to an increase of SI promoter activity (Fig.5C). Moreover, it induced a long-term increase in endogenous SI expression (Fig.5D) (see SuppFig.6B for quantification of vector-mediated overexpression). Strikingly, overexpression of H2A.Z.1 had no major effect, confirming that the two H2A.Z isoforms play specific functions in the activation of enterocyte markers expression in Caco2 cells.

These last data suggest that H2A.Z.2 favors enterocyte differentiation in Caco-2/15 cells. To test whether its chromatin incorporation is important for this function, we intended to work on H2A.Z incorporators, namely SRCAP and p400 complexes. Interestingly, in a previous study (Lamaa *et al*., 2020), we showed that p400 binds more efficiently to H2A.Z.2, suggesting that it could preferentially incorporate H2A.Z.2.

We first confirmed the preference of p400 for binding to H2A.Z.2 in Caco2-15 cells by co-immunoprecipitation experiments (SuppFig.10). We next analyzed the incorporation of H2A.Z.1 and H2A.Z.2, after the depletion of SRCAP or p400 in Caco-2/15 tagged for the respective H2A.Z isoforms, by ChIP experiments (Fig.6A). We observed that depletion of H2A.Z incorporator SRCAP induced a decrease of both H2A.Z.1 and H2A.Z.2 enrichment at many promoters linked to intestine homeostasis. In contrast, p400 siRNA led to the specific reduction of the incorporation of H2A.Z.2 whereas H2A.Z.1 enrichment remained almost unaffected, suggesting a preferred incorporation of H2A.Z.2 by p400 complex due to a higher interaction ability.

**Figure 6:**
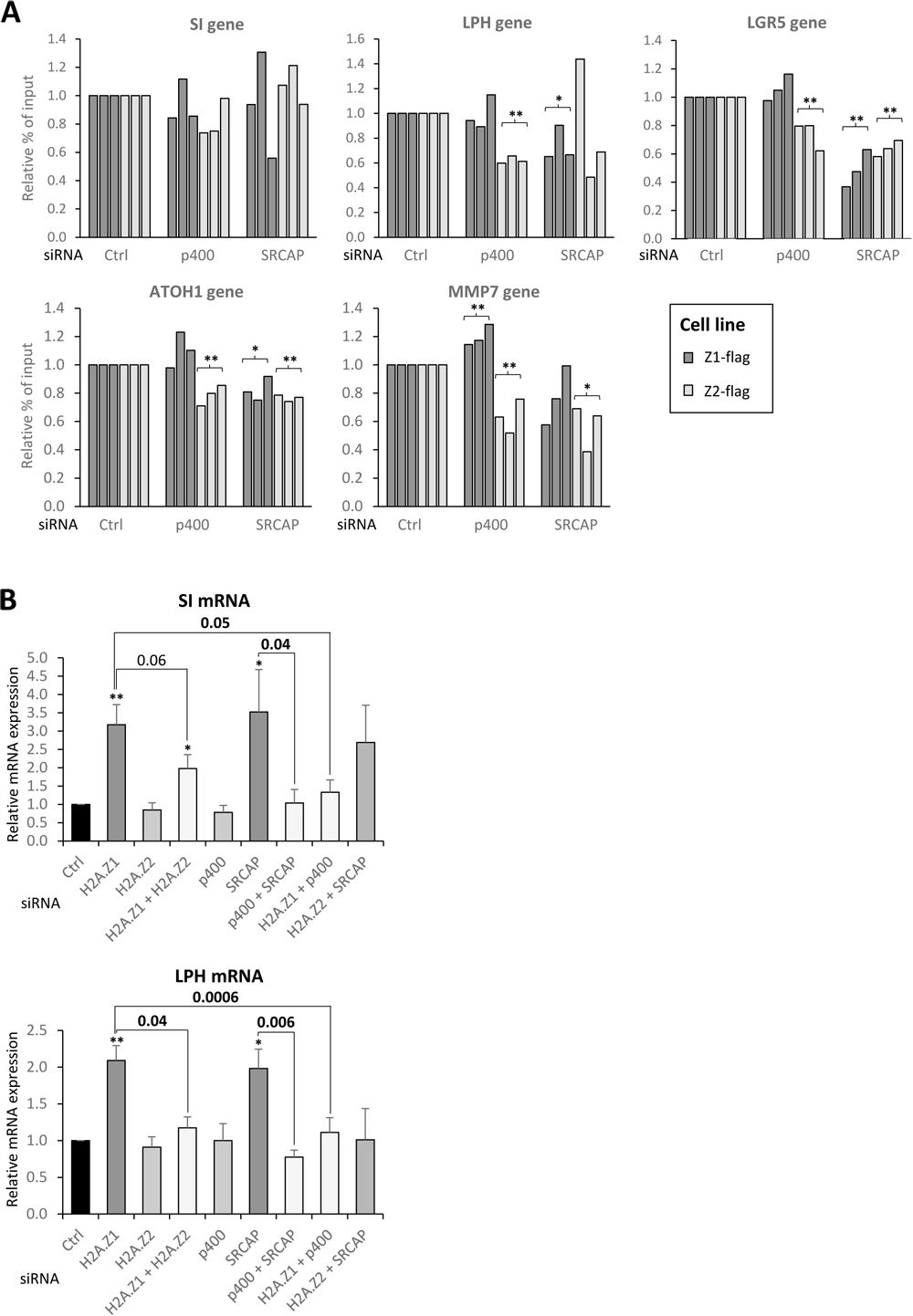
Modulating H2A.Z-related enterocyte differentiation by SRCAP and p400. **A:** Incorporation of H2A.Z isoforms upon siRNA-mediated knockdown of p400 or SRCAP. Caco-2/15 cells genetically edited to express Flag-tagged H2A.Z isoforms were transfected with the indicated siRNA. 72 hours later, chromatin immunoprecipitation was performed using anti-Flag antibody to assess enrichment of H2A.Z.1-Flag or H2A.Z.2-Flag. The amount of indicated promoters were then analyzed by qPCR and calculated relative to 1 for the control siRNA. Three independent experiments are shown. Statistical analysis was done using Student’s t-Test and the p-values relative to control siRNA condition are indicated in the graphic (*<0.05; **<0.02). **B**: SI and LPH mRNA expression in Caco-2/15 cells transfected with the various combinations of siRNAs, as indicated below the graphs. The mean and standard deviation from 5 independent experiments is represented. Statistical analysis was done using the Student’s t-Test. p-values are indicated relative to control siRNA condition (*<0.05; **<0.02) or between other relevant conditions (numeric values).

We next tested the effects of SRCAP and p400 depletion on enterocyte markers. Interestingly, by using siRNA (see SuppFig.6A for efficiencies), we observed that the depletion of SRCAP in Caco-2/15 cells led to an important increase of SI and LPH expression, as observed for H2A.Z.1 depletion (Fig.6B), whereas p400 depletion had no effect. However, p400 depletion reversed the induction of SI and LPH expression observed upon SRCAP or H2A.Z.1 depletion, in a manner similar to the H2A.Z.2 depletion-linked reversion of the H2A.Z.1 effects. These results indicate that p400 depletion phenocopies H2A.Z.2 depletion, and is consistent with a function of H2A.Z.2 in enterocyte differentiation intimately linked to its incorporation in chromatin.

Altogether, these data confirm the specific role of H2A.Z isoforms in enterocyte differentiation and suggest that their ratio is a key regulator for the expression of enterocyte differentiation markers.

### The H2A.Z.2/H2A.Z.1 ratio increases during differentiation

Careful examination of Supplemental Figure 1, showing H2A.Z isoforms expression using the cell lines we generated, suggested that tagged H2A.Z.2 expression slightly increased when cells reach confluence, whereas tagged H2A.Z.1 expression decreased, if anything (SuppFig.1B), which could result in changes of the H2A.Z.1/H2A.Z.2 ratio. Given the specificity of H2A.Z.2 in activating enterocyte markers expression as shown above, this change in ratio could be important for enterocyte differentiation.

Since this experiment compared two different cell lines, we intended to monitor the evolution of H2A.Z isoforms expression when parental Caco-2/15 cells reach confluence. Since no antibody allowing to discriminate between H2A.Z isoforms is available, we performed RT-qPCR experiments. We found that, in agreement with protein expression in tagged cells, the H2A.Z.1-encoding mRNA levels remained more or less stable at confluence whereas H2A.Z.2-encoding mRNA levels increased (Fig.7A). As a consequence, the ratio between H2A.Z isoforms increases in favor of H2A.Z.2.

**Figure 7:**
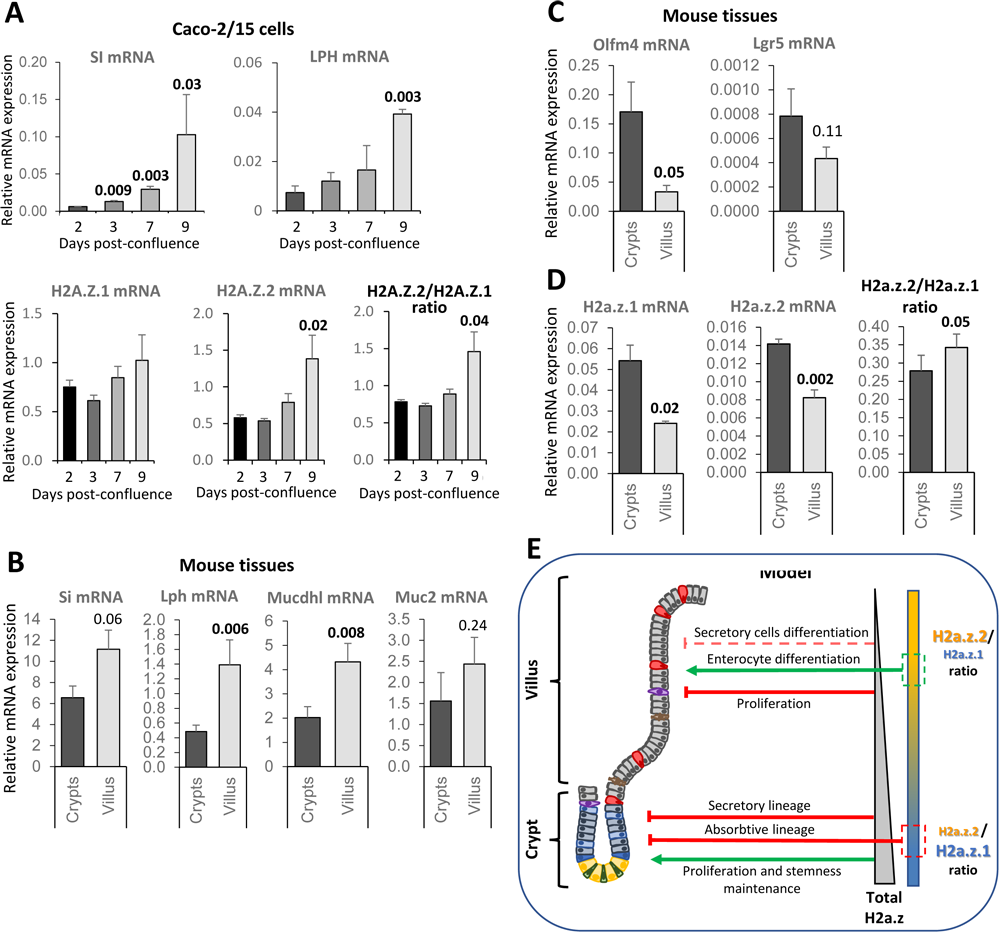
The ratio between H2A.Z isoforms correlates with intestinal differentiation. **A:** Caco-2/15 were induced to confluence and harvested at the indicated time. Enterocyte-differentiation markers and H2A.Z isoforms mRNAs were quantified. The H2A.Z.2/H2A.Z.1 mRNAs ratio is also shown. The mean and standard deviation from 3 independent experiments is shown. Statistical analysis was done using the Student’s t-Test *vs* Day 2 and p-values are indicated when below 0.1. **B:** Expression of differentiation markers in isolated crypts or villi of wild-type mice. After epithelium fractionation as described in the Methods section, mRNA was extracted and expression of differentiation markers was analyzed by RT-qPCR. The mean and standard deviation from 4 mice is shown. Statistical analysis was done using the paired Student’s t-Test *vs* Crypts. **C:** As in B, except that the expression of crypt/stem cell markers was analyzed. **D:** As in B, except that the expression of H2.A.Z isoforms was assessed. The H2A.Z.2/H2A.Z.1 mRNAs ratio is also shown. **E:** Model of control of intestinal epithelial homeostasis by H2A.Z isoform dynamic. In the crypt, the high quantity of total H2A.Z is essential for maintaining the undifferentiated state of cells by facilitating proliferation and maintaining stemness, and by inhibiting the differentiation. During the migration of cells along the villus, the decrease of H2A.Z coupled to the increase of H2A.Z.2/H2A.Z.1 ratio allows differentiation and favors the absorptive lineage.

To test whether this is also true *in vivo*, we set-up the dissociation of the mouse intestinal epithelium into crypt or villus -enriched fractions. We confirm the enrichment by RT-qPCR of differentiated (Fig.7B) and stem (Fig.7C) cells markers in the villi and the crypts samples, respectively. We observed, as expected (Kazakevych *et al*., 2017), a global reduction of H2A.Z expression in the differentiated compartment enriched in villi, since both H2A.Z.1-and H2A.Z.2-encoding mRNA expression decreased (Fig.7D). However, the H2A.Z.2/H2A.Z.1 ratio increased correlatively with the differentiation state of the fractions.

Thus, altogether, these data obtained in Caco-2/15 cells and in mice are consistent with an increase of the H2A.Z.2/H2A.Z.1 ratio upon enterocyte differentiation, at least at the level of the availability of their mRNA. Considering the respective role of H2A.Z.1 and H2A.Z.2 on enterocyte differentiation we observed in Figure 5, our results suggest that this change of ratio could participate in the expression of enterocyte-specific genes during differentiation (see our model in Figure 7E).

## Discussion

In this study, we highlighted the respective contributions of two isoforms of the same histone variant in a crucial homeostatic process: the permanent intestinal epithelium renewing. We showed that the functions of the two isoforms are redundant for stem cell proliferation and tissue maintenance, as already shown (Zhao *et al*., 2019), as well as for the control of commitment towards epithelial secretory cell lineage. The molecular basis for this redundancy probably relies on the strong correlation we observed for the incorporation of H2A.Z.1 and of H2A.Z.2 at TSS and in gene bodies. The function of these two isoforms when incorporated is probably similar on the expression of genes important for these processes. The redundancy between H2A.Z.1 and H2A.Z.2 could participate in the robustness of the mechanisms important for intestine homeostasis.

In contrast, we observed that the two isoforms have an opposite function for enterocyte differentiation, with the activating effect of H2A.Z.1 depletion being dependent on H2A.Z.2. Our data thus suggest that H2A.Z.1 functions as a repressor of enterocyte differentiation, as we already showed (Rispal *et al*., 2019) whereas H2A.Z.2 favors it. Importantly, distinct functions between H2A.Z.1 and H2A.Z.2 have already been described in other differentiation pathways, such as neuronal differentiation (Shen *et al*., 2018) or craniofacial development (Greenberg *et al*., 2019). However, to our knowledge, our data are the first to demonstrate an opposite function of the two isoforms on a differentiation process. They underline the importance of understanding how the expression of H2A.Z.1 and H2A.Z.2 are regulated with respect to intestine homeostasis. We previously showed that the promoter of the H2A.Z.1-encoding gene is a target of the WNT signaling pathway, leading to its decrease in expression during differentiation (Rispal *et al*., 2019). However, how the H2A.Z.2-encoding genes is regulated by signaling pathways linked to intestine homeostasis is still unknown.

What could be the mechanism of these opposite roles? First, H2A.Z1 and H2A.Z.2 could be incorporated at different genomic locations, regulating different set of genes. Our data obtained upon depletion of H2A.Z incorporators are in agreement with a different localization of H2A.Z.1 and H2A.Z.2 being important for enterocyte differentiation: indeed, the depletion of p400, which induces a reduced incorporation of H2A.Z.2, mimics the effect of H2A.Z.2 depletion, whereas the depletion of SRCAP mimics the depletion of H2A.Z.1. It could thus be envisioned that p400 and SRCAP could be recruited at different genomic locations, such as at genes important for enterocyte differentiation. This would lead to different H2A.Z.1/H2A.Z.2 ratio from one gene to the other and different transcriptional response of these genes. By peak calling analysis, we found a significant number of promoters or gene bodies specifically harboring peaks of H2A.Z.1 (SuppFig.5), which could thus represent genes specifically regulated by H2A.Z.1, further supporting this hypothesis. We also found few genes with specific peaks of H2A.Z.2 in their promoters or gene bodies.

However, despite these genes with specific peaks, we observed that the amounts of H2A.Z.1 and H2A.Z.2 at TSS and gene bodies are highly correlated. This suggests an alternate hypothesis that would be that H2A.Z isoforms play opposite role when incorporated on some promoters important for enterocyte differentiation, with H2A.Z.2 activating them and H2A.Z.1 repressing them. Given that the two H2A.Z isoforms can replace each other in chromatin when depleted (Lamaa *et al*., 2020), isoform replacement would explain why the activation of enterocyte specific genes observed upon H2A.Z.1 knock-down is H2A.Z.2-dependent. Moreover, we already observed such an opposite function of H2A.Z.1 and H2A.Z.2 relying on the recruitment of specific effectors with opposite function with respect to gene expression (Lamaa *et al*., 2020). Such a mechanism could operate here for genes important for enterocyte differentiation. In this case, the global change in the H2A.Z.1/H2A.Z.2 ratio we observed during differentiation would result in change in the expression of these genes. Interestingly, p400 depletion mimics H2A.Z.2 depletion, whereas SRCAP depletion mimics H2A.Z.1 depletion. This raises the interesting hypothesis that these factors, besides being H2A.Z incorporators, could also be specific effectors of H2A.Z isoforms. Along this line, p400 is present in a complex which contains the histone acetyl transferase Tip60. The presence of H2A.Z.2 in chromatin could perhaps favor the local recruitment of Tip60, leading to histone acetylation and gene activation.

Finally, our genome-wide analyses uncover the unexpected link between H2A.Z isoforms occupancy at gene bodies and the propensity of genes to be induced in enterocyte differentiation. Most importantly, this is not related to differences in expression level since a similar difference is observed when compared with genes with similar expression levels. In plants, it was previously shown that genes annotated as “highly variable” also harbor high H2A.Z occupancy in their bodies compared to house-keeping genes, although in that case it was proposed that it was due to differences in gene expression levels. This was linked to an inverse correlation on gene bodies between DNA methylation and H2A.Z presence (Coleman-Derr and Zilberman, 2012). Despite this link between H2A.Z incorporation and DNA methylation was also shown in mammalian cells (Conerly *et al*., 2010), that H2A.Z presence at gene bodies confers a mark dictating the behavior of genes has never been observed in mammals to our knowledge. Furthermore, in the case of enterocyte differentiation, it specifically marks genes that are going to be activated during differentiation and not genes that are going to be repressed, indicating that, contrary to plants, it does not mark all genes with a high propensity to vary depending on the environmental conditions.

These findings uncover a possible function on gene expression of H2A.Z incorporation at gene bodies. Whether this is linked to DNA methylation, and whether and how H2A.Z presence at gene bodies participates in gene activation during differentiation are important avenues for future investigation.

## Materials and methods

### Ethics statement

Experiments involving animals were conducted according to French governmental norms. Authors have complied with all relevant ethical regulations for animal testing and research. This study was approved by the Ethics Committee of the institute “Centre de Biologie Intégrative” (FR3743) and was authorized by the French Ministry of Education and Research (approval APAFIS #4528-2016031109479615 v3).

We have complied with all relevant ethical regulations.

### Animals

Mice homozygously floxed on the *H2afz* and *H2afv* genes were obtained from RIKEN BRC (Ibaraki, Japan) and back-crossed with C57Bl/6J mice (Charles-River, L’Arbresle, France). The F1 offspring were then crossed with each other to generate *H2afz^fl/fl^ and H2afv^fl/fl^ homozygous* genitors. These mice were then crossed with the *Lgr5*-CreERT2 mouse model (from Jackson Laboratory) to obtain F1 littermates also crossed with each other to obtain *Lgr5*-CreERT2/*H2afz^fl/fl^* (thereafter called *H2a.z.1^fl/fl^*) mice, *Lgr5*-CreERT2/*H2afv^fl/fl^* (called *H2a.z.2^fl/fl^*) mice or *Lgr5*-CreERT2/*H2afz^fl/fl^&H2afv^fl/fl^* (called *H2a.z.1&2^fl/fl^*) mice. Genotyping for H2afz and Cre alleles was done by PCR analysis of tail DNA samples (see SuppTable1A for primer sequences). Note that CRE recombinase-coding sequence being inserted the Lgr5 essential stem cell marker gene to replace the coding sequence, littermates only exhibit homozygous wild-type Lgr5 genotype or heterozygous CRE/wt one.

To induce H2A.Z isoforms knock-out, 6-8 weeks-old male mice were fed a Tamoxifen-containing diet (Tam Diet TD130857 (500mg/kg), ENVIGO, USA) for ten days. The administration route and dosage had previously been studied and validated (Rispal *et al*., 2019). During treatment, macro-physiological parameters (weight, behavior, activity, aspect,…) of mice were monitored daily.

### Tissue sampling and organoids culture

After treatment, the mice were euthanized, dissected and the 2nd third of the small intestine (jejunum) was opened longitudinally. Then, cells were harvested by scraping the mucosae using a scalpel blade, before being subjected to DNA, RNA or protein extraction. The proximal part of each tissue sample was fixed in formalin before being paraffin-embedded, sectioned and used in immunofluorescence experiments.

Dissected tissues were also used to isolate intestinal epithelial stem cells plated into 3D culture to form intestinal organoids, as previously described (Sato *et al*., 2009). Briefly, EDTA-mediated dissociation of epithelial tissue was subjected to mechanical shaking to separate crypts from villi. Crypt-enriched fraction was then obtained by using 70µm filter cartridges and collecting the filtrate. After mixing with Matrigel (BD Bioscience), suspension was plated in 24-well plates. After Matrigel polymerization, IntestiCult^TM^ Organoid Growth Medium (#06005, StemCell) was added and renewed every two days.

Note that the same isolation protocol was used to obtain crypt and villus fractions for RNA and protein extraction.

### Tissue sections and immunoflurorescence

Fixed intestines were dehydrated and embedded in paraffin before being cut into 5µm sections. For immunofluorescence, tissue sections were deparaffinized with Histo-clear II (Euromedex) and then rehydrated in successive baths of ethanol (100%, 95%, 70%) and distilled water. Slides were heated in unmasking solution (Eurobio) with a pressure cooker to reveal antigens. Then tissue sections were permeabilized for 5 minutes with Triton X-100 1% and blocked with BSA 1% for 45 minutes. Sections were incubated with primary antibodies (diluted 1:500) for 2 hours at room temperature. The following antibodies were used: H2A.Z (#39114, Active Motif), Sucrase-Isomaltase (sc-393470, Santa-Cruz), Mucin 2 (sc-15334, Santa Cruz), Lysosyme (ab108508, Abcam), Chromogranin-A (#251190, Abbiotec), Olfm4 (#39141, Cell Signaling Technology) and Ki67 (ab15580, Abcam). Sections were incubated with secondary antibodies (dilution 1:500) coupled with Alexa Fluor® (Fisher Life Science, anti-mouse -AF488 #A21202 and -AF594 #A11032; anti-rabbit -AF488 #A11034 and -AF594 #A11037) for 1 hour at room temperature, and nuclei were stained with DAPI for 3 minutes at room temperature. Finally, slides were mounted with ProLong® (Fisher life Science) and stored at 4°C.

### Cell culture, transfections and treatments

The colorectal cancer cell line Caco-2/15 was obtained from the Jean-François Beaulieu’s lab (Université de Sherbrooke, Québec, Canada), cultured in Dulbecco’s Modified Eagle’s Medium (DMEM) and splitted every 3 days at sub-confluence. Cells were transfected with siRNA (see SuppTable1B for sequences) by electroporation (Amaxa) and analysis was performed 3 days after transfection.

In overexpression experiments, H2A.Z.1 or H2A.Z.2 expression vectors (kind gift from Peter Cheung, University of Toronto, ON Canada) transfected using Caco-2 Avalanche^TM^ transfection reagent (EZT-CACO-1, EZ Biosystems), alone (for RT-qPCR and proliferation assays) or in combination with PGL3 vector encoding for minimal promoter for SI gene driving the expression of Firefly luciferase and pRLTK normalizer vector constitutively expressing Renilla luciferase (kind gift from Jean-François Beaulieu) (for luciferase assays).

For proliferation assays, transfected cells were seeded in 96-well plates (2000 cells per well). To assess cell proliferation, cells were incubated with the cell proliferation reagent WST-1 (Sigma) for 2h at 37°C, and cell numbers were measured by optical density at 450nm.

### Chromatin Immunoprecipitation and co-IP

ChIP experiments were performed using standard procedures. Cells were fixed with formaldehyde 1% for 15 minutes, then cells and nuclei were lysed. The recovered chromatin was sonicated: 10 cycles of 10 seconds (1s on, 1s off) and precleared. 50µg for H2A.Z and 200µg for CDX2 of chromatin were used for immunoprecipitation with 2µg of anti-Flag antibody (M2 clone, Sigma-Aldrich). Then, chromatin was incubated with A/G beads for 2 hours. Crosslinking was reversed by incubation of beads with SDS at 70°C and proteins were degraded with proteinase K. Finally, DNA was purified using the GFX™ DNA purification kit (GE Healthcare), and ChIP was analyzed by qPCR using specific primers (see SuppTable1C). For co-IP experiment, cells were lysed in IP buffer (10 mM Tris pH 8, 0.4% NP40, 300 mM NaCl, 10% glycerol, 1 mM DTT, antiprotease (Roche), anti-phosphatase (Sigma) inhibitors). Lysates were diluted using one volume of dilution buffer (10 mM Tris pH 8, 0.4% NP40, 5 mM CaCl2, 2 U/mL RQ1 DNAse (Promega)) and then incubated with anti-Flag or anti-p400 antibodies coupled to agarose beads (Sigma) overnight at 4°C. Beads were washed four times with IP buffer, eluted with Laemmli sample buffer without reducing agent and analyzed by western blotting.

### CRISPR-Cas9 -mediated genome editing

We used the ouabain based co-selection strategy, as we previously described in (Rispal *et al*., 2019) and in (Lamaa *et al*., 2020). Briefly, we co-transfected of a RNA guide + DNA donor that are able to provide cells with ouabain resistance following genome editing in the ATP1A1 gene (Na+/K+ pump) (Agudelo *et al*., 2017). Cells were thus transfected using JetPEI (Polyplus) and plasmids encoding for Cas9, PAM and guide for NA/K ATPase and PAM sequences allowing the targeting of *H2AFZ* and *H2AFV*. Three days after transfection, 0.4µM ouabain was added for 72 hours to select recombinant resistant clones. Following clones isolation, screening of genome editing at the *H2AFZ* and *H2AFV* genes was performed by genomic DNA purification and out-out PCR as described (Lamaa *et al*., 2020). Heterozygous clones were then checked for correct genome editing by sequencing. Screening primers and guides sequences are detailed in SuppTable3D.

### RT-qPCR and western-blot analysis

Genotyping for *H2afz* alleles in intestinal epithelial samples was done by PCR analysis of DNA samples (see SuppTable3A for primer sequences).

For RT-qPCR, total RNA from cells or tissues was extracted using the RNeasy Kit (Qiagen) according to the manufacturer’s protocol, then it was reverse transcribed into cDNA using AMV reverse transcriptase (Promega). Finally, qPCR was performed using specific primers (SuppTable1E), and RPLP0/Rplp0 and β2M/β2m were used as control normalizing genes.

For western-blot, proteins were extracted in lysis buffer (Triton X100 1%, SDS 2%, NaCl 150mM, NaOrthovanadate 200µM, Tris/HCl 50mM). Proteins were separated in NuPage BisTris 4-12% gels (Invitrogen), then transferred onto PVDF membranes. Membranes were incubated with primary antibodies (diluted 1:500): p400 (ab5201, Abcam), SRCAP (ESL103, Kerafast), SI (NBP1-62362, Novus), Lactase (NBP2-50513, Novus), CDX2 (ab88129, Abcam), Flag (M2, Sigma-Aldrich), H2A.Z (Abcam, ab4174), and GAPDH (Chemicon, Mab374). Finally, they were incubated with secondary antibodies (dilution 1:500) coupled with HRP (BioRad, anti-mouse-HRP #1706516 and anti-rabbit-HRP #1706515), and revealed with Lumi-lightPLUS substrate (Roche).

### Genome-wide analysis

#### RNA-sequencing

Samples for RNA-sequencing experiments were prepared in duplicates from subconfluent (Sc) or 7 days post-confluent (Pc) populations of Caco-2/15 cells expressing flagged version of H2A.Z.1 or H2A.Z.2 isoform. 1ug of total RNA of each sample was submitted to EMBL-GeneCore (Heidelberg, Germany), where the ribosomal RNA depletion and the library preparation were done using Truseq Stranded Total RNA protocol (Illumina). Paired-end sequencing was performed by Illumina’s NextSeq 500 technology with a depth of sequencing of at least 59M reads per sample. Two replicates of each sample were sequenced. The quality of each raw sequencing file (fastq) was checked with FastQC. First, total reads were aligned to the rDNA genome and unmapped reads were aligned onto reference human genome (hg38) in paired-end mode with STAR Version 2.5.2b (Dobin *et al*., 2013) and processed (sorting and indexing) with samtools (Li *et al*., 2009) keeping only primary alignment. Raw reads were counted by exons, per gene_id, using HT-seq Version 0.6.1 (Anders *et al*., 2015) on the NCBI RefSeq annotation .gtf file from UCSC, in a strand specific mode with default parameters. Differential analysis (subconfluent (Sc) versus postconfluent (Pc) cell populations) were done with DESeq2 Bioconductor R package, Version 1.22.1 (Love *et al*., 2014) with default parameters, BH correction and HKG normalization using RPLP0 and B2M genes, and filtering genes with less than 1 read per sample. Gene Ontology analysis of TOP1000 most upregulated or downregulated genes have been done using GeneCodis4 (Garcia-Moreno *et al*., 2022) (SuppFig.2C).

#### ChIP-sequencing

For ChIP-seq experiments, Caco-2/15 cells expressing Flag-H2A.Z.1 and Flag-H2A.Z.2, 100 µg of chromatin supplemented with 10 ng of Spike-in chromatin (Active Motif) were used per ChIP experiment. For each reaction, 4 µg of Flag M2 antibody (Sigma) and 1 µg of Spike-in antibody (Active Motif), were used.

About 10 ng of immunoprecipated DNA (quantified with quantiFluor dsDNA system, Promega) were obtained at each sample, and samples were submitted to EMBL-GeneCore Heidelberg for sequencing, that was performed by Illumina’s NextSeq 500 technology (single-end, 85 bp reads). The quality of each raw sequencing file was verified with FastQC. Files were aligned to the reference human genome (hg38) in single-end mode with BWA (Li and Durbin, 2009) and processed (sorting, PCR duplicates, minimum quality of 25 and indexing) with Samtools. The coverage was computed with Deeptools bamCoverage (Ramirez *et al*., 2016) using exactScaling parameters and CPM normalization.

The enriched region over the background were identified by peak calling using MACS3 callpeak (Zhang *et al*., 2008) with 0.001 q-value cutoff. Only peaks common between replicates are kept and merged. Gene Ontology analysis have been done using GeneCodis4 (Garcia-Moreno *et al*., 2022) (SuppFig.4).

Global heatmap gene profile across experiment (Fig.1B) were computed using Deeptools computeMatrix and Deeptools plotHeatmap in a window of +/- 2 kb around TSS filtering genes shorter than 5kb. To obtain experiment metaprofiles, Deeptools computeMatrix scale-region are used to generate fixed windows between TSS-2kb to TSS+4b and TES to TES+3kb and a scale-region on gene body (parameters: “-bs 50 -m 10000 -b 6000 -a 3000 --skipZeros – missingDataAsZero”). Rscript were used to load gene matrix and generate the diverse visualizations with tidyverse, plyranges, sparklyr, data.table and ggvenn libraries. Genes were classified into eight expression classes (Fig.1C) based on the number of normalized HKG reads in RNA-seq.

The TOP500 most up-or down-regulated genes in RNA-seq were also selected using quartile on Log2(FoldChange) expression.

## Data availability

The authors declare that all data supporting the findings of this study are available within the paper and its supplementary information files or from the corresponding author [FE] upon request.

All data from genome-wide experiments were deposited to Gene Expression Omnibus (GEO) public functional genomics data repository with the following reference: GSE232114 (https://www.ncbi.nlm.nih.gov/projects/geo/query/acc.cgi?acc=GSE232114)

## Acknowledgements

We thank Dr Arnaud Besson and Yvan Canitrot (CBI-MCD) for their technical support in mice sampling and CRISPR-Cas9 experiments, respectively. We also thank Paul Duchesne and all the DT’s team for their help or helpful discussions.

We acknowledge the CBI animal facility for housing the mice, the CBI-LITC facility for microscopy experiments analysis, and the CBI-Big-A bioinformatics platform members (Marion Aguirrebengoa and Vincent Rocher) for their biostatistical support.

This work was supported by a grant from the Ligue Nationale Contre le Cancer as a “Equipe labellisée” to DT. JR was recipient of a studentship from the French Ministry of Research.

## Author contributions

J.R., C.R., V.J., C.L., L.D., M.C.-B., F.E.: Acquisition of data, analysis and interpretation of data.

J.R., M.C-B., V.J., D.T., F.E.: Critical revision of the manuscript for important intellectual content.

J.R., V.J., F.E., DT: Statistical analysis, writing of the manuscript.

D.T., F.E.: Study supervision.

D.T.: Obtained funding

