## Supplemental material for "Control of intestinal stemness and cell lineage by histone variant H2A.Z isoforms"

### Supp Fig 1

A

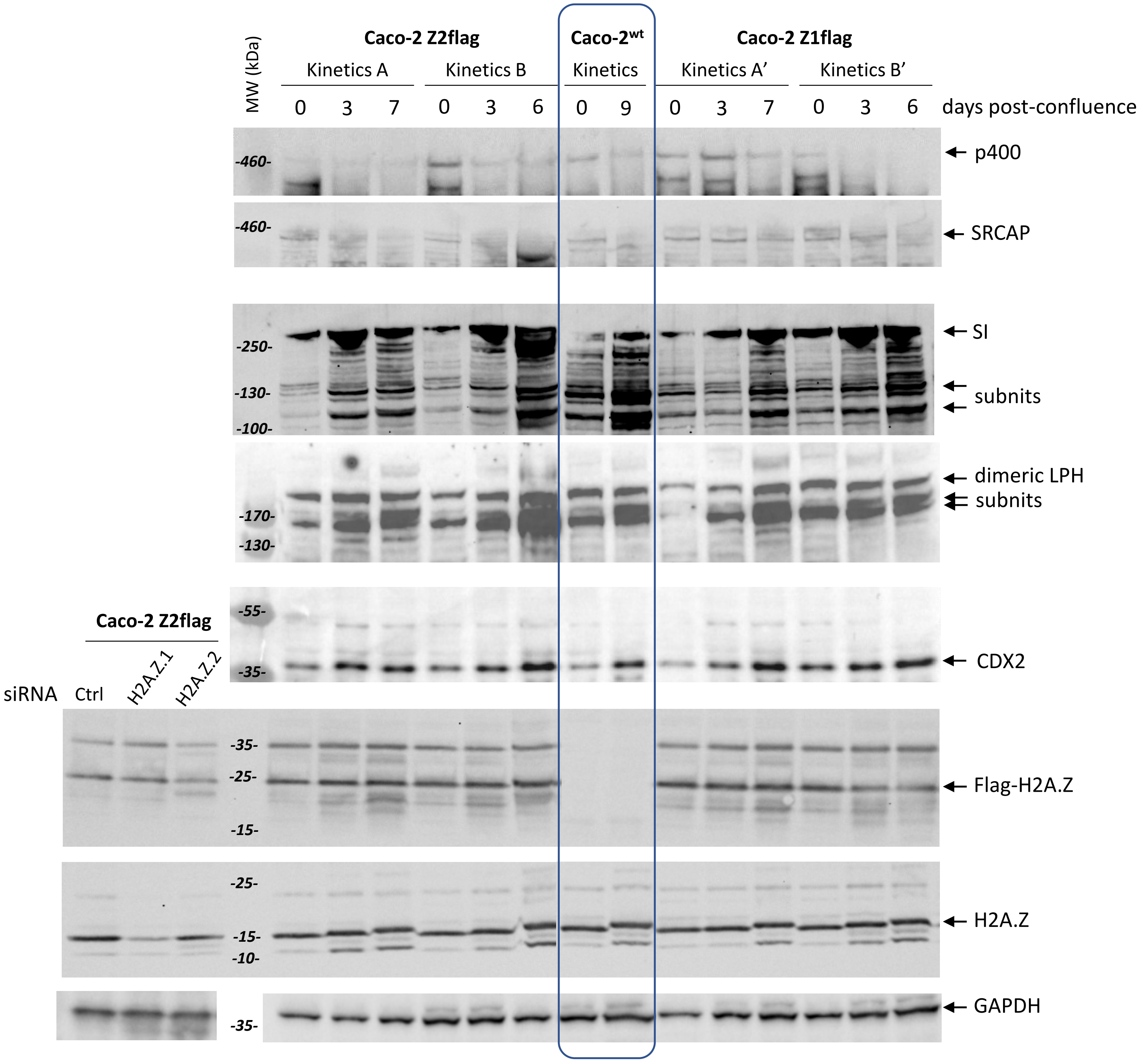
Caco-2 Z2flag
Caco-2$^{wt}$
Caco-2 Z1flag
Kinetics A
Kinetics B
Kinetics
Kinetics A'
Kinetics B'
MW (kDa)
0 3 7 0 3 6 0 9 0 3 7 0 3 6
days post-confluence
p400
SRCAP
SI
subnits
dimeric LPH
subnits
CDX2
Flag-H2A.Z
H2A.Z
GAPDH
Caco-2 Z2flag
siRNA
Ctrl
H2A.Z.1
H2A.Z.2
-460-
-460-
-250-
-130-
-100-
-170-
-130-
-55-
-35-
-35-
-25-
-15-
-25-
-15-
-10-
-35-

**Supp Fig 1**

**B**

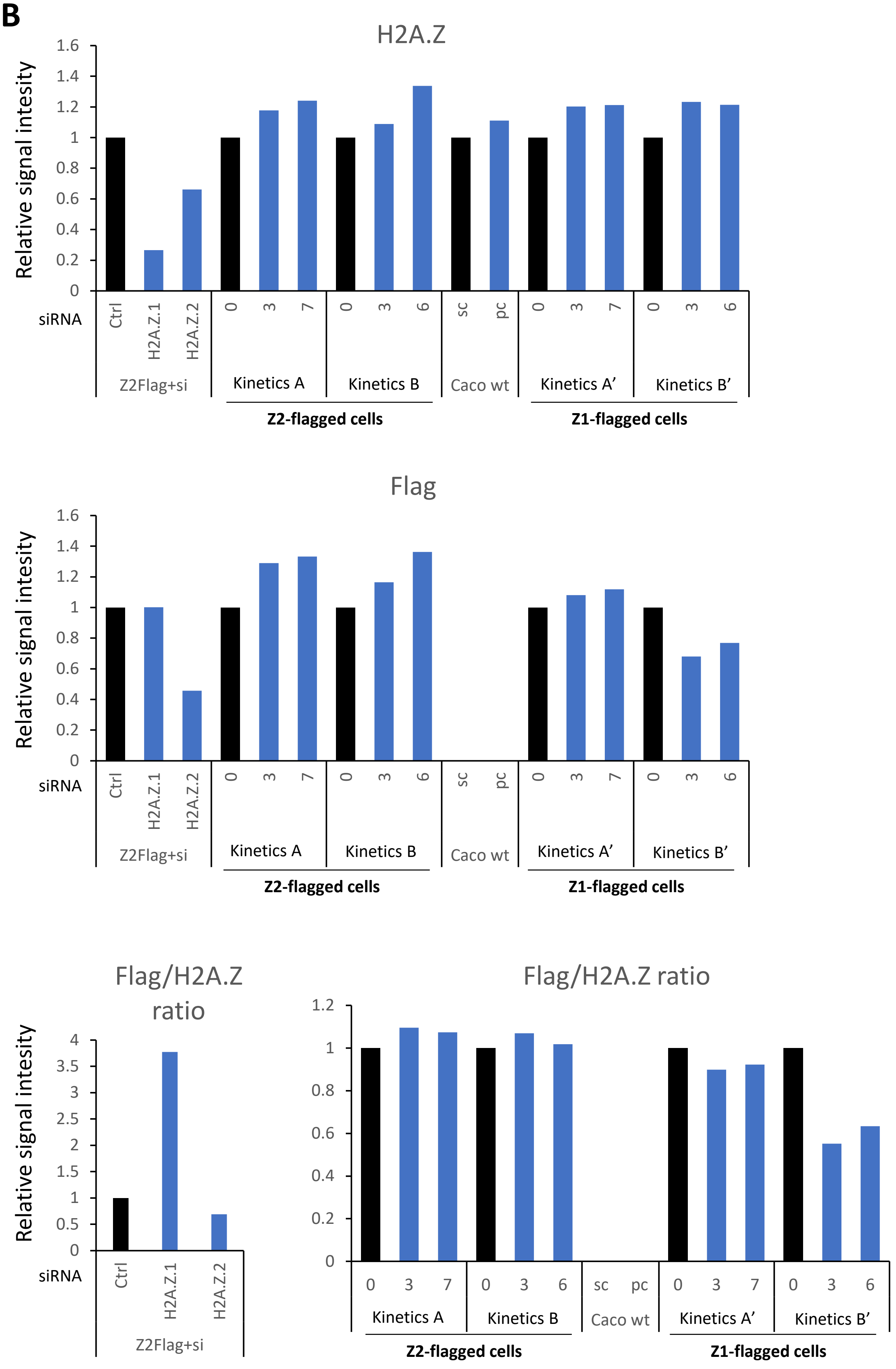


**Supplementary Figure 1: CRISPR-Cas9 cell models characterization and western-blot quantification.**
**A:** Western-blot characterization of these models. As for parental wild-type Caco-2/15 cells (boxed lines), the genome edited cell lines expressing tagged version of H2A.Z.1 or H2A.Z.2 show a decrease in p400 and SRCAP expression with the confluence, which correlates the increase of CDX2 and LPH and SI expression. **B:** Quantification of bands in western-blot shown in (A) for wild-type and tagged H2A.Z isoforms. Each point of siRNA experiment or kinetics (in blue) is represented relative to their own control (in black). Note that there is a slight increase in the global H2A.Z level upon confluence. However, while H2A.Z.2-flagged version follow the same tendency, H2A.Z.1-flag is almost stable or decreasing during the confluence process. This causes the global increase in H2A.Z.2/H2A.Z.1 ratio during the enterocyte-like differentiation.

**Supp Fig 2**

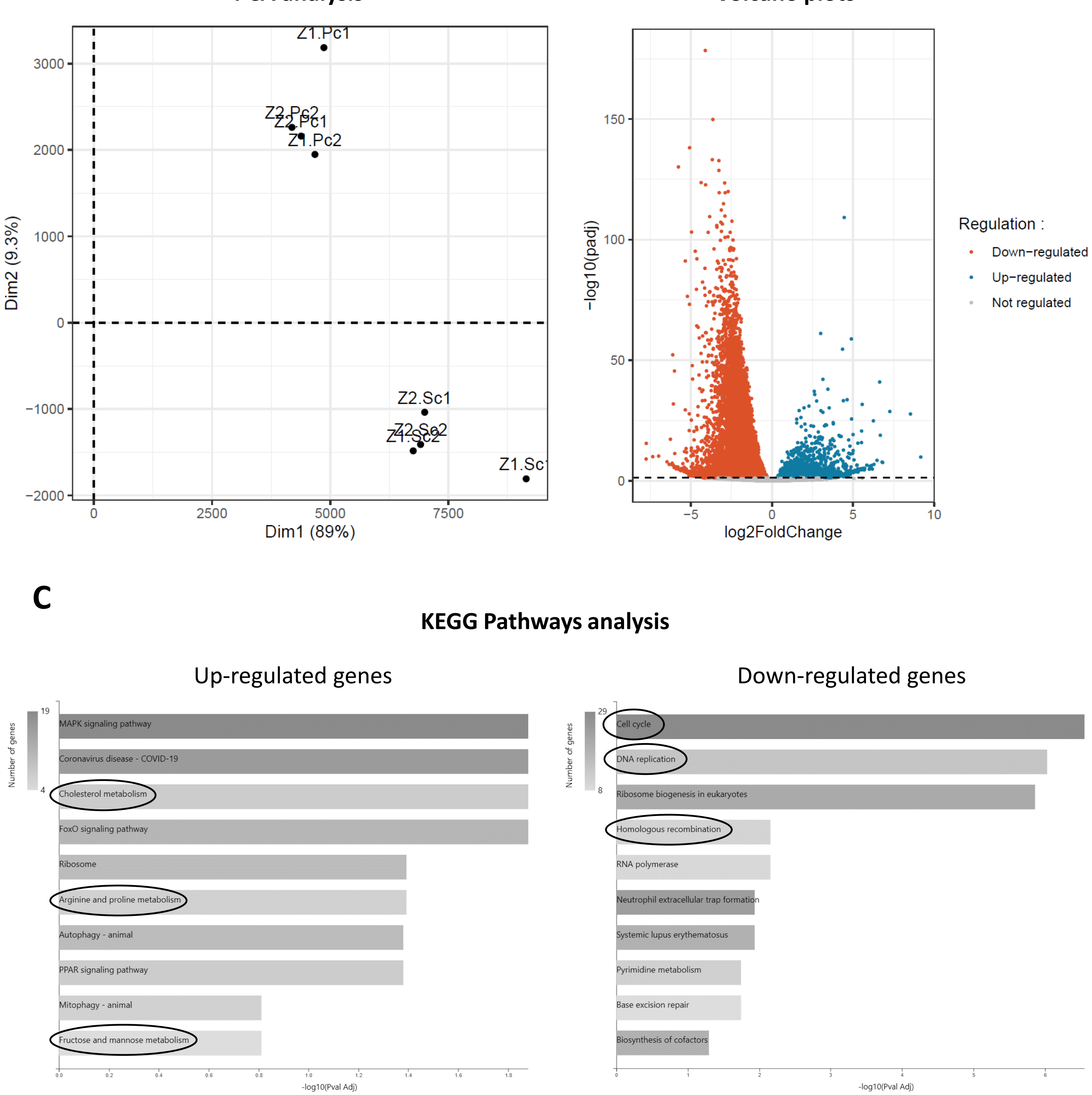


**Supplementary Figure 2: Gene ontology of differentially expresssed genes during differentiation of Caco-2/15 cells.**
**A:** Principal Component Analysis of RNA-seq centered data after HKG normalization using FactoMineR R library.
**B:** Representation of differentially expressed genes in post-confluence and sub-confluence cells using a volcano plot. Based on the RNA-seq analysis, genes are plotted using log2-fold change (x-axis) and -log10 adjusted p-value (y-axis) parameters. Differentially expressed genes have an adjusted p-value above 0.05 (dashed line). **C:** A gene ontology analysis of significatively expressed (more than 50 reads) TOP1000 upregulated and downregulated genes was performed. The first 10 KEGG Pathways are shown ordered by adjusted p-value.

**A** PCA analysis

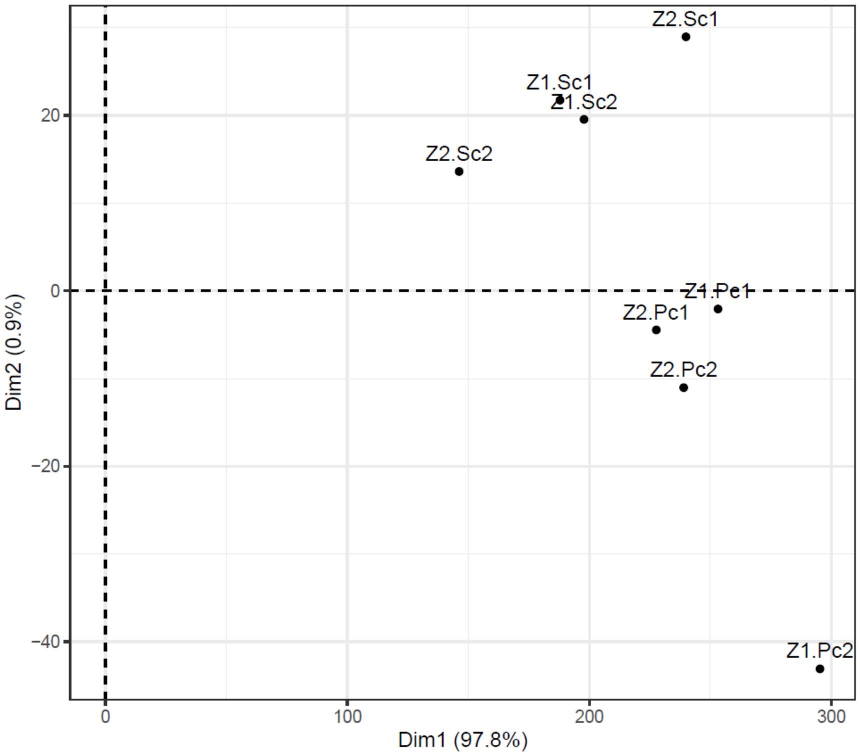


**B**

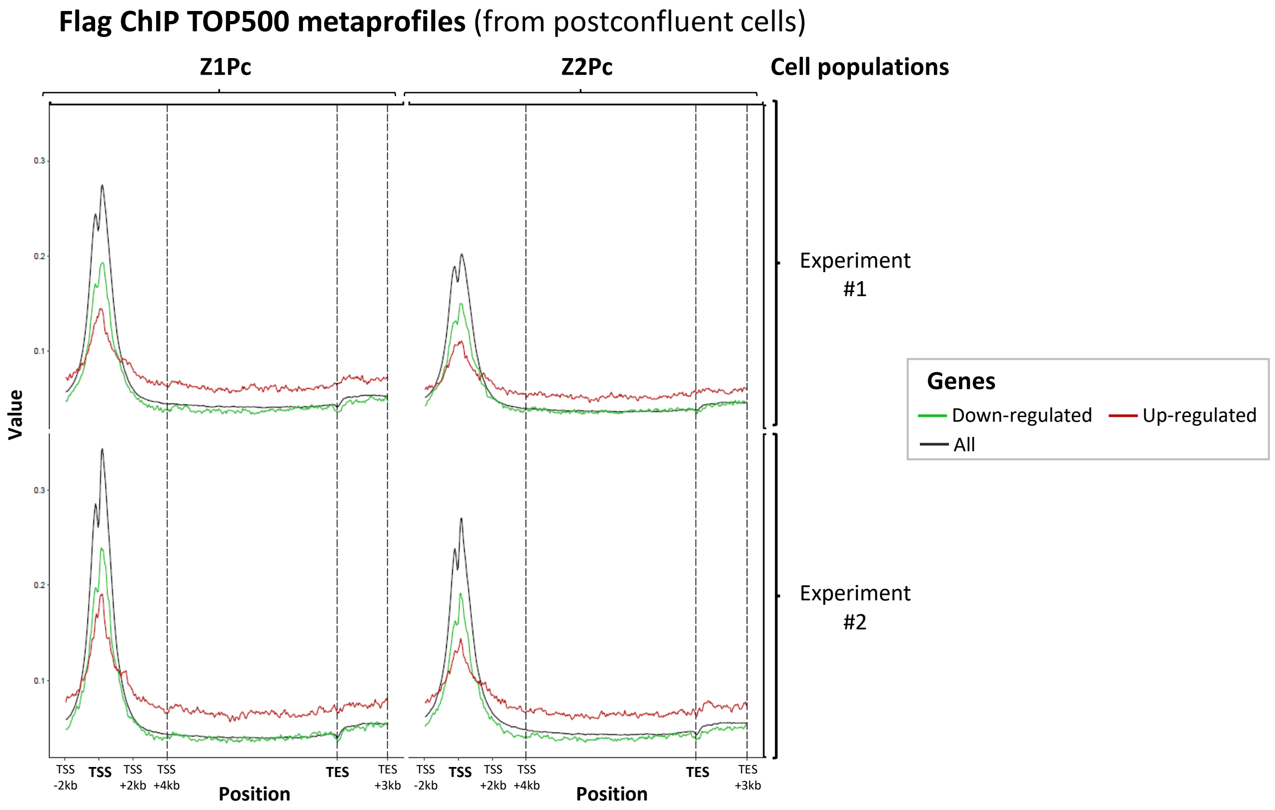


<u>Supplementary Figure 3</u>**: ChIP-seq PCA analysis and Metaprofiles for H2A.Z.1 or H2A.Z.2 recruitment on TOP500 regulated genes during the differentiation-like process of Caco-2/15 cells.**
**A:** Principal Component Analysis of the ChIP-seq data considering raw peak counts on non-overlapping TSS and gene body regions. **B:** Metaprofile in post-confluent cell populations. Based on RNA-seq data obtained in both sub-confluent and 7-days post-confluent cells, genes were classified by their expression fold change. Metaprofiles for TOP500 upregulated (in red) or downregulated (green) genes, as well as for all genes (black) of the RNA-seq experiments were drawn using enrichment data obtained in postconfluent populations.

**Supp Fig 4**

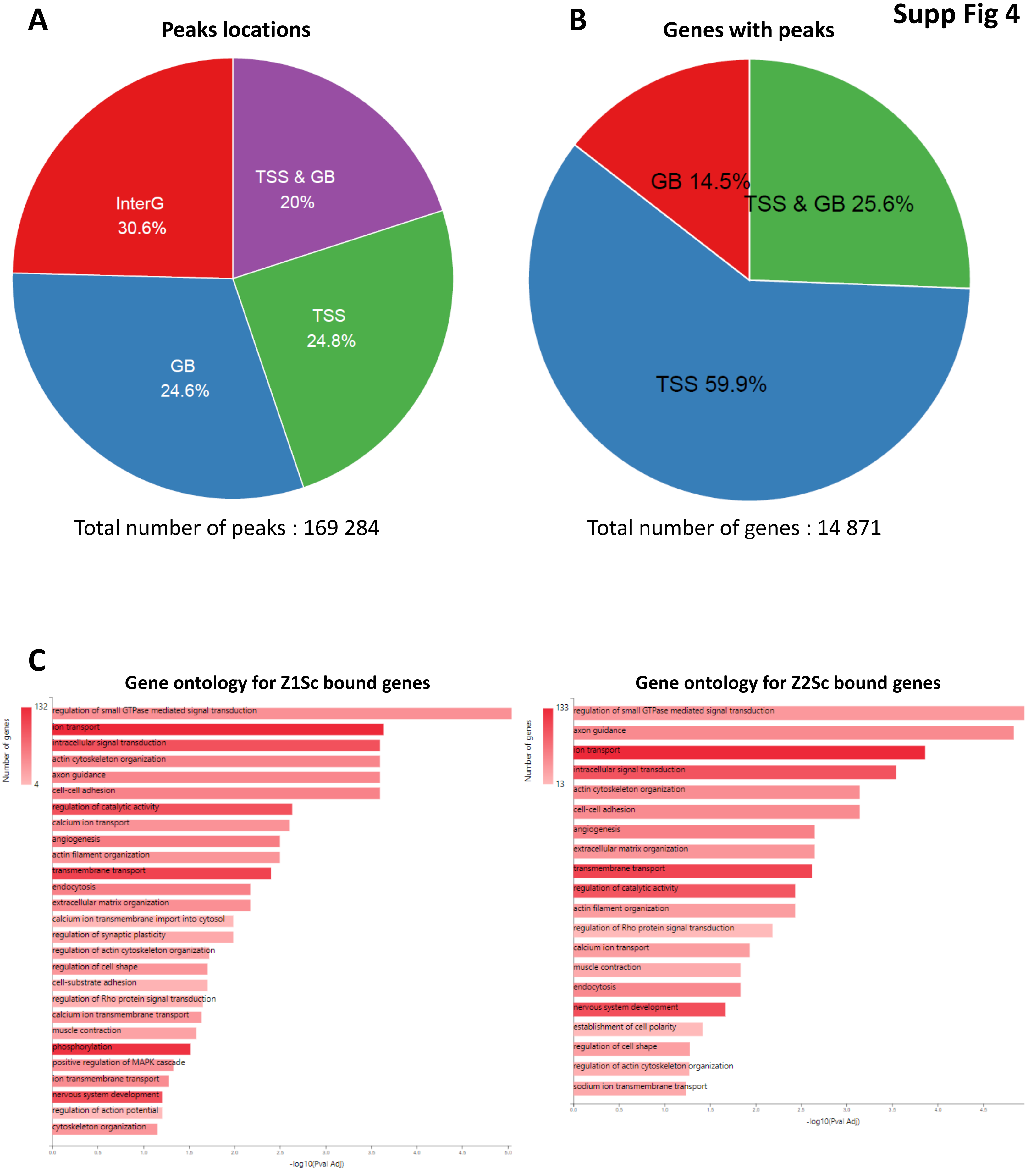


<u>**Supplementary Figure 4**</u>**: Distribution of H2A.Z peaks along the genome.**
**A:** Pie chart showing the proportion of peaks localized within intergenic regions, TSS, gene bodies or overlapping TSS and a gene body. Total number of detected peaks is also shown. **B:** Pie chart showing the percentage of genes with peaks in their body, at their TSS or at both locations. Total number of genes with peaks is shown. **C:** Gene ontology for biological processes -related genes with positive log2(GB/TSS) for H2A.Z.1 or H2A.Z.2 in subconfluent cells.

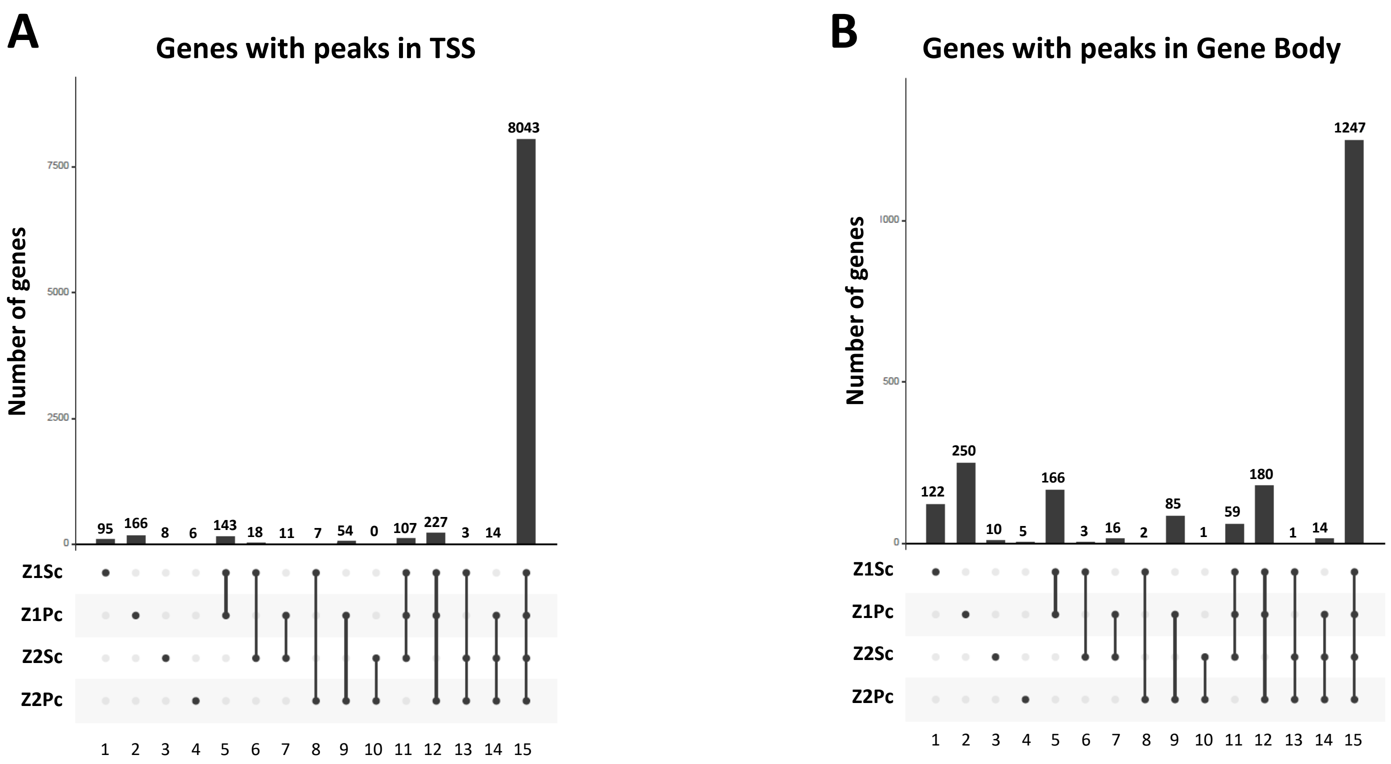


**<u>Supplemental Figure 5</u>: Localization of H2A.Z isoforms peaks within genes regarding cell state and expression level**

**A**: Upset representation of the number of genes with peaks in their TSS regions (-1kb to +1kb), regarding the H2A.Z isoform and differentiation state of cells. **B:** same as in A for genes with peaks in their gene body.

A

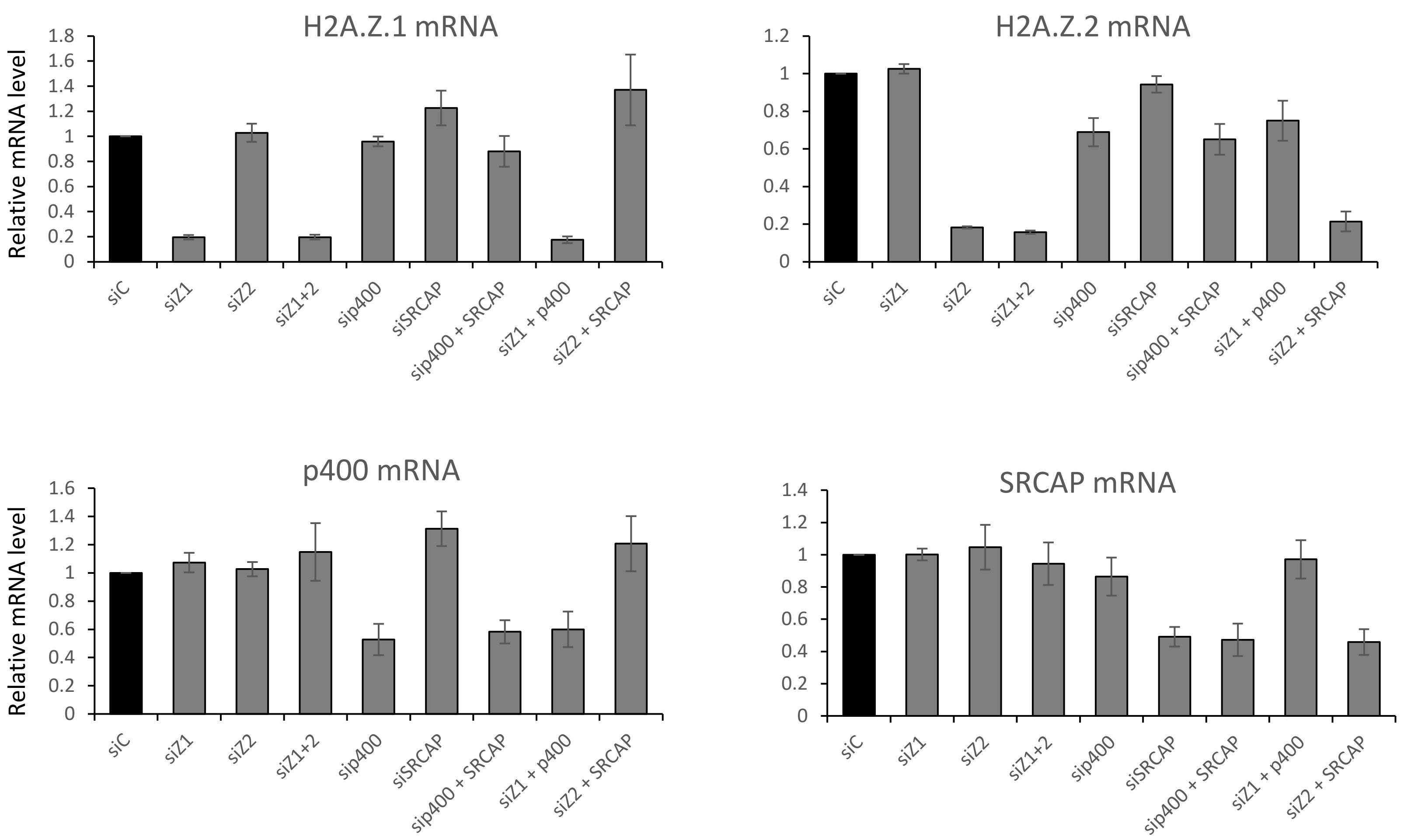


B

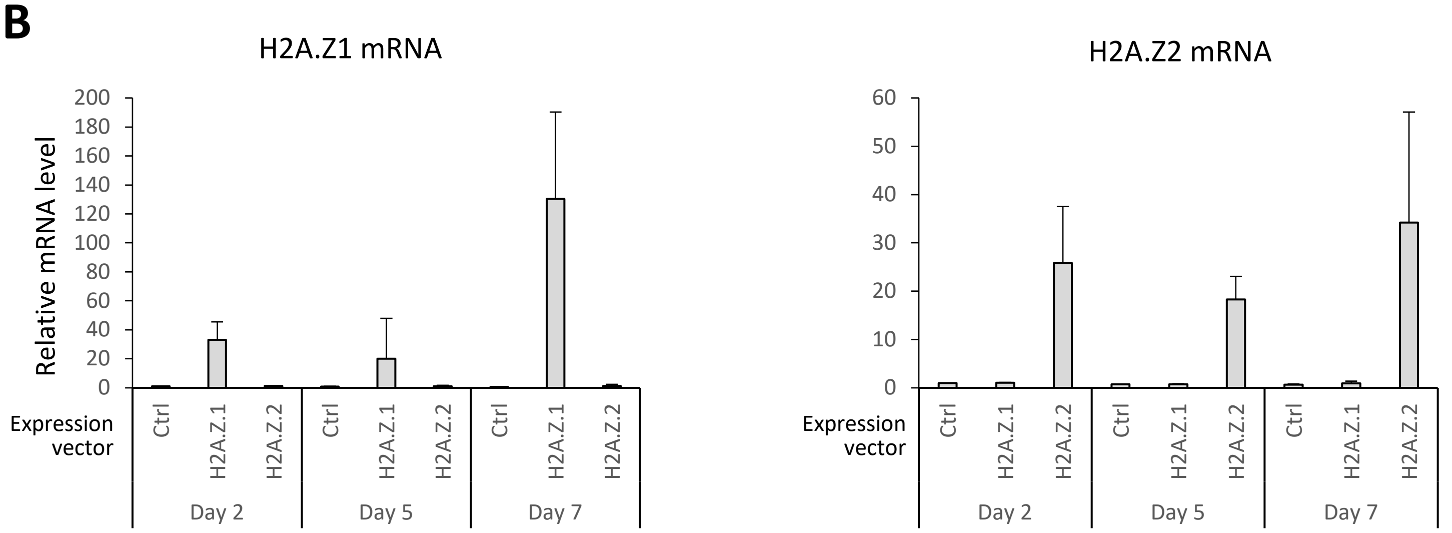


<u>Supplementary Figure 6</u>: **Quantification of mRNA in siRNA-mediated depletion or vector-mediated overexpression experiments in Caco-2/15 cells.**

**A:** Quantification of mRNA encoding for isoforms H2A.Z.1 and H2A.Z.2, as well as for their incorporators p400 and SRCAP, 3 days after siRNA transfection. The mean and standard deviation from five independent experiments are shown. **B:** Quantification of indicated mRNA in cells transfected with H2A.Z isoforms expression vectors. Cells were harvested after the indicated times. The mean and standard deviation from three independent experiments are shown.

### Supp Fig 7

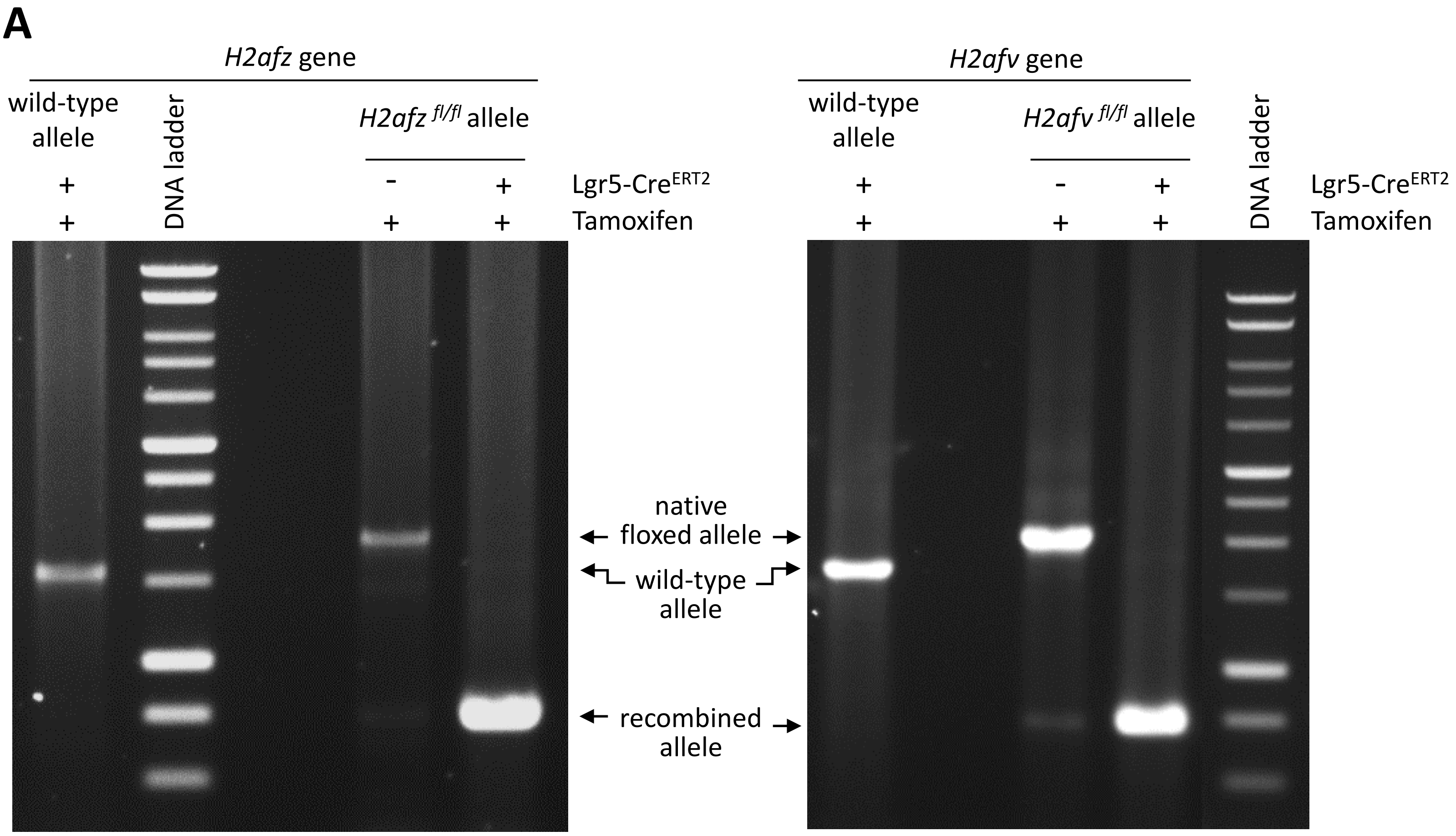


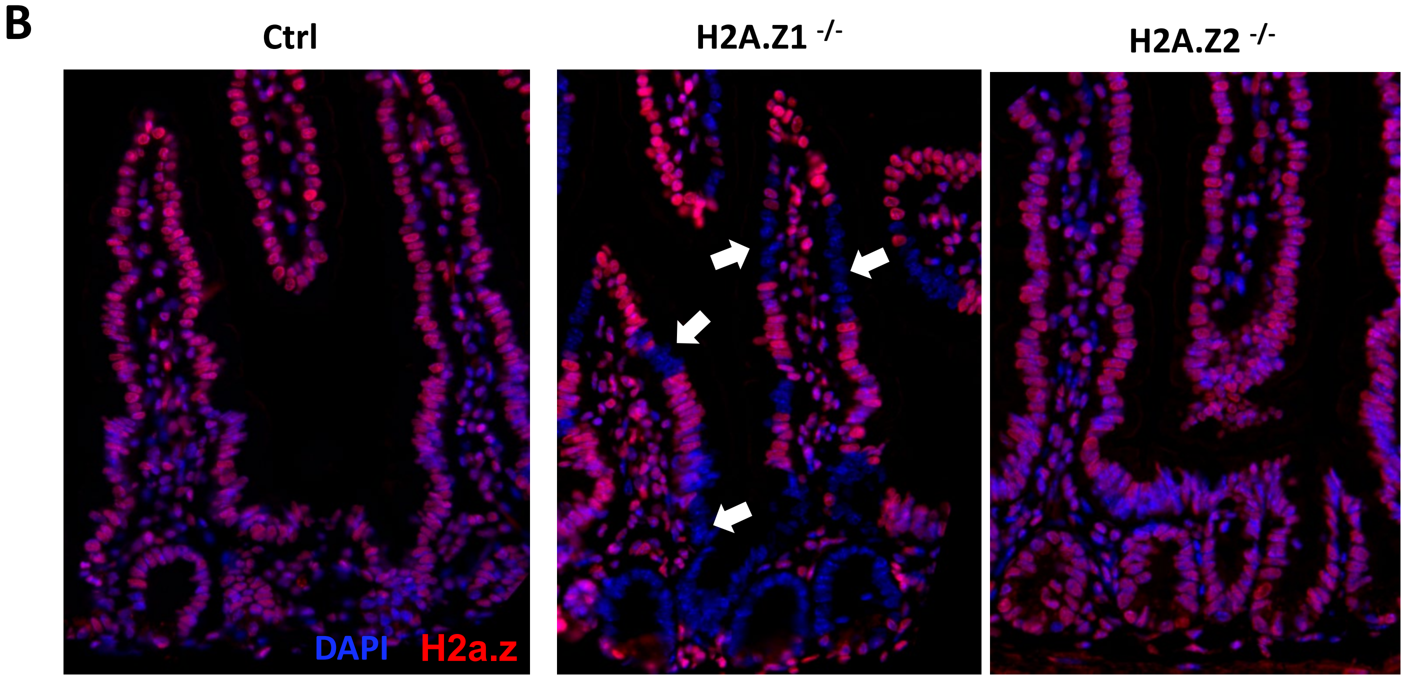


**Supplementary Figure 7: Characterization of inducible knockout mice models.**

**A:** Genomic DNA extracted from intestinal epithelium of representative mice, habouring the indicated genotypes, and after Tamoxifen treatment, was analyzed by PCR. The DNA recombination is observed only *Lgr5-CRE*$^{ERT2}$ positive strains. Note however that the larger non-recombined allele will certainly have disadvantage for amplification compared to the shorter recombined band and could be minored in this panel (recombination efficiency may be overestimated). **B:** Immunoflurescence on intestinal sections using anti-H2A.Z antibody. Mosaic depletion in the staining is observed in *H2a.z.1*$^{fl/fl}$ mice (arrows), but no major decrease in H2A.Z staining is observed upon H2A.Z.2 knockout (in *H2a.z.2*$^{fl/fl}$ model), probably due to the respective abundance of the two isoforms.

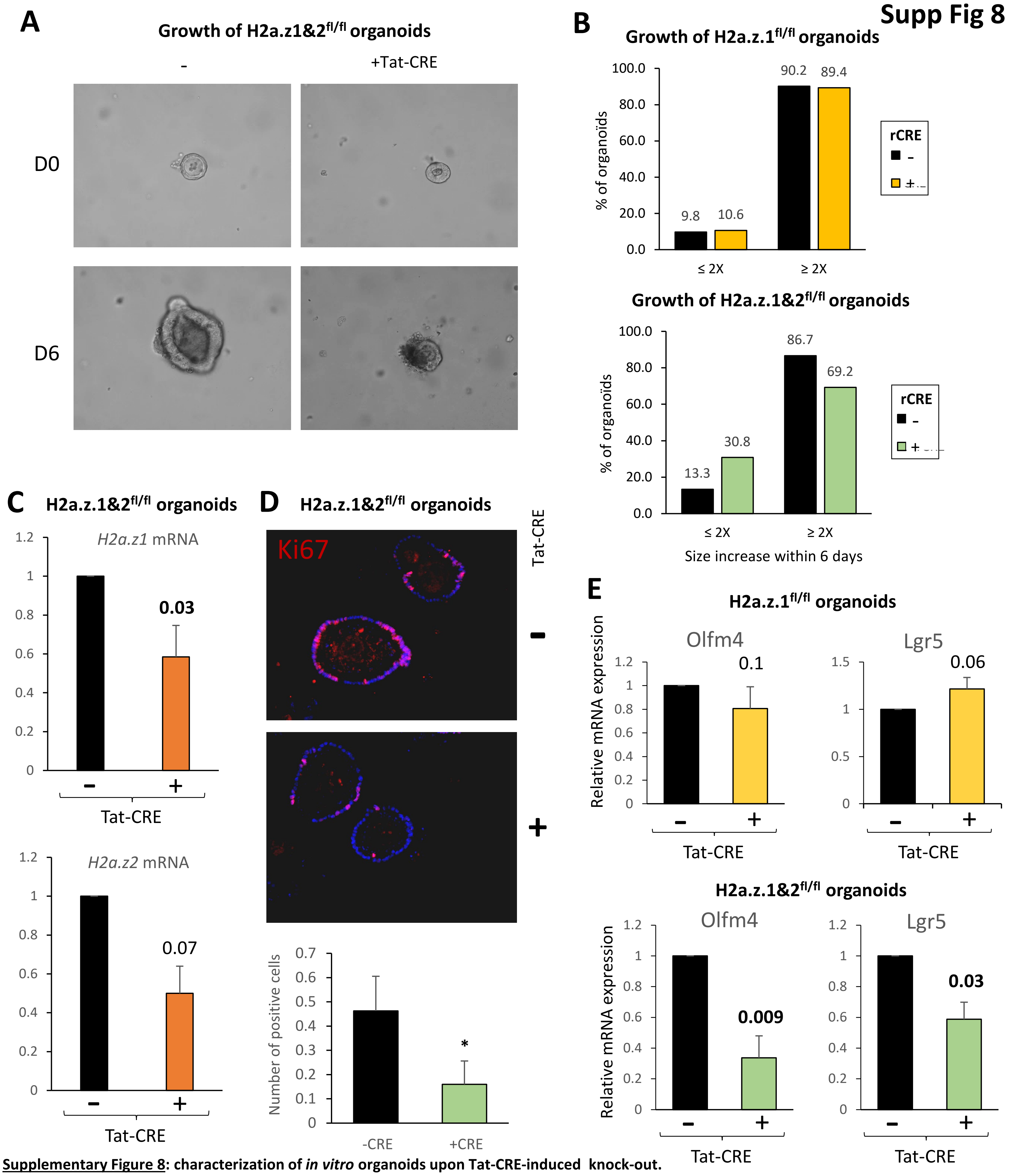


**Supplementary Figure 8**: **characterization of *in vitro* organoids upon Tat-CRE-induced knock-out.**

**A**: Representative picture of growth of H2a.z.1&2$^{fl/fl}$ organoids treated with (+) or without (-) Tat-CRE. **B**: Measurements of the 3D *in vitro* growth of *H2a.z.1$^{fl/fl}$* or *H2a.z.1&2$^{fl/fl}$* organoids with (+) or without (-) *in vitro* treatment with recombinant Tat-CRE. Size measurements (around 100 organoids from 3 mice per condition) are performed the day of treatment (D0) and 6 days after (D6). Organoids are pooled in 2 classes: those whose size increases less (≤ 2X) or more (≥ 2X) than 2 times in 6 days. **C**: H2a.z.1 and H2a.z.2 expression in H2a.z.1&2$^{fl/fl}$ organoids treated with (+) or without (-) Tat-CRE. The mean and standard deviation from five independent experiments are shown. Statistical analysis was done using the one-way Student's t-Test and the p-values of treated organoids compared to untreated are stated in the graphic. **D**: Ki67 immunostaining on sections of organoids from *H2a.z.1&2$^{fl/fl}$* mice treated or not with recombinant Tat-CRE recombinase. Nuclei are stained with DAPI in blue. **E**: Olfm4 and Lgr5 expression in H2a.z.1$^{fl/fl}$ and H2a.z.1&2$^{fl/fl}$ organoids treated with (+) or without (-) Tat-CRE. The mean and standard deviation from five independent experiments are shown. Statistical analysis was done using the one-way Student's t-Test and the p-values of treated organoids compared to untreated are stated in the graphic

**Supp Fig 9**

**A**

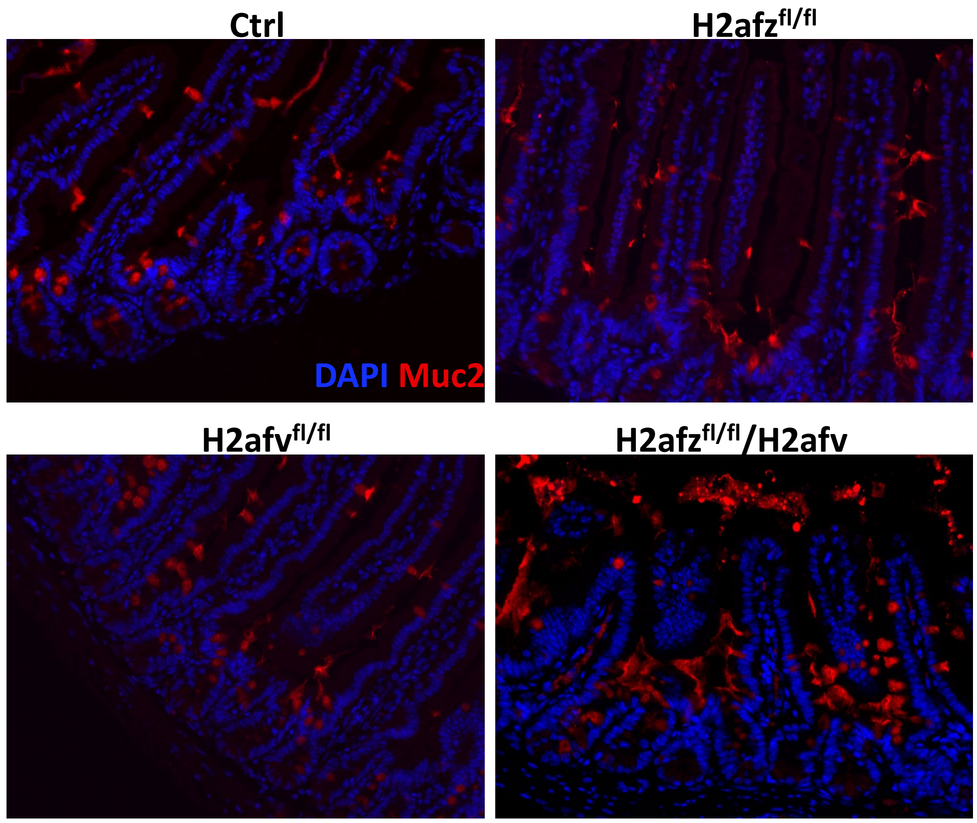


**B**

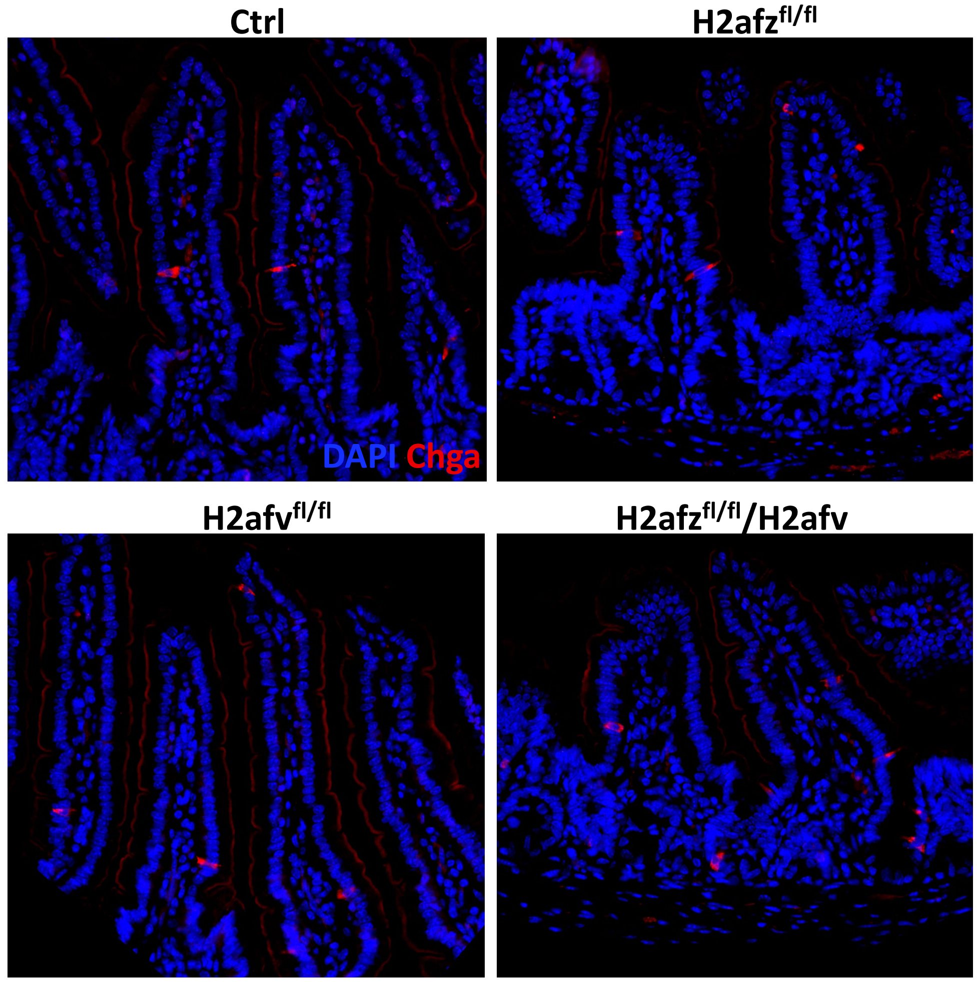


**<u>Supplementary Figure 9</u>: Immunohistofluorescence for Mucin2 and ChromograninA staining on jejunum sections of mice from indicated genotypes.**

**A:** Immunoflurescence on intestinal sections using anti-Muc2 antibody. DAPI was used to stain nuclei. **B:** same as in A for ChgA staining.

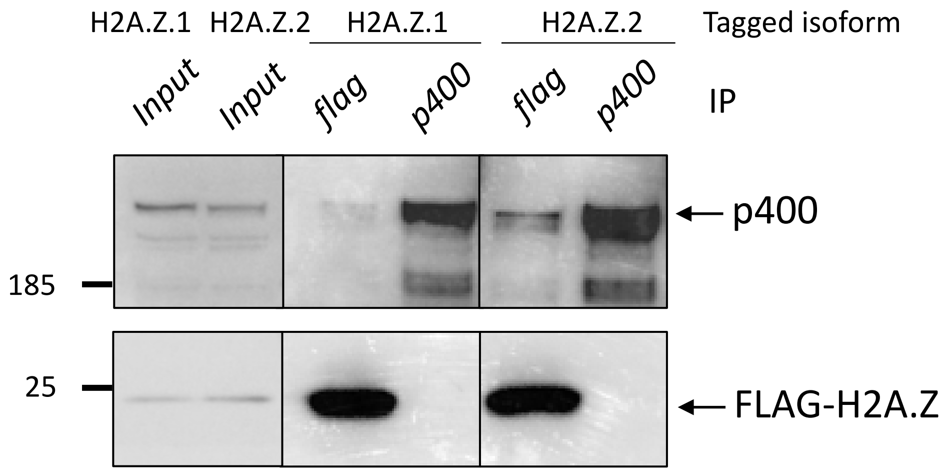


**<u>Supplementary Figure 11</u>: Immunoprecipitation with anti-Flag or anti-p400 antibodies in Caco-2/15 cells tagged for H2A.Z.1 or H2A.Z.2.**

Extracts from Caco-2/15 cells expressing either H2A.Z.1-Flag or H2A.Z.2-Flag following gene editing were immunoprecipitated by the indicated antibody. Immunoprecipitates were analyzed by western-Blot with anti-Flag or anti-p400 antibodies.

| gene_id | gene_name | **Z1Sc condition** |
|---|---|---|
| ENSG00000001084 | GCLC | |
| ENSG00000001626 | CFTR | |
| ENSG00000001630 | CYP51A1 | |
| ENSG00000002016 | RAD52 | |
| ENSG00000002726 | AOC1 | |
| ENSG00000002746 | HECW1 | |
| ENSG00000002919 | SNX11 | |
| ENSG00000003436 | TFPI | |
| ENSG00000004139 | SARM1 | |
| ENSG00000004846 | ABCB5 | |
| ENSG00000005020 | SKAP2 | |
| ENSG00000005102 | MEOX1 | |
| ENSG00000005187 | ACSM3 | |
| ENSG00000005243 | COPZ2 | |
| ENSG00000005379 | TSPOAP1 | |
| ENSG00000005471 | ABCB4 | |
| ENSG00000005486 | RHBDD2 | |
| ENSG00000005844 | ITGAL | |
| ENSG00000005882 | PDK2 | |
| ENSG00000005961 | ITGA2B | |
| ENSG00000006016 | CRLF1 | |
| ENSG00000006071 | ABCC8 | |
| ENSG00000006194 | ZNF263 | |
| ENSG00000006210 | CX3CL1 | |
| ENSG00000006282 | SPATA20 | |
| ENSG00000006283 | CACNA1G | |
| ENSG00000006453 | BAIAP2L1 | |
| ENSG00000006652 | IFRD1 | |
| ENSG00000006715 | VPS41 | |
| ENSG00000007047 | MARK4 | |
| ENSG00000007171 | NOS2 | |
| ENSG00000007216 | SLC13A2 | |
| ENSG00000007237 | GAS7 | |
| ENSG00000007402 | CACNA2D2 | |
| ENSG00000007516 | BAIAP3 | |
| ENSG00000008324 | SS18L2 | |
| ENSG00000008441 | NFIX | |
| ENSG00000008516 | MMP25 | |
| ENSG00000008710 | PKD1 | |
| ENSG00000008838 | MED24 | |
| ENSG00000008853 | RHOBTB2 | |
| ENSG00000009790 | TRAF3IP3 | |
| ENSG00000010295 | IFFO1 | |
| ENSG00000010310 | GIPR | |
| ENSG00000010327 | STAB1 | |
| ENSG00000010379 | SLC6A13 | |
| ENSG00000010404 | IDS | |
| ENSG00000010610 | CD4 | |
| ENSG00000010626 | LRRC23 | |

| | |
|---|---|
| ENSG00000010810 | FYN |
| ENSG00000010818 | HIVEP2 |
| ENSG00000010932 | FMO1 |
| ENSG00000011028 | MRC2 |
| ENSG00000011083 | SLC6A7 |
| ENSG00000011105 | TSPAN9 |
| ENSG00000011332 | DPF1 |
| ENSG00000011451 | WIZ |
| ENSG00000012061 | ERCC1 |
| ENSG00000012232 | EXTL3 |
| ENSG00000013364 | MVP |
| ENSG00000013523 | ANGEL1 |
| ENSG00000013561 | RNF14 |
| ENSG00000015413 | DPEP1 |
| ENSG00000017260 | ATP2C1 |
| ENSG00000018408 | WWTR1 |
| ENSG00000018625 | ATP1A2 |
| ENSG00000019102 | VSIG2 |
| ENSG00000019144 | PHLDB1 |
| ENSG00000019169 | MARCO |
| ENSG00000019485 | PRDM11 |
| ENSG00000020181 | ADGRA2 |
| ENSG00000020256 | ZFP64 |
| ENSG00000020633 | RUNX3 |
| ENSG00000021300 | PLEKHB1 |
| ENSG00000021645 | NRXN3 |
| ENSG00000022567 | SLC45A4 |
| ENSG00000023171 | GRAMD1B |
| ENSG00000023287 | RB1CC1 |
| ENSG00000023892 | DEF6 |
| ENSG00000023902 | PLEKHO1 |
| ENSG00000024422 | EHD2 |
| ENSG00000027075 | PRKCH |
| ENSG00000028116 | VRK2 |
| ENSG00000028137 | TNFRSF1B |
| ENSG00000028277 | POU2F2 |
| ENSG00000029534 | ANK1 |
| ENSG00000031003 | FAM13B |
| ENSG00000033011 | ALG1 |
| ENSG00000034677 | RNF19A |
| ENSG00000035403 | VCL |
| ENSG00000035664 | DAPK2 |
| ENSG00000036257 | CUL3 |
| ENSG00000036448 | MYOM2 |
| ENSG00000036672 | USP2 |
| ENSG00000039139 | DNAH5 |
| ENSG00000039523 | RIPOR1 |
| ENSG00000039600 | SOX30 |
| ENSG00000039650 | PNKP |
| ENSG00000040608 | RTN4R |

| | |
|---|---|
| ENSG00000041353 | RAB27B |
| ENSG00000041515 | MYO16 |
| ENSG00000042062 | RIPOR3 |
| ENSG00000042781 | USH2A |
| ENSG00000042832 | TG |
| ENSG00000043093 | DCUN1D1 |
| ENSG00000044115 | CTNNA1 |
| ENSG00000047056 | WDR37 |
| ENSG00000047346 | FAM214A |
| ENSG00000047617 | ANO2 |
| ENSG00000047662 | FAM184B |
| ENSG00000047936 | ROS1 |
| ENSG00000048342 | CC2D2A |
| ENSG00000048405 | ZNF800 |
| ENSG00000048471 | SNX29 |
| ENSG00000049283 | EPN3 |
| ENSG00000049449 | RCN1 |
| ENSG00000049618 | ARID1B |
| ENSG00000050730 | TNIP3 |
| ENSG00000052749 | RRP12 |
| ENSG00000053108 | FSTL4 |
| ENSG00000053254 | FOXN3 |
| ENSG00000053524 | MCF2L2 |
| ENSG00000053918 | KCNQ1 |
| ENSG00000054116 | TRAPPC3 |
| ENSG00000054611 | TBC1D22A |
| ENSG00000054938 | CHRDL2 |
| ENSG00000055070 | SZRD1 |
| ENSG00000055118 | KCNH2 |
| ENSG00000055163 | CYFIP2 |
| ENSG00000055208 | TAB2 |
| ENSG00000055955 | ITIH4 |
| ENSG00000057608 | GDI2 |
| ENSG00000057704 | TMCC3 |
| ENSG00000057935 | MTA3 |
| ENSG00000058091 | CDK14 |
| ENSG00000058335 | RASGRF1 |
| ENSG00000058404 | CAMK2B |
| ENSG00000058453 | CROCC |
| ENSG00000058668 | ATP2B4 |
| ENSG00000058866 | DGKG |
| ENSG00000059145 | UNKL |
| ENSG00000059728 | MXD1 |
| ENSG00000061273 | HDAC7 |
| ENSG00000061337 | LZTS1 |
| ENSG00000061938 | TNK2 |
| ENSG00000062038 | CDH3 |
| ENSG00000062282 | DGAT2 |
| ENSG00000062370 | ZNF112 |
| ENSG00000062598 | ELMO2 |

| | |
|---|---|
| ENSG00000063245 | EPN1 |
| ENSG00000063438 | AHRR |
| ENSG00000063978 | RNF4 |
| ENSG00000064012 | CASP8 |
| ENSG00000064300 | NGFR |
| ENSG00000064886 | CHI3L2 |
| ENSG00000065150 | IPO5 |
| ENSG00000065320 | NTN1 |
| ENSG00000065357 | DGKA |
| ENSG00000065534 | MYLK |
| ENSG00000065618 | COL17A1 |
| ENSG00000065717 | TLE2 |
| ENSG00000065809 | FAM107B |
| ENSG00000065882 | TBC1D1 |
| ENSG00000065989 | PDE4A |
| ENSG00000066248 | NGEF |
| ENSG00000066405 | CLDN18 |
| ENSG00000066629 | EML1 |
| ENSG00000066735 | KIF26A |
| ENSG00000067208 | EVI5 |
| ENSG00000067840 | PDZD4 |
| ENSG00000067842 | ATP2B3 |
| ENSG00000068028 | RASSF1 |
| ENSG00000068383 | INPP5A |
| ENSG00000068615 | REEP1 |
| ENSG00000068724 | TTC7A |
| ENSG00000068745 | IP6K2 |
| ENSG00000068971 | PPP2R5B |
| ENSG00000069020 | MAST4 |
| ENSG00000069424 | KCNAB2 |
| ENSG00000069667 | RORA |
| ENSG00000069702 | TGFBR3 |
| ENSG00000069956 | MAPK6 |
| ENSG00000069966 | GNB5 |
| ENSG00000069974 | RAB27A |
| ENSG00000069998 | HDHD5 |
| ENSG00000070081 | NUCB2 |
| ENSG00000070087 | PFN2 |
| ENSG00000070182 | SPTB |
| ENSG00000070269 | TMEM260 |
| ENSG00000070526 | ST6GALNAC1 |
| ENSG00000070729 | CNGB1 |
| ENSG00000070808 | CAMK2A |
| ENSG00000070915 | SLC12A3 |
| ENSG00000071073 | MGAT4A |
| ENSG00000071242 | RPS6KA2 |
| ENSG00000071909 | MYO3B |
| ENSG00000072134 | EPN2 |
| ENSG00000072135 | PTPN18 |
| ENSG00000072163 | LIMS2 |

| | |
|---|---|
| ENSG00000072182 | ASIC4 |
| ENSG00000072195 | SPEG |
| ENSG00000072201 | LNX1 |
| ENSG00000072422 | RHOBTB1 |
| ENSG00000072609 | CHFR |
| ENSG00000072682 | P4HA2 |
| ENSG00000072864 | NDE1 |
| ENSG00000072952 | MRVI1 |
| ENSG00000073060 | SCARB1 |
| ENSG00000073146 | MOV10L1 |
| ENSG00000073350 | LLGL2 |
| ENSG00000073605 | GSDMB |
| ENSG00000073670 | ADAM11 |
| ENSG00000073754 | CD5L |
| ENSG00000074047 | GLI2 |
| ENSG00000074219 | TEAD2 |
| ENSG00000074416 | MGLL |
| ENSG00000074964 | ARHGEF10L |
| ENSG00000074966 | TXK |
| ENSG00000075073 | TACR2 |
| ENSG00000075131 | TIPIN |
| ENSG00000075142 | SRI |
| ENSG00000075213 | SEMA3A |
| ENSG00000075275 | CELSR1 |
| ENSG00000075290 | WNT8B |
| ENSG00000075292 | ZNF638 |
| ENSG00000075618 | FSCN1 |
| ENSG00000075624 | ACTB |
| ENSG00000075891 | PAX2 |
| ENSG00000075945 | KIFAP3 |
| ENSG00000076356 | PLXNA2 |
| ENSG00000076554 | TPD52 |
| ENSG00000076555 | ACACB |
| ENSG00000076706 | MCAM |
| ENSG00000077063 | CTTNBP2 |
| ENSG00000077092 | RARB |
| ENSG00000077514 | POLD3 |
| ENSG00000077942 | FBLN1 |
| ENSG00000078070 | MCCC1 |
| ENSG00000078237 | TIGAR |
| ENSG00000078246 | TULP3 |
| ENSG00000078399 | HOXA9 |
| ENSG00000078687 | TNRC6C |
| ENSG00000078804 | TP53INP2 |
| ENSG00000079112 | CDH17 |
| ENSG00000079308 | TNS1 |
| ENSG00000079335 | CDC14A |
| ENSG00000079337 | RAPGEF3 |
| ENSG00000079385 | CEACAM1 |
| ENSG00000079393 | DUSP13 |

| | |
|---|---|
| ENSG00000079435 | LIPE |
| ENSG00000079459 | FDFT1 |
| ENSG00000080503 | SMARCA2 |
| ENSG00000080824 | HSP90AA1 |
| ENSG00000080845 | DLGAP4 |
| ENSG00000080854 | IGSF9B |
| ENSG00000081087 | OSTM1 |
| ENSG00000081189 | MEF2C |
| ENSG00000081248 | CACNA1S |
| ENSG00000081320 | STK17B |
| ENSG00000081377 | CDC14B |
| ENSG00000082014 | SMARCD3 |
| ENSG00000082074 | FYB1 |
| ENSG00000082458 | DLG3 |
| ENSG00000082781 | ITGB5 |
| ENSG00000083457 | ITGAE |
| ENSG00000083807 | SLC27A5 |
| ENSG00000083828 | ZNF586 |
| ENSG00000084070 | SMAP2 |
| ENSG00000084072 | PPIE |
| ENSG00000084628 | NKAIN1 |
| ENSG00000084693 | AGBL5 |
| ENSG00000084710 | EFR3B |
| ENSG00000085449 | WDFY1 |
| ENSG00000085465 | OVGP1 |
| ENSG00000085563 | ABCB1 |
| ENSG00000085644 | ZNF213 |
| ENSG00000085788 | DDHD2 |
| ENSG00000085831 | TTC39A |
| ENSG00000085978 | ATG16L1 |
| ENSG00000085998 | POMGNT1 |
| ENSG00000086015 | MAST2 |
| ENSG00000086200 | IPO11 |
| ENSG00000086289 | EPDR1 |
| ENSG00000086570 | FAT2 |
| ENSG00000086730 | LAT2 |
| ENSG00000086967 | MYBPC2 |
| ENSG00000086991 | NOX4 |
| ENSG00000087008 | ACOX3 |
| ENSG00000087237 | CETP |
| ENSG00000087258 | GNAO1 |
| ENSG00000087303 | NID2 |
| ENSG00000087460 | GNAS |
| ENSG00000087495 | PHACTR3 |
| ENSG00000087903 | RFX2 |
| ENSG00000088002 | SULT2B1 |
| ENSG00000088367 | EPB41L1 |
| ENSG00000088726 | TMEM40 |
| ENSG00000088756 | ARHGAP28 |
| ENSG00000088826 | SMOX |

| | |
|---|---|
| ENSG00000088833 | NSFL1C |
| ENSG00000088876 | ZNF343 |
| ENSG00000088881 | EBF4 |
| ENSG00000088992 | TESC |
| ENSG00000089053 | ANAPC5 |
| ENSG00000089057 | SLC23A2 |
| ENSG00000089060 | SLC8B1 |
| ENSG00000089101 | CFAP61 |
| ENSG00000089159 | PXN |
| ENSG00000089351 | GRAMD1A |
| ENSG00000089356 | FXYD3 |
| ENSG00000089692 | LAG3 |
| ENSG00000089693 | MLF2 |
| ENSG00000089818 | NECAP1 |
| ENSG00000089820 | ARHGAP4 |
| ENSG00000089847 | ANKRD24 |
| ENSG00000090006 | LTBP4 |
| ENSG00000090020 | SLC9A1 |
| ENSG00000090097 | PCBP4 |
| ENSG00000090266 | NDUFB2 |
| ENSG00000090512 | FETUB |
| ENSG00000090905 | TNRC6A |
| ENSG00000090975 | PITPNM2 |
| ENSG00000091128 | LAMB4 |
| ENSG00000091428 | RAPGEF4 |
| ENSG00000091482 | SMPX |
| ENSG00000091513 | TF |
| ENSG00000091536 | MYO15A |
| ENSG00000091583 | APOH |
| ENSG00000091947 | TMEM101 |
| ENSG00000091986 | CCDC80 |
| ENSG00000092200 | RPGRIP1 |
| ENSG00000092295 | TGM1 |
| ENSG00000092529 | CAPN3 |
| ENSG00000092607 | TBX15 |
| ENSG00000092621 | PHGDH |
| ENSG00000092758 | COL9A3 |
| ENSG00000092964 | DPYSL2 |
| ENSG00000093010 | COMT |
| ENSG00000093167 | LRRFIP2 |
| ENSG00000093183 | SEC22C |
| ENSG00000095066 | HOOK2 |
| ENSG00000095303 | PTGS1 |
| ENSG00000095370 | SH2D3C |
| ENSG00000095397 | WHRN |
| ENSG00000095585 | BLNK |
| ENSG00000095587 | TLL2 |
| ENSG00000095932 | SMIM24 |
| ENSG00000096060 | FKBP5 |
| ENSG00000096088 | PGC |

| | |
|---|---|
| ENSG00000099204 | ABLIM1 |
| ENSG00000099284 | H2AFY2 |
| ENSG00000099308 | MAST3 |
| ENSG00000099622 | CIRBP |
| ENSG00000099797 | TECR |
| ENSG00000099864 | PALM |
| ENSG00000099940 | SNAP29 |
| ENSG00000099953 | MMP11 |
| ENSG00000099954 | CECR2 |
| ENSG00000099957 | P2RX6 |
| ENSG00000099968 | BCL2L13 |
| ENSG00000099994 | SUSD2 |
| ENSG00000099999 | RNF215 |
| ENSG00000100031 | GGT1 |
| ENSG00000100033 | PRODH |
| ENSG00000100034 | PPM1F |
| ENSG00000100092 | SH3BP1 |
| ENSG00000100106 | TRIOBP |
| ENSG00000100154 | TTC28 |
| ENSG00000100170 | SLC5A1 |
| ENSG00000100207 | TCF20 |
| ENSG00000100218 | RSPH14 |
| ENSG00000100228 | RAB36 |
| ENSG00000100242 | SUN2 |
| ENSG00000100276 | RASL10A |
| ENSG00000100280 | AP1B1 |
| ENSG00000100307 | CBX7 |
| ENSG00000100346 | CACNA1I |
| ENSG00000100385 | IL2RB |
| ENSG00000100416 | TRMU |
| ENSG00000100433 | KCNK10 |
| ENSG00000100490 | CDKL1 |
| ENSG00000100612 | DHRS7 |
| ENSG00000100628 | ASB2 |
| ENSG00000100629 | CEP128 |
| ENSG00000100650 | SRSF5 |
| ENSG00000100802 | C14orf93 |
| ENSG00000100902 | PSMA6 |
| ENSG00000100934 | SEC23A |
| ENSG00000101000 | PROCR |
| ENSG00000101004 | NINL |
| ENSG00000101096 | NFATC2 |
| ENSG00000101098 | RIMS4 |
| ENSG00000101144 | BMP7 |
| ENSG00000101188 | NTSR1 |
| ENSG00000101198 | NKAIN4 |
| ENSG00000101203 | COL20A1 |
| ENSG00000101204 | CHRNA4 |
| ENSG00000101224 | CDC25B |
| ENSG00000101255 | TRIB3 |

| | |
|---|---|
| ENSG00000101276 | SLC52A3 |
| ENSG00000101294 | HM13 |
| ENSG00000101306 | MYLK2 |
| ENSG00000101331 | CCM2L |
| ENSG00000101342 | TLDC2 |
| ENSG00000101464 | PIGU |
| ENSG00000101489 | CELF4 |
| ENSG00000101544 | ADNP2 |
| ENSG00000101605 | MYOM1 |
| ENSG00000101680 | LAMA1 |
| ENSG00000101773 | RBBP8 |
| ENSG00000101871 | MID1 |
| ENSG00000101901 | ALG13 |
| ENSG00000101935 | AMMECR1 |
| ENSG00000101940 | WDR13 |
| ENSG00000101974 | ATP11C |
| ENSG00000102003 | SYP |
| ENSG00000102057 | KCND1 |
| ENSG00000102144 | PGK1 |
| ENSG00000102174 | PHEX |
| ENSG00000102359 | SRPX2 |
| ENSG00000102445 | RUBCNL |
| ENSG00000102554 | KLF5 |
| ENSG00000102606 | ARHGEF7 |
| ENSG00000102755 | FLT1 |
| ENSG00000102780 | DGKH |
| ENSG00000102901 | CENPT |
| ENSG00000102921 | N4BP1 |
| ENSG00000102962 | CCL22 |
| ENSG00000102984 | ZNF821 |
| ENSG00000103021 | CCDC113 |
| ENSG00000103034 | NDRG4 |
| ENSG00000103043 | VAC14 |
| ENSG00000103056 | SMPD3 |
| ENSG00000103091 | WDR59 |
| ENSG00000103196 | CRISPLD2 |
| ENSG00000103227 | LMF1 |
| ENSG00000103264 | FBXO31 |
| ENSG00000103365 | GGA2 |
| ENSG00000103375 | AQP8 |
| ENSG00000103544 | C16orf62 |
| ENSG00000103549 | RNF40 |
| ENSG00000103647 | CORO2B |
| ENSG00000103710 | RASL12 |
| ENSG00000103740 | ACSBG1 |
| ENSG00000103769 | RAB11A |
| ENSG00000103811 | CTSH |
| ENSG00000104044 | OCA2 |
| ENSG00000104059 | FAM189A1 |
| ENSG00000104067 | TJP1 |

| | |
|---|---|
| ENSG00000104368 | PLAT |
| ENSG00000104381 | GDAP1 |
| ENSG00000104408 | EIF3E |
| ENSG00000104415 | WISP1 |
| ENSG00000104419 | NDRG1 |
| ENSG00000104447 | TRPS1 |
| ENSG00000104490 | NCALD |
| ENSG00000104626 | ERI1 |
| ENSG00000104691 | UBXN8 |
| ENSG00000104714 | ERICH1 |
| ENSG00000104728 | ARHGEF10 |
| ENSG00000104765 | BNIP3L |
| ENSG00000104866 | PPP1R37 |
| ENSG00000104870 | FCGRT |
| ENSG00000104883 | PEX11G |
| ENSG00000104889 | RNASEH2A |
| ENSG00000104892 | KLC3 |
| ENSG00000104936 | DMPK |
| ENSG00000104941 | RSPH6A |
| ENSG00000104957 | CCDC130 |
| ENSG00000104983 | CCDC61 |
| ENSG00000105088 | OLFM2 |
| ENSG00000105176 | URI1 |
| ENSG00000105220 | GPI |
| ENSG00000105227 | PRX |
| ENSG00000105327 | BBC3 |
| ENSG00000105339 | DENND3 |
| ENSG00000105357 | MYH14 |
| ENSG00000105401 | CDC37 |
| ENSG00000105409 | ATP1A3 |
| ENSG00000105464 | GRIN2D |
| ENSG00000105514 | RAB3D |
| ENSG00000105523 | FAM83E |
| ENSG00000105538 | RASIP1 |
| ENSG00000105552 | BCAT2 |
| ENSG00000105613 | MAST1 |
| ENSG00000105642 | KCNN1 |
| ENSG00000105650 | PDE4C |
| ENSG00000105668 | UPK1A |
| ENSG00000105696 | TMEM59L |
| ENSG00000105732 | ZNF574 |
| ENSG00000105793 | GTPBP10 |
| ENSG00000105889 | STEAP1B |
| ENSG00000105929 | ATP6V0A4 |
| ENSG00000105963 | ADAP1 |
| ENSG00000105971 | CAV2 |
| ENSG00000105997 | HOXA3 |
| ENSG00000106003 | LFNG |
| ENSG00000106066 | CPVL |
| ENSG00000106069 | CHN2 |

| | |
|---|---|
| ENSG00000106125 | MINDY4 |
| ENSG00000106302 | HYAL4 |
| ENSG00000106327 | TFR2 |
| ENSG00000106355 | LSM5 |
| ENSG00000106366 | SERPINE1 |
| ENSG00000106392 | C1GALT1 |
| ENSG00000106404 | CLDN15 |
| ENSG00000106541 | AGR2 |
| ENSG00000106617 | PRKAG2 |
| ENSG00000106689 | LHX2 |
| ENSG00000106852 | LHX6 |
| ENSG00000106948 | AKNA |
| ENSG00000106976 | DNM1 |
| ENSG00000106991 | ENG |
| ENSG00000107077 | KDM4C |
| ENSG00000107104 | KANK1 |
| ENSG00000107130 | NCS1 |
| ENSG00000107185 | RGP1 |
| ENSG00000107187 | LHX3 |
| ENSG00000107249 | GLIS3 |
| ENSG00000107263 | RAPGEF1 |
| ENSG00000107371 | EXOSC3 |
| ENSG00000107551 | RASSF4 |
| ENSG00000107593 | PKD2L1 |
| ENSG00000107719 | PALD1 |
| ENSG00000107731 | UNC5B |
| ENSG00000107736 | CDH23 |
| ENSG00000107738 | VSIR |
| ENSG00000107742 | SPOCK2 |
| ENSG00000107798 | LIPA |
| ENSG00000107821 | KAZALD1 |
| ENSG00000107859 | PITX3 |
| ENSG00000107902 | LHPP |
| ENSG00000107951 | MTPAP |
| ENSG00000107954 | NEURL1 |
| ENSG00000107957 | SH3PXD2A |
| ENSG00000108100 | CCNY |
| ENSG00000108176 | DNAJC12 |
| ENSG00000108262 | GIT1 |
| ENSG00000108370 | RGS9 |
| ENSG00000108379 | WNT3 |
| ENSG00000108387 | sept-04 |
| ENSG00000108389 | MTMR4 |
| ENSG00000108465 | CDK5RAP3 |
| ENSG00000108511 | HOXB6 |
| ENSG00000108557 | RAI1 |
| ENSG00000108561 | C1QBP |
| ENSG00000108576 | SLC6A4 |
| ENSG00000108641 | B9D1 |
| ENSG00000108666 | C17orf75 |

| | |
|---|---|
| ENSG00000108679 | LGALS3BP |
| ENSG00000108786 | HSD17B1 |
| ENSG00000108823 | SGCA |
| ENSG00000108846 | ABCC3 |
| ENSG00000108852 | MPP2 |
| ENSG00000108878 | CACNG1 |
| ENSG00000109016 | DHRS7B |
| ENSG00000109101 | FOXN1 |
| ENSG00000109180 | OCIAD1 |
| ENSG00000109332 | UBE2D3 |
| ENSG00000109339 | MAPK10 |
| ENSG00000109390 | NDUFC1 |
| ENSG00000109472 | CPE |
| ENSG00000109572 | CLCN3 |
| ENSG00000109610 | SOD3 |
| ENSG00000109667 | SLC2A9 |
| ENSG00000109680 | TBC1D19 |
| ENSG00000109743 | BST1 |
| ENSG00000109846 | CRYAB |
| ENSG00000109906 | ZBTB16 |
| ENSG00000110013 | SIAE |
| ENSG00000110047 | EHD1 |
| ENSG00000110076 | NRXN2 |
| ENSG00000110274 | CEP164 |
| ENSG00000110375 | UPK2 |
| ENSG00000110660 | SLC35F2 |
| ENSG00000110693 | SOX6 |
| ENSG00000110696 | C11orf58 |
| ENSG00000110777 | POU2AF1 |
| ENSG00000110799 | VWF |
| ENSG00000110811 | P3H3 |
| ENSG00000110844 | PRPF40B |
| ENSG00000110871 | COQ5 |
| ENSG00000110876 | SELPLG |
| ENSG00000110934 | BIN2 |
| ENSG00000110987 | BCL7A |
| ENSG00000111144 | LTA4H |
| ENSG00000111145 | ELK3 |
| ENSG00000111181 | SLC6A12 |
| ENSG00000111186 | WNT5B |
| ENSG00000111319 | SCNN1A |
| ENSG00000111321 | LTBR |
| ENSG00000111405 | ENDOU |
| ENSG00000111424 | VDR |
| ENSG00000111452 | ADGRD1 |
| ENSG00000111664 | GNB3 |
| ENSG00000111671 | SPSB2 |
| ENSG00000111676 | ATN1 |
| ENSG00000111716 | LDHB |
| ENSG00000111846 | GCNT2 |

| | |
|---|---|
| ENSG00000111859 | NEDD9 |
| ENSG00000111860 | CEP85L |
| ENSG00000111863 | ADTRP |
| ENSG00000111879 | FAM184A |
| ENSG00000111886 | GABRR2 |
| ENSG00000111907 | TPD52L1 |
| ENSG00000111912 | NCOA7 |
| ENSG00000111913 | RIPOR2 |
| ENSG00000111961 | SASH1 |
| ENSG00000112033 | PPARD |
| ENSG00000112053 | SLC26A8 |
| ENSG00000112096 | SOD2 |
| ENSG00000112137 | PHACTR1 |
| ENSG00000112146 | FBXO9 |
| ENSG00000112182 | BACH2 |
| ENSG00000112245 | PTP4A1 |
| ENSG00000112297 | CRYBG1 |
| ENSG00000112339 | HBS1L |
| ENSG00000112486 | CCR6 |
| ENSG00000112499 | SLC22A2 |
| ENSG00000112541 | PDE10A |
| ENSG00000112562 | SMOC2 |
| ENSG00000112576 | CCND3 |
| ENSG00000112584 | FAM120B |
| ENSG00000112695 | COX7A2 |
| ENSG00000112699 | GMDS |
| ENSG00000112782 | CLIC5 |
| ENSG00000112787 | FBRSL1 |
| ENSG00000112812 | PRSS16 |
| ENSG00000112981 | NME5 |
| ENSG00000113231 | PDE8B |
| ENSG00000113240 | CLK4 |
| ENSG00000113387 | SUB1 |
| ENSG00000113389 | NPR3 |
| ENSG00000113391 | FAM172A |
| ENSG00000113448 | PDE4D |
| ENSG00000113504 | SLC12A7 |
| ENSG00000113580 | NR3C1 |
| ENSG00000113594 | LIFR |
| ENSG00000113645 | WWC1 |
| ENSG00000113657 | DPYSL3 |
| ENSG00000113721 | PDGFRB |
| ENSG00000113722 | CDX1 |
| ENSG00000113749 | HRH2 |
| ENSG00000113758 | DBN1 |
| ENSG00000113763 | UNC5A |
| ENSG00000113790 | EHHADH |
| ENSG00000113889 | KNG1 |
| ENSG00000114115 | RBP1 |
| ENSG00000114124 | GRK7 |

| | |
|---|---|
| ENSG00000114268 | PFKFB4 |
| ENSG00000114349 | GNAT1 |
| ENSG00000114378 | HYAL1 |
| ENSG00000114416 | FXR1 |
| ENSG00000114541 | FRMD4B |
| ENSG00000114544 | SLC41A3 |
| ENSG00000114654 | EFCC1 |
| ENSG00000114735 | HEMK1 |
| ENSG00000114738 | MAPKAPK3 |
| ENSG00000114812 | VIPR1 |
| ENSG00000114982 | KANSL3 |
| ENSG00000115041 | KCNIP3 |
| ENSG00000115053 | NCL |
| ENSG00000115155 | OTOF |
| ENSG00000115165 | CYTIP |
| ENSG00000115194 | SLC30A3 |
| ENSG00000115239 | ASB3 |
| ENSG00000115266 | APC2 |
| ENSG00000115310 | RTN4 |
| ENSG00000115457 | IGFBP2 |
| ENSG00000115504 | EHBP1 |
| ENSG00000115525 | ST3GAL5 |
| ENSG00000115556 | PLCD4 |
| ENSG00000115590 | IL1R2 |
| ENSG00000115594 | IL1R1 |
| ENSG00000115604 | IL18R1 |
| ENSG00000115641 | FHL2 |
| ENSG00000115648 | MLPH |
| ENSG00000115850 | LCT |
| ENSG00000115944 | COX7A2L |
| ENSG00000115963 | RND3 |
| ENSG00000115977 | AAK1 |
| ENSG00000115998 | C2orf42 |
| ENSG00000116016 | EPAS1 |
| ENSG00000116039 | ATP6V1B1 |
| ENSG00000116044 | NFE2L2 |
| ENSG00000116062 | MSH6 |
| ENSG00000116128 | BCL9 |
| ENSG00000116183 | PAPPA2 |
| ENSG00000116251 | RPL22 |
| ENSG00000116254 | CHD5 |
| ENSG00000116299 | KIAA1324 |
| ENSG00000116473 | RAP1A |
| ENSG00000116584 | ARHGEF2 |
| ENSG00000116670 | MAD2L2 |
| ENSG00000116675 | DNAJC6 |
| ENSG00000116701 | NCF2 |
| ENSG00000116833 | NR5A2 |
| ENSG00000116852 | KIF21B |
| ENSG00000116857 | TMEM9 |

| | |
|---|---|
| ENSG00000116977 | LGALS8 |
| ENSG00000116991 | SIPA1L2 |
| ENSG00000117115 | PADI2 |
| ENSG00000117245 | KIF17 |
| ENSG00000117298 | ECE1 |
| ENSG00000117305 | HMGCL |
| ENSG00000117400 | MPL |
| ENSG00000117425 | PTCH2 |
| ENSG00000117472 | TSPAN1 |
| ENSG00000117616 | RSRP1 |
| ENSG00000117632 | STMN1 |
| ENSG00000117643 | MAN1C1 |
| ENSG00000117859 | OSBPL9 |
| ENSG00000117971 | CHRNB4 |
| ENSG00000117983 | MUC5B |
| ENSG00000118004 | COLEC11 |
| ENSG00000118046 | STK11 |
| ENSG00000118094 | TREH |
| ENSG00000118160 | SLC8A2 |
| ENSG00000118242 | MREG |
| ENSG00000118257 | NRP2 |
| ENSG00000118308 | LRMP |
| ENSG00000118515 | SGK1 |
| ENSG00000118557 | PMFBP1 |
| ENSG00000118564 | FBXL5 |
| ENSG00000118690 | ARMC2 |
| ENSG00000118777 | ABCG2 |
| ENSG00000118855 | MFSD1 |
| ENSG00000118898 | PPL |
| ENSG00000118922 | KLF12 |
| ENSG00000118960 | HS1BP3 |
| ENSG00000119042 | SATB2 |
| ENSG00000119125 | GDA |
| ENSG00000119283 | TRIM67 |
| ENSG00000119396 | RAB14 |
| ENSG00000119403 | PHF19 |
| ENSG00000119574 | ZBTB45 |
| ENSG00000119630 | PGF |
| ENSG00000119650 | IFT43 |
| ENSG00000119681 | LTBP2 |
| ENSG00000119685 | TTLL5 |
| ENSG00000119698 | PPP4R4 |
| ENSG00000119771 | KLHL29 |
| ENSG00000119772 | DNMT3A |
| ENSG00000119888 | EPCAM |
| ENSG00000119913 | TECTB |
| ENSG00000119917 | IFIT3 |
| ENSG00000119922 | IFIT2 |
| ENSG00000119927 | GPAM |
| ENSG00000119943 | PYROXD2 |

| | |
|---|---|
| ENSG00000119946 | CNNM1 |
| ENSG00000119950 | MXI1 |
| ENSG00000119969 | HELLS |
| ENSG00000120049 | KCNIP2 |
| ENSG00000120071 | KANSL1 |
| ENSG00000120093 | HOXB3 |
| ENSG00000120332 | TNN |
| ENSG00000120341 | SEC16B |
| ENSG00000120440 | TTLL2 |
| ENSG00000120549 | KIAA1217 |
| ENSG00000120616 | EPC1 |
| ENSG00000120645 | IQSEC3 |
| ENSG00000120647 | CCDC77 |
| ENSG00000120693 | SMAD9 |
| ENSG00000120725 | SIL1 |
| ENSG00000120784 | ZFP30 |
| ENSG00000120896 | SORBS3 |
| ENSG00000120899 | PTK2B |
| ENSG00000120903 | CHRNA2 |
| ENSG00000120913 | PDLIM2 |
| ENSG00000121075 | TBX4 |
| ENSG00000121270 | ABCC11 |
| ENSG00000121380 | BCL2L14 |
| ENSG00000121410 | A1BG |
| ENSG00000121454 | LHX4 |
| ENSG00000121577 | POPDC2 |
| ENSG00000121653 | MAPK8IP1 |
| ENSG00000121753 | ADGRB2 |
| ENSG00000121898 | CPXM2 |
| ENSG00000121933 | TMIGD3 |
| ENSG00000122012 | SV2C |
| ENSG00000122025 | FLT3 |
| ENSG00000122188 | LAX1 |
| ENSG00000122367 | LDB3 |
| ENSG00000122390 | NAA60 |
| ENSG00000122547 | EEPD1 |
| ENSG00000122711 | SPINK4 |
| ENSG00000122786 | CALD1 |
| ENSG00000122877 | EGR2 |
| ENSG00000123080 | CDKN2C |
| ENSG00000123096 | SSPN |
| ENSG00000123106 | CCDC91 |
| ENSG00000123130 | ACOT9 |
| ENSG00000123358 | NR4A1 |
| ENSG00000123384 | LRP1 |
| ENSG00000123415 | SMUG1 |
| ENSG00000123453 | SARDH |
| ENSG00000123472 | ATPAF1 |
| ENSG00000123643 | SLC36A1 |
| ENSG00000123992 | DNPEP |

| | |
|---|---|
| ENSG00000124003 | MOGAT1 |
| ENSG00000124006 | OBSL1 |
| ENSG00000124126 | PREX1 |
| ENSG00000124159 | MATN4 |
| ENSG00000124203 | ZNF831 |
| ENSG00000124205 | EDN3 |
| ENSG00000124212 | PTGIS |
| ENSG00000124215 | CDH26 |
| ENSG00000124225 | PMEPA1 |
| ENSG00000124313 | IQSEC2 |
| ENSG00000124343 | XG |
| ENSG00000124357 | NAGK |
| ENSG00000124440 | HIF3A |
| ENSG00000124496 | TRERF1 |
| ENSG00000124507 | PACSIN1 |
| ENSG00000124570 | SERPINB6 |
| ENSG00000124588 | NQO2 |
| ENSG00000124596 | OARD1 |
| ENSG00000124613 | ZNF391 |
| ENSG00000124743 | KLHL31 |
| ENSG00000124749 | COL21A1 |
| ENSG00000124772 | CPNE5 |
| ENSG00000124783 | SSR1 |
| ENSG00000124813 | RUNX2 |
| ENSG00000124831 | LRRFIP1 |
| ENSG00000124839 | RAB17 |
| ENSG00000124942 | AHNAK |
| ENSG00000125037 | EMC3 |
| ENSG00000125046 | SSUH2 |
| ENSG00000125089 | SH3TC1 |
| ENSG00000125246 | CLYBL |
| ENSG00000125337 | KIF25 |
| ENSG00000125462 | C1orf61 |
| ENSG00000125618 | PAX8 |
| ENSG00000125637 | PSD4 |
| ENSG00000125648 | SLC25A23 |
| ENSG00000125726 | CD70 |
| ENSG00000125730 | C3 |
| ENSG00000125735 | TNFSF14 |
| ENSG00000125741 | OPA3 |
| ENSG00000125744 | RTN2 |
| ENSG00000125746 | EML2 |
| ENSG00000125775 | SDCBP2 |
| ENSG00000125780 | TGM3 |
| ENSG00000125818 | PSMF1 |
| ENSG00000125821 | DTD1 |
| ENSG00000125864 | BFSP1 |
| ENSG00000125871 | MGME1 |
| ENSG00000125888 | BANF2 |
| ENSG00000125966 | MMP24 |

| | |
|---|---|
| ENSG00000126005 | MMP24-AS1 |
| ENSG00000126214 | KLC1 |
| ENSG00000126217 | MCF2L |
| ENSG00000126262 | FFAR2 |
| ENSG00000126351 | THRA |
| ENSG00000126583 | PRKCG |
| ENSG00000126705 | AHDC1 |
| ENSG00000126759 | CFP |
| ENSG00000126777 | KTN1 |
| ENSG00000126803 | HSPA2 |
| ENSG00000126856 | PRDM7 |
| ENSG00000127124 | HIVEP3 |
| ENSG00000127220 | ABHD8 |
| ENSG00000127241 | MASP1 |
| ENSG00000127325 | BEST3 |
| ENSG00000127329 | PTPRB |
| ENSG00000127415 | IDUA |
| ENSG00000127452 | FBXL12 |
| ENSG00000127585 | FBXL16 |
| ENSG00000127603 | MACF1 |
| ENSG00000127838 | PNKD |
| ENSG00000127863 | TNFRSF19 |
| ENSG00000127946 | HIP1 |
| ENSG00000127948 | POR |
| ENSG00000128011 | LRFN1 |
| ENSG00000128266 | GNAZ |
| ENSG00000128268 | MGAT3 |
| ENSG00000128284 | APOL3 |
| ENSG00000128294 | TPST2 |
| ENSG00000128487 | SPECC1 |
| ENSG00000128563 | PRKRIP1 |
| ENSG00000128683 | GAD1 |
| ENSG00000128805 | ARHGAP22 |
| ENSG00000128849 | CGNL1 |
| ENSG00000128918 | ALDH1A2 |
| ENSG00000128928 | IVD |
| ENSG00000128973 | CLN6 |
| ENSG00000129116 | PALLD |
| ENSG00000129159 | KCNC1 |
| ENSG00000129214 | SHBG |
| ENSG00000129244 | ATP1B2 |
| ENSG00000129353 | SLC44A2 |
| ENSG00000129521 | EGLN3 |
| ENSG00000129595 | EPB41L4A |
| ENSG00000129646 | QRICH2 |
| ENSG00000129657 | SEC14L1 |
| ENSG00000129673 | AANAT |
| ENSG00000129680 | MAP7D3 |
| ENSG00000129682 | FGF13 |
| ENSG00000130035 | GALNT8 |

| | |
|---|---|
| ENSG00000130038 | CRACR2A |
| ENSG00000130052 | STARD8 |
| ENSG00000130055 | GDPD2 |
| ENSG00000130255 | RPL36 |
| ENSG00000130287 | NCAN |
| ENSG00000130307 | USHBP1 |
| ENSG00000130312 | MRPL34 |
| ENSG00000130340 | SNX9 |
| ENSG00000130475 | FCHO1 |
| ENSG00000130477 | UNC13A |
| ENSG00000130513 | GDF15 |
| ENSG00000130528 | HRC |
| ENSG00000130561 | SAG |
| ENSG00000130584 | ZBTB46 |
| ENSG00000130643 | CALY |
| ENSG00000130700 | GATA5 |
| ENSG00000130701 | RBBP8NL |
| ENSG00000130703 | OSBPL2 |
| ENSG00000130707 | ASS1 |
| ENSG00000130711 | PRDM12 |
| ENSG00000130751 | NPAS1 |
| ENSG00000130812 | ANGPTL6 |
| ENSG00000130816 | DNMT1 |
| ENSG00000130830 | MPP1 |
| ENSG00000130844 | ZNF331 |
| ENSG00000130940 | CASZ1 |
| ENSG00000131016 | AKAP12 |
| ENSG00000131018 | SYNE1 |
| ENSG00000131037 | EPS8L1 |
| ENSG00000131044 | TTLL9 |
| ENSG00000131050 | BPIFA2 |
| ENSG00000131089 | ARHGEF9 |
| ENSG00000131115 | ZNF227 |
| ENSG00000131126 | TEX101 |
| ENSG00000131149 | GSE1 |
| ENSG00000131188 | PRR7 |
| ENSG00000131196 | NFATC1 |
| ENSG00000131242 | RAB11FIP4 |
| ENSG00000131370 | SH3BP5 |
| ENSG00000131374 | TBC1D5 |
| ENSG00000131378 | RFTN1 |
| ENSG00000131379 | C3orf20 |
| ENSG00000131398 | KCNC3 |
| ENSG00000131408 | NR1H2 |
| ENSG00000131409 | LRRC4B |
| ENSG00000131435 | PDLIM4 |
| ENSG00000131446 | MGAT1 |
| ENSG00000131459 | GFPT2 |
| ENSG00000131473 | ACLY |
| ENSG00000131477 | RAMP2 |

| | |
|---|---|
| ENSG00000131508 | UBE2D2 |
| ENSG00000131584 | ACAP3 |
| ENSG00000131620 | ANO1 |
| ENSG00000131759 | RARA |
| ENSG00000131781 | FMO5 |
| ENSG00000131848 | ZSCAN5A |
| ENSG00000131914 | LIN28A |
| ENSG00000131981 | LGALS3 |
| ENSG00000132002 | DNAJB1 |
| ENSG00000132003 | ZSWIM4 |
| ENSG00000132005 | RFX1 |
| ENSG00000132026 | RTBDN |
| ENSG00000132205 | EMILIN2 |
| ENSG00000132256 | TRIM5 |
| ENSG00000132359 | RAP1GAP2 |
| ENSG00000132405 | TBC1D14 |
| ENSG00000132436 | FIGNL1 |
| ENSG00000132437 | DDC |
| ENSG00000132510 | KDM6B |
| ENSG00000132518 | GUCY2D |
| ENSG00000132563 | REEP2 |
| ENSG00000132604 | TERF2 |
| ENSG00000132623 | ANKEF1 |
| ENSG00000132669 | RIN2 |
| ENSG00000132670 | PTPRA |
| ENSG00000132702 | HAPLN2 |
| ENSG00000132821 | VSTM2L |
| ENSG00000132874 | SLC14A2 |
| ENSG00000132915 | PDE6A |
| ENSG00000132932 | ATP8A2 |
| ENSG00000132938 | MTUS2 |
| ENSG00000132965 | ALOX5AP |
| ENSG00000133027 | PEMT |
| ENSG00000133048 | CHI3L1 |
| ENSG00000133121 | STARD13 |
| ENSG00000133131 | MORC4 |
| ENSG00000133135 | RNF128 |
| ENSG00000133142 | TCEAL4 |
| ENSG00000133195 | SLC39A11 |
| ENSG00000133226 | SRRM1 |
| ENSG00000133243 | BTBD2 |
| ENSG00000133256 | PDE6B |
| ENSG00000133328 | HRASLS2 |
| ENSG00000133424 | LARGE1 |
| ENSG00000133454 | MYO18B |
| ENSG00000133597 | ADCK2 |
| ENSG00000133657 | ATP13A3 |
| ENSG00000133687 | TMTC1 |
| ENSG00000133808 | MICALCL |
| ENSG00000133812 | SBF2 |

| | |
|---|---|
| ENSG00000133816 | MICAL2 |
| ENSG00000133818 | RRAS2 |
| ENSG00000133943 | C14orf159 |
| ENSG00000133961 | NUMB |
| ENSG00000133962 | CATSPERB |
| ENSG00000133980 | VRTN |
| ENSG00000134042 | MRO |
| ENSG00000134193 | REG4 |
| ENSG00000134201 | GSTM5 |
| ENSG00000134207 | SYT6 |
| ENSG00000134245 | WNT2B |
| ENSG00000134247 | PTGFRN |
| ENSG00000134318 | ROCK2 |
| ENSG00000134324 | LPIN1 |
| ENSG00000134369 | NAV1 |
| ENSG00000134504 | KCTD1 |
| ENSG00000134516 | DOCK2 |
| ENSG00000134686 | PHC2 |
| ENSG00000134769 | DTNA |
| ENSG00000134871 | COL4A2 |
| ENSG00000134954 | ETS1 |
| ENSG00000134955 | SLC37A2 |
| ENSG00000134986 | NREP |
| ENSG00000135048 | TMEM2 |
| ENSG00000135077 | HAVCR2 |
| ENSG00000135083 | CCNJL |
| ENSG00000135093 | USP30 |
| ENSG00000135114 | OASL |
| ENSG00000135127 | BICDL1 |
| ENSG00000135205 | CCDC146 |
| ENSG00000135218 | CD36 |
| ENSG00000135250 | SRPK2 |
| ENSG00000135253 | KCP |
| ENSG00000135362 | PRR5L |
| ENSG00000135373 | EHF |
| ENSG00000135406 | PRPH |
| ENSG00000135424 | ITGA7 |
| ENSG00000135439 | AGAP2 |
| ENSG00000135443 | KRT85 |
| ENSG00000135446 | CDK4 |
| ENSG00000135452 | TSPAN31 |
| ENSG00000135472 | FAIM2 |
| ENSG00000135549 | PKIB |
| ENSG00000135596 | MICAL1 |
| ENSG00000135631 | RAB11FIP5 |
| ENSG00000135636 | DYSF |
| ENSG00000135678 | CPM |
| ENSG00000135723 | FHOD1 |
| ENSG00000135740 | SLC9A5 |
| ENSG00000135744 | AGT |

| | |
|---|---|
| ENSG00000135766 | EGLN1 |
| ENSG00000135773 | CAPN9 |
| ENSG00000135835 | KIAA1614 |
| ENSG00000135845 | PIGC |
| ENSG00000135899 | SP110 |
| ENSG00000135919 | SERPINE2 |
| ENSG00000135925 | WNT10A |
| ENSG00000135931 | ARMC9 |
| ENSG00000135951 | TSGA10 |
| ENSG00000136014 | USP44 |
| ENSG00000136059 | VILL |
| ENSG00000136153 | LMO7 |
| ENSG00000136235 | GPNMB |
| ENSG00000136240 | KDELR2 |
| ENSG00000136250 | AOAH |
| ENSG00000136267 | DGKB |
| ENSG00000136273 | HUS1 |
| ENSG00000136286 | MYO1G |
| ENSG00000136379 | ABHD17C |
| ENSG00000136425 | CIB2 |
| ENSG00000136449 | MYCBPAP |
| ENSG00000136560 | TANK |
| ENSG00000136574 | GATA4 |
| ENSG00000136689 | IL1RN |
| ENSG00000136830 | FAM129B |
| ENSG00000136848 | DAB2IP |
| ENSG00000136854 | STXBP1 |
| ENSG00000136877 | FPGS |
| ENSG00000136895 | GARNL3 |
| ENSG00000136928 | GABBR2 |
| ENSG00000136935 | GOLGA1 |
| ENSG00000136944 | LMX1B |
| ENSG00000137038 | DMAC1 |
| ENSG00000137074 | APTX |
| ENSG00000137090 | DMRT1 |
| ENSG00000137135 | ARHGEF39 |
| ENSG00000137198 | GMPR |
| ENSG00000137203 | TFAP2A |
| ENSG00000137270 | GCM1 |
| ENSG00000137275 | RIPK1 |
| ENSG00000137411 | VARS2 |
| ENSG00000137463 | MGARP |
| ENSG00000137491 | SLCO2B1 |
| ENSG00000137504 | CREBZF |
| ENSG00000137507 | LRRC32 |
| ENSG00000137509 | PRCP |
| ENSG00000137672 | TRPC6 |
| ENSG00000137674 | MMP20 |
| ENSG00000137699 | TRIM29 |
| ENSG00000137709 | POU2F3 |

| | |
|---|---|
| ENSG00000137726 | FXYD6 |
| ENSG00000137767 | SQOR |
| ENSG00000137809 | ITGA11 |
| ENSG00000137815 | RTF1 |
| ENSG00000137819 | PAQR5 |
| ENSG00000137834 | SMAD6 |
| ENSG00000137841 | PLCB2 |
| ENSG00000137842 | TMEM62 |
| ENSG00000137843 | PAK6 |
| ENSG00000137857 | DUOX1 |
| ENSG00000137869 | CYP19A1 |
| ENSG00000137871 | ZNF280D |
| ENSG00000137877 | SPTBN5 |
| ENSG00000137960 | GIPC2 |
| ENSG00000137962 | ARHGAP29 |
| ENSG00000137968 | SLC44A5 |
| ENSG00000138018 | SELENOI |
| ENSG00000138030 | KHK |
| ENSG00000138061 | CYP1B1 |
| ENSG00000138079 | SLC3A1 |
| ENSG00000138080 | EMILIN1 |
| ENSG00000138100 | TRIM54 |
| ENSG00000138101 | DTNB |
| ENSG00000138119 | MYOF |
| ENSG00000138138 | ATAD1 |
| ENSG00000138152 | BTBD16 |
| ENSG00000138162 | TACC2 |
| ENSG00000138190 | EXOC6 |
| ENSG00000138315 | OIT3 |
| ENSG00000138316 | ADAMTS14 |
| ENSG00000138356 | AOX1 |
| ENSG00000138385 | SSB |
| ENSG00000138411 | HECW2 |
| ENSG00000138413 | IDH1 |
| ENSG00000138435 | CHRNA1 |
| ENSG00000138617 | PARP16 |
| ENSG00000138621 | PPCDC |
| ENSG00000138641 | HERC3 |
| ENSG00000138669 | PRKG2 |
| ENSG00000138670 | RASGEF1B |
| ENSG00000138696 | BMPR1B |
| ENSG00000138757 | G3BP2 |
| ENSG00000138760 | SCARB2 |
| ENSG00000138771 | SHROOM3 |
| ENSG00000138792 | ENPEP |
| ENSG00000138798 | EGF |
| ENSG00000138821 | SLC39A8 |
| ENSG00000138835 | RGS3 |
| ENSG00000139083 | ETV6 |
| ENSG00000139154 | AEBP2 |

| | |
|---|---|
| ENSG00000139160 | ETFBKMT |
| ENSG00000139174 | PRICKLE1 |
| ENSG00000139209 | SLC38A4 |
| ENSG00000139269 | INHBE |
| ENSG00000139364 | TMEM132B |
| ENSG00000139438 | FAM222A |
| ENSG00000139549 | DHH |
| ENSG00000139567 | ACVRL1 |
| ENSG00000139644 | TMBIM6 |
| ENSG00000139725 | RHOF |
| ENSG00000139865 | TTC6 |
| ENSG00000139890 | REM2 |
| ENSG00000139926 | FRMD6 |
| ENSG00000139946 | PELI2 |
| ENSG00000139988 | RDH12 |
| ENSG00000140057 | AK7 |
| ENSG00000140107 | SLC25A47 |
| ENSG00000140254 | DUOXA1 |
| ENSG00000140264 | SERF2 |
| ENSG00000140274 | DUOXA2 |
| ENSG00000140280 | LYSMD2 |
| ENSG00000140391 | TSPAN3 |
| ENSG00000140443 | IGF1R |
| ENSG00000140470 | ADAMTS17 |
| ENSG00000140479 | PCSK6 |
| ENSG00000140481 | CCDC33 |
| ENSG00000140564 | FURIN |
| ENSG00000140675 | SLC5A2 |
| ENSG00000140678 | ITGAX |
| ENSG00000140682 | TGFB1I1 |
| ENSG00000140743 | CDR2 |
| ENSG00000140795 | MYLK3 |
| ENSG00000140836 | ZFHX3 |
| ENSG00000140859 | KIFC3 |
| ENSG00000140939 | NOL3 |
| ENSG00000140941 | MAP1LC3B |
| ENSG00000140950 | TLDC1 |
| ENSG00000140961 | OSGIN1 |
| ENSG00000140992 | PDPK1 |
| ENSG00000141052 | MYOCD |
| ENSG00000141068 | KSR1 |
| ENSG00000141127 | PRPSAP2 |
| ENSG00000141252 | VPS53 |
| ENSG00000141279 | NPEPPS |
| ENSG00000141293 | SKAP1 |
| ENSG00000141314 | RHBDL3 |
| ENSG00000141316 | SPACA3 |
| ENSG00000141337 | ARSG |
| ENSG00000141391 | PRELID3A |
| ENSG00000141404 | GNAL |

| | |
|---|---|
| ENSG00000141485 | SLC13A5 |
| ENSG00000141524 | TMC6 |
| ENSG00000141526 | SLC16A3 |
| ENSG00000141527 | CARD14 |
| ENSG00000141556 | TBCD |
| ENSG00000141564 | RPTOR |
| ENSG00000141574 | SECTM1 |
| ENSG00000141639 | MAPK4 |
| ENSG00000141736 | ERBB2 |
| ENSG00000141837 | CACNA1A |
| ENSG00000141905 | NFIC |
| ENSG00000141971 | MVB12A |
| ENSG00000142046 | TMEM91 |
| ENSG00000142089 | IFITM3 |
| ENSG00000142185 | TRPM2 |
| ENSG00000142227 | EMP3 |
| ENSG00000142235 | LMTK3 |
| ENSG00000142319 | SLC6A3 |
| ENSG00000142347 | MYO1F |
| ENSG00000142459 | EVI5L |
| ENSG00000142552 | RCN3 |
| ENSG00000142583 | SLC2A5 |
| ENSG00000142599 | RERE |
| ENSG00000142609 | CFAP74 |
| ENSG00000142611 | PRDM16 |
| ENSG00000142619 | PADI3 |
| ENSG00000142621 | FHAD1 |
| ENSG00000142623 | PADI1 |
| ENSG00000142632 | ARHGEF19 |
| ENSG00000142661 | MYOM3 |
| ENSG00000142765 | SYTL1 |
| ENSG00000142949 | PTPRF |
| ENSG00000142973 | CYP4B1 |
| ENSG00000143126 | CELSR2 |
| ENSG00000143127 | ITGA10 |
| ENSG00000143167 | GPA33 |
| ENSG00000143183 | TMCO1 |
| ENSG00000143199 | ADCY10 |
| ENSG00000143217 | NECTIN4 |
| ENSG00000143321 | HDGF |
| ENSG00000143382 | ADAMTSL4 |
| ENSG00000143515 | ATP8B2 |
| ENSG00000143549 | TPM3 |
| ENSG00000143552 | NUP210L |
| ENSG00000143590 | EFNA3 |
| ENSG00000143603 | KCNN3 |
| ENSG00000143641 | GALNT2 |
| ENSG00000143669 | LYST |
| ENSG00000143753 | DEGS1 |
| ENSG00000143772 | ITPKB |

| | |
|---|---|
| ENSG00000143786 | CNIH3 |
| ENSG00000143850 | PLEKHA6 |
| ENSG00000143858 | SYT2 |
| ENSG00000143870 | PDIA6 |
| ENSG00000143882 | ATP6V1C2 |
| ENSG00000143951 | WDPCP |
| ENSG00000144029 | MRPS5 |
| ENSG00000144040 | SFXN5 |
| ENSG00000144118 | RALB |
| ENSG00000144130 | NT5DC4 |
| ENSG00000144229 | THSD7B |
| ENSG00000144407 | PTH2R |
| ENSG00000144488 | ESPNL |
| ENSG00000144560 | VGLL4 |
| ENSG00000144596 | GRIP2 |
| ENSG00000144648 | ACKR2 |
| ENSG00000144711 | IQSEC1 |
| ENSG00000144712 | CAND2 |
| ENSG00000144736 | SHQ1 |
| ENSG00000144741 | SLC25A26 |
| ENSG00000144791 | LIMD1 |
| ENSG00000144824 | PHLDB2 |
| ENSG00000144847 | IGSF11 |
| ENSG00000144852 | NR1I2 |
| ENSG00000144867 | SRPRB |
| ENSG00000145016 | RUBCN |
| ENSG00000145214 | DGKQ |
| ENSG00000145287 | PLAC8 |
| ENSG00000145332 | KLHL8 |
| ENSG00000145348 | TBCK |
| ENSG00000145362 | ANK2 |
| ENSG00000145391 | SETD7 |
| ENSG00000145642 | FAM159B |
| ENSG00000145685 | LHFPL2 |
| ENSG00000145901 | TNIP1 |
| ENSG00000145949 | MYLK4 |
| ENSG00000146090 | RASGEF1C |
| ENSG00000146122 | DAAM2 |
| ENSG00000146205 | ANO7 |
| ENSG00000146216 | TTBK1 |
| ENSG00000146285 | SCML4 |
| ENSG00000146426 | TIAM2 |
| ENSG00000146433 | TMEM181 |
| ENSG00000146535 | GNA12 |
| ENSG00000146592 | CREB5 |
| ENSG00000146648 | EGFR |
| ENSG00000146700 | SSC4D |
| ENSG00000146729 | NIPSNAP2 |
| ENSG00000146733 | PSPH |
| ENSG00000146828 | SLC12A9 |

| | |
|---|---|
| ENSG00000146872 | TLK2 |
| ENSG00000146966 | DENND2A |
| ENSG00000147133 | TAF1 |
| ENSG00000147164 | SNX12 |
| ENSG00000147394 | ZNF185 |
| ENSG00000147403 | RPL10 |
| ENSG00000147485 | PXDNL |
| ENSG00000147488 | ST18 |
| ENSG00000147509 | RGS20 |
| ENSG00000147526 | TACC1 |
| ENSG00000147642 | SYBU |
| ENSG00000147676 | MAL2 |
| ENSG00000147689 | FAM83A |
| ENSG00000147872 | PLIN2 |
| ENSG00000148180 | GSN |
| ENSG00000148204 | CRB2 |
| ENSG00000148219 | ASTN2 |
| ENSG00000148344 | PTGES |
| ENSG00000148357 | HMCN2 |
| ENSG00000148400 | NOTCH1 |
| ENSG00000148408 | CACNA1B |
| ENSG00000148671 | ADIRF |
| ENSG00000148700 | ADD3 |
| ENSG00000148702 | HABP2 |
| ENSG00000148795 | CYP17A1 |
| ENSG00000148814 | LRRC27 |
| ENSG00000148834 | GSTO1 |
| ENSG00000148848 | ADAM12 |
| ENSG00000148935 | GAS2 |
| ENSG00000148985 | PGAP2 |
| ENSG00000149016 | TUT1 |
| ENSG00000149090 | PAMR1 |
| ENSG00000149091 | DGKZ |
| ENSG00000149115 | TNKS1BP1 |
| ENSG00000149256 | TENM4 |
| ENSG00000149403 | GRIK4 |
| ENSG00000149485 | FADS1 |
| ENSG00000149527 | PLCH2 |
| ENSG00000149564 | ESAM |
| ENSG00000149575 | SCN2B |
| ENSG00000149596 | JPH2 |
| ENSG00000149599 | DUSP15 |
| ENSG00000149633 | KIAA1755 |
| ENSG00000149646 | CNBD2 |
| ENSG00000149823 | VPS51 |
| ENSG00000149925 | ALDOA |
| ENSG00000149932 | TMEM219 |
| ENSG00000149972 | CNTN5 |
| ENSG00000150054 | MPP7 |
| ENSG00000150093 | ITGB1 |

| | |
|---|---|
| ENSG00000150471 | ADGRL3 |
| ENSG00000150527 | CTAGE5 |
| ENSG00000150656 | CNDP1 |
| ENSG00000150687 | PRSS23 |
| ENSG00000150722 | PPP1R1C |
| ENSG00000150750 | C11orf53 |
| ENSG00000150764 | DIXDC1 |
| ENSG00000150967 | ABCB9 |
| ENSG00000151062 | CACNA2D4 |
| ENSG00000151065 | DCP1B |
| ENSG00000151067 | CACNA1C |
| ENSG00000151117 | TMEM86A |
| ENSG00000151240 | DIP2C |
| ENSG00000151320 | AKAP6 |
| ENSG00000151364 | KCTD14 |
| ENSG00000151388 | ADAMTS12 |
| ENSG00000151413 | NUBPL |
| ENSG00000151468 | CCDC3 |
| ENSG00000151474 | FRMD4A |
| ENSG00000151491 | EPS8 |
| ENSG00000151572 | ANO4 |
| ENSG00000151617 | EDNRA |
| ENSG00000151632 | AKR1C2 |
| ENSG00000151655 | ITIH2 |
| ENSG00000151726 | ACSL1 |
| ENSG00000151834 | GABRA2 |
| ENSG00000151835 | SACS |
| ENSG00000151882 | CCL28 |
| ENSG00000151914 | DST |
| ENSG00000152128 | TMEM163 |
| ENSG00000152253 | SPC25 |
| ENSG00000152284 | TCF7L1 |
| ENSG00000152503 | TRIM36 |
| ENSG00000152558 | TMEM123 |
| ENSG00000152601 | MBNL1 |
| ENSG00000152684 | PELO |
| ENSG00000152689 | RASGRP3 |
| ENSG00000152700 | SAR1B |
| ENSG00000152936 | LMNTD1 |
| ENSG00000153029 | MR1 |
| ENSG00000153046 | CDYL |
| ENSG00000153060 | TEKT5 |
| ENSG00000153064 | BANK1 |
| ENSG00000153093 | ACOXL |
| ENSG00000153113 | CAST |
| ENSG00000153130 | SCOC |
| ENSG00000153179 | RASSF3 |
| ENSG00000153233 | PTPRR |
| ENSG00000153234 | NR4A2 |
| ENSG00000153283 | CD96 |

| | |
|---|---|
| ENSG00000153294 | ADGRF4 |
| ENSG00000153303 | FRMD1 |
| ENSG00000153339 | TRAPPC8 |
| ENSG00000153531 | ADPRHL1 |
| ENSG00000153707 | PTPRD |
| ENSG00000153814 | JAZF1 |
| ENSG00000153902 | LGI4 |
| ENSG00000153914 | SREK1 |
| ENSG00000153930 | ANKFN1 |
| ENSG00000154016 | GRAP |
| ENSG00000154025 | SLC5A10 |
| ENSG00000154118 | JPH3 |
| ENSG00000154134 | ROBO3 |
| ENSG00000154146 | NRGN |
| ENSG00000154153 | RETREG1 |
| ENSG00000154227 | CERS3 |
| ENSG00000154262 | ABCA6 |
| ENSG00000154305 | MIA3 |
| ENSG00000154319 | FAM167A |
| ENSG00000154358 | OBSCN |
| ENSG00000154556 | SORBS2 |
| ENSG00000154678 | PDE1C |
| ENSG00000154710 | RABGEF1 |
| ENSG00000154783 | FGD5 |
| ENSG00000154928 | EPHB1 |
| ENSG00000155093 | PTPRN2 |
| ENSG00000155096 | AZIN1 |
| ENSG00000155265 | GOLGA7B |
| ENSG00000155324 | GRAMD2B |
| ENSG00000155465 | SLC7A7 |
| ENSG00000155506 | LARP1 |
| ENSG00000155545 | MIER3 |
| ENSG00000155629 | PIK3AP1 |
| ENSG00000155749 | ALS2CR12 |
| ENSG00000155754 | C2CD6 |
| ENSG00000155849 | ELMO1 |
| ENSG00000155893 | PXYLP1 |
| ENSG00000155926 | SLA |
| ENSG00000155959 | VBP1 |
| ENSG00000155980 | KIF5A |
| ENSG00000156011 | PSD3 |
| ENSG00000156030 | ELMSAN1 |
| ENSG00000156113 | KCNMA1 |
| ENSG00000156127 | BATF |
| ENSG00000156170 | NDUFAF6 |
| ENSG00000156206 | CFAP161 |
| ENSG00000156219 | ART3 |
| ENSG00000156222 | SLC28A1 |
| ENSG00000156273 | BACH1 |
| ENSG00000156411 | C14orf2 |

| | |
|---|---|
| ENSG00000156475 | PPP2R2B |
| ENSG00000156500 | FAM122C |
| ENSG00000156515 | HK1 |
| ENSG00000156574 | NODAL |
| ENSG00000156671 | SAMD8 |
| ENSG00000156711 | MAPK13 |
| ENSG00000156802 | ATAD2 |
| ENSG00000156853 | ZNF689 |
| ENSG00000157064 | NMNAT2 |
| ENSG00000157152 | SYN2 |
| ENSG00000157214 | STEAP2 |
| ENSG00000157216 | SSBP3 |
| ENSG00000157224 | CLDN12 |
| ENSG00000157349 | DDX19B |
| ENSG00000157350 | ST3GAL2 |
| ENSG00000157368 | IL34 |
| ENSG00000157388 | CACNA1D |
| ENSG00000157399 | ARSE |
| ENSG00000157429 | ZNF19 |
| ENSG00000157450 | RNF111 |
| ENSG00000157470 | FAM81A |
| ENSG00000157510 | AFAP1L1 |
| ENSG00000157538 | DSCR3 |
| ENSG00000157601 | MX1 |
| ENSG00000157625 | TAB3 |
| ENSG00000157657 | ZNF618 |
| ENSG00000157703 | SVOPL |
| ENSG00000157800 | SLC37A3 |
| ENSG00000157833 | GAREM2 |
| ENSG00000157851 | DPYSL5 |
| ENSG00000157856 | DRC1 |
| ENSG00000157890 | MEGF11 |
| ENSG00000158008 | EXTL1 |
| ENSG00000158019 | BABAM2 |
| ENSG00000158022 | TRIM63 |
| ENSG00000158023 | WDR66 |
| ENSG00000158062 | UBXN11 |
| ENSG00000158104 | HPD |
| ENSG00000158113 | LRRC43 |
| ENSG00000158169 | FANCC |
| ENSG00000158270 | COLEC12 |
| ENSG00000158290 | CUL4B |
| ENSG00000158321 | AUTS2 |
| ENSG00000158445 | KCNB1 |
| ENSG00000158458 | NRG2 |
| ENSG00000158525 | CPA5 |
| ENSG00000158552 | ZFAND2B |
| ENSG00000158555 | GDPD5 |
| ENSG00000158560 | DYNC1I1 |
| ENSG00000158683 | PKD1L1 |

| | |
|---|---|
| ENSG00000158691 | ZSCAN12 |
| ENSG00000158747 | NBL1 |
| ENSG00000158856 | DMTN |
| ENSG00000158955 | WNT9B |
| ENSG00000158985 | CDC42SE2 |
| ENSG00000159164 | SV2A |
| ENSG00000159173 | TNNI1 |
| ENSG00000159176 | CSRP1 |
| ENSG00000159200 | RCAN1 |
| ENSG00000159216 | RUNX1 |
| ENSG00000159261 | CLDN14 |
| ENSG00000159307 | SCUBE1 |
| ENSG00000159314 | ARHGAP27 |
| ENSG00000159337 | PLA2G4D |
| ENSG00000159403 | C1R |
| ENSG00000159450 | TCHH |
| ENSG00000159618 | ADGRG5 |
| ENSG00000159625 | DRC7 |
| ENSG00000159650 | UROC1 |
| ENSG00000159674 | SPON2 |
| ENSG00000159685 | CHCHD6 |
| ENSG00000159714 | ZDHHC1 |
| ENSG00000159733 | ZFYVE28 |
| ENSG00000159842 | ABR |
| ENSG00000159871 | LYPD5 |
| ENSG00000159899 | NPR2 |
| ENSG00000159921 | GNE |
| ENSG00000160111 | CPAMD8 |
| ENSG00000160145 | KALRN |
| ENSG00000160179 | ABCG1 |
| ENSG00000160183 | TMPRSS3 |
| ENSG00000160190 | SLC37A1 |
| ENSG00000160191 | PDE9A |
| ENSG00000160194 | NDUFV3 |
| ENSG00000160216 | AGPAT3 |
| ENSG00000160255 | ITGB2 |
| ENSG00000160323 | ADAMTS13 |
| ENSG00000160408 | ST6GALNAC6 |
| ENSG00000160447 | PKN3 |
| ENSG00000160460 | SPTBN4 |
| ENSG00000160593 | JAML |
| ENSG00000160746 | ANO10 |
| ENSG00000160789 | LMNA |
| ENSG00000160799 | CCDC12 |
| ENSG00000160808 | MYL3 |
| ENSG00000160838 | LRRC71 |
| ENSG00000160959 | LRRC14 |
| ENSG00000161010 | MRNIP |
| ENSG00000161031 | PGLYRP2 |
| ENSG00000161091 | MFSD12 |

| | |
|---|---|
| ENSG00000161217 | PCYT1A |
| ENSG00000161249 | DMKN |
| ENSG00000161267 | BDH1 |
| ENSG00000161270 | NPHS1 |
| ENSG00000161395 | PGAP3 |
| ENSG00000161509 | GRIN2C |
| ENSG00000161542 | PRPSAP1 |
| ENSG00000161544 | CYGB |
| ENSG00000161558 | TMEM143 |
| ENSG00000161609 | CCDC155 |
| ENSG00000161642 | ZNF385A |
| ENSG00000161800 | RACGAP1 |
| ENSG00000161813 | LARP4 |
| ENSG00000161888 | SPC24 |
| ENSG00000161914 | ZNF653 |
| ENSG00000162078 | ZG16B |
| ENSG00000162104 | ADCY9 |
| ENSG00000162148 | PPP1R32 |
| ENSG00000162341 | TPCN2 |
| ENSG00000162383 | SLC1A7 |
| ENSG00000162398 | LEXM |
| ENSG00000162413 | KLHL21 |
| ENSG00000162415 | ZSWIM5 |
| ENSG00000162438 | CTRC |
| ENSG00000162458 | FBLIM1 |
| ENSG00000162490 | DRAXIN |
| ENSG00000162510 | MATN1 |
| ENSG00000162511 | LAPTM5 |
| ENSG00000162576 | MXRA8 |
| ENSG00000162585 | FAAP20 |
| ENSG00000162599 | NFIA |
| ENSG00000162733 | DDR2 |
| ENSG00000162746 | FCRLB |
| ENSG00000162761 | LMX1A |
| ENSG00000162772 | ATF3 |
| ENSG00000162849 | KIF26B |
| ENSG00000162896 | PIGR |
| ENSG00000162909 | CAPN2 |
| ENSG00000162946 | DISC1 |
| ENSG00000162949 | CAPN13 |
| ENSG00000162972 | MAIP1 |
| ENSG00000163002 | NUP35 |
| ENSG00000163017 | ACTG2 |
| ENSG00000163050 | COQ8A |
| ENSG00000163072 | NOSTRIN |
| ENSG00000163131 | CTSS |
| ENSG00000163138 | PACRGL |
| ENSG00000163171 | CDC42EP3 |
| ENSG00000163218 | PGLYRP4 |
| ENSG00000163263 | C1orf189 |

| | |
|---|---|
| ENSG00000163288 | GABRB1 |
| ENSG00000163297 | ANTXR2 |
| ENSG00000163322 | ABRAXAS1 |
| ENSG00000163346 | PBXIP1 |
| ENSG00000163357 | DCST1 |
| ENSG00000163395 | IGFN1 |
| ENSG00000163406 | SLC15A2 |
| ENSG00000163412 | EIF4E3 |
| ENSG00000163430 | FSTL1 |
| ENSG00000163453 | IGFBP7 |
| ENSG00000163485 | ADORA1 |
| ENSG00000163520 | FBLN2 |
| ENSG00000163531 | NFASC |
| ENSG00000163535 | SGO2 |
| ENSG00000163635 | ATXN7 |
| ENSG00000163637 | PRICKLE2 |
| ENSG00000163704 | PRRT3 |
| ENSG00000163714 | U2SURP |
| ENSG00000163738 | MTHFD2L |
| ENSG00000163785 | RYK |
| ENSG00000163803 | PLB1 |
| ENSG00000163806 | SPDYA |
| ENSG00000163818 | LZTFL1 |
| ENSG00000163827 | LRRC2 |
| ENSG00000163866 | SMIM12 |
| ENSG00000163870 | TPRA1 |
| ENSG00000163885 | CFAP100 |
| ENSG00000163898 | LIPH |
| ENSG00000163904 | SENP2 |
| ENSG00000163923 | RPL39L |
| ENSG00000163932 | PRKCD |
| ENSG00000163959 | SLC51A |
| ENSG00000163975 | MELTF |
| ENSG00000164007 | CLDN19 |
| ENSG00000164038 | SLC9B2 |
| ENSG00000164066 | INTU |
| ENSG00000164081 | TEX264 |
| ENSG00000164093 | PITX2 |
| ENSG00000164096 | C4orf3 |
| ENSG00000164124 | TMEM144 |
| ENSG00000164175 | SLC45A2 |
| ENSG00000164253 | WDR41 |
| ENSG00000164292 | RHOBTB3 |
| ENSG00000164303 | ENPP6 |
| ENSG00000164309 | CMYA5 |
| ENSG00000164363 | SLC6A18 |
| ENSG00000164398 | ACSL6 |
| ENSG00000164402 | sept-08 |
| ENSG00000164465 | DCBLD1 |
| ENSG00000164488 | DACT2 |

| | |
|---|---|
| ENSG00000164512 | ANKRD55 |
| ENSG00000164530 | PI16 |
| ENSG00000164535 | DAGLB |
| ENSG00000164574 | GALNT10 |
| ENSG00000164638 | SLC29A4 |
| ENSG00000164674 | SYTL3 |
| ENSG00000164690 | SHH |
| ENSG00000164741 | DLC1 |
| ENSG00000164742 | ADCY1 |
| ENSG00000164808 | SPIDR |
| ENSG00000164830 | OXR1 |
| ENSG00000164855 | TMEM184A |
| ENSG00000164867 | NOS3 |
| ENSG00000164941 | INTS8 |
| ENSG00000164951 | PDP1 |
| ENSG00000164972 | C9orf24 |
| ENSG00000165124 | SVEP1 |
| ENSG00000165138 | ANKS6 |
| ENSG00000165140 | FBP1 |
| ENSG00000165238 | WNK2 |
| ENSG00000165424 | ZCCHC24 |
| ENSG00000165449 | SLC16A9 |
| ENSG00000165490 | DDIAS |
| ENSG00000165507 | C10orf10 |
| ENSG00000165548 | TMEM63C |
| ENSG00000165568 | AKR1E2 |
| ENSG00000165617 | DACT1 |
| ENSG00000165626 | BEND7 |
| ENSG00000165695 | AK8 |
| ENSG00000165702 | GFI1B |
| ENSG00000165752 | STK32C |
| ENSG00000165757 | JCAD |
| ENSG00000165775 | FUNDC2 |
| ENSG00000165810 | BTNL9 |
| ENSG00000165821 | SALL2 |
| ENSG00000165868 | HSPA12A |
| ENSG00000165914 | TTC7B |
| ENSG00000165917 | RAPSN |
| ENSG00000165929 | TC2N |
| ENSG00000165959 | CLMN |
| ENSG00000165973 | NELL1 |
| ENSG00000165995 | CACNB2 |
| ENSG00000166016 | ABTB2 |
| ENSG00000166025 | AMOTL1 |
| ENSG00000166033 | HTRA1 |
| ENSG00000166035 | LIPC |
| ENSG00000166111 | SVOP |
| ENSG00000166126 | AMN |
| ENSG00000166159 | LRTM2 |
| ENSG00000166220 | TBATA |

| | |
|---|---|
| ENSG00000166268 | MYRFL |
| ENSG00000166278 | C2 |
| ENSG00000166317 | SYNPO2L |
| ENSG00000166341 | DCHS1 |
| ENSG00000166359 | WDR88 |
| ENSG00000166391 | MOGAT2 |
| ENSG00000166394 | CYB5R2 |
| ENSG00000166402 | TUB |
| ENSG00000166411 | IDH3A |
| ENSG00000166415 | WDR72 |
| ENSG00000166444 | ST5 |
| ENSG00000166509 | CLEC3A |
| ENSG00000166532 | RIMKLB |
| ENSG00000166579 | NDEL1 |
| ENSG00000166669 | ATF7IP2 |
| ENSG00000166682 | TMPRSS5 |
| ENSG00000166743 | ACSM1 |
| ENSG00000166793 | YPEL4 |
| ENSG00000166819 | PLIN1 |
| ENSG00000166825 | ANPEP |
| ENSG00000166833 | NAV2 |
| ENSG00000166840 | GLYATL1 |
| ENSG00000166863 | TAC3 |
| ENSG00000166888 | STAT6 |
| ENSG00000166900 | STX3 |
| ENSG00000166922 | SCG5 |
| ENSG00000167074 | TEF |
| ENSG00000167077 | MEI1 |
| ENSG00000167100 | SAMD14 |
| ENSG00000167123 | CERCAM |
| ENSG00000167165 | UGT1A6 |
| ENSG00000167173 | C15orf39 |
| ENSG00000167193 | CRK |
| ENSG00000167207 | NOD2 |
| ENSG00000167244 | IGF2 |
| ENSG00000167264 | DUS2 |
| ENSG00000167291 | TBC1D16 |
| ENSG00000167434 | CA4 |
| ENSG00000167460 | TPM4 |
| ENSG00000167483 | FAM129C |
| ENSG00000167549 | CORO6 |
| ENSG00000167553 | TUBA1C |
| ENSG00000167562 | ZNF701 |
| ENSG00000167601 | AXL |
| ENSG00000167608 | TMC4 |
| ENSG00000167614 | TTYH1 |
| ENSG00000167632 | TRAPPC9 |
| ENSG00000167641 | PPP1R14A |
| ENSG00000167642 | SPINT2 |
| ENSG00000167685 | ZNF444 |

| | |
|---|---|
| ENSG00000167723 | TRPV3 |
| ENSG00000167733 | HSD11B1L |
| ENSG00000167766 | ZNF83 |
| ENSG00000167767 | KRT80 |
| ENSG00000167800 | TBX10 |
| ENSG00000167815 | PRDX2 |
| ENSG00000167840 | ZNF232 |
| ENSG00000167861 | HID1 |
| ENSG00000167880 | EVPL |
| ENSG00000167895 | TMC8 |
| ENSG00000167964 | RAB26 |
| ENSG00000167986 | DDB1 |
| ENSG00000168000 | BSCL2 |
| ENSG00000168036 | CTNNB1 |
| ENSG00000168071 | CCDC88B |
| ENSG00000168079 | SCARA5 |
| ENSG00000168214 | RBPJ |
| ENSG00000168234 | TTC39C |
| ENSG00000168256 | NKIRAS2 |
| ENSG00000168259 | DNAJC7 |
| ENSG00000168280 | KIF5C |
| ENSG00000168314 | MOBP |
| ENSG00000168350 | DEGS2 |
| ENSG00000168386 | FILIP1L |
| ENSG00000168398 | BDKRB2 |
| ENSG00000168421 | RHOH |
| ENSG00000168461 | RAB31 |
| ENSG00000168477 | TNXB |
| ENSG00000168490 | PHYHIP |
| ENSG00000168522 | FNTA |
| ENSG00000168528 | SERINC2 |
| ENSG00000168575 | SLC20A2 |
| ENSG00000168631 | DPCR1 |
| ENSG00000168646 | AXIN2 |
| ENSG00000168675 | LDLRAD4 |
| ENSG00000168874 | ATOH8 |
| ENSG00000168916 | ZNF608 |
| ENSG00000168961 | LGALS9 |
| ENSG00000168993 | CPLX1 |
| ENSG00000169045 | HNRNPH1 |
| ENSG00000169057 | MECP2 |
| ENSG00000169084 | DHRSX |
| ENSG00000169085 | C8orf46 |
| ENSG00000169118 | CSNK1G1 |
| ENSG00000169129 | AFAP1L2 |
| ENSG00000169169 | CPT1C |
| ENSG00000169184 | MN1 |
| ENSG00000169213 | RAB3B |
| ENSG00000169220 | RGS14 |
| ENSG00000169251 | NMD3 |

| | |
|---|---|
| ENSG00000169306 | IL1RAPL1 |
| ENSG00000169403 | PTAFR |
| ENSG00000169435 | RASSF6 |
| ENSG00000169499 | PLEKHA2 |
| ENSG00000169641 | LUZP1 |
| ENSG00000169758 | TMEM266 |
| ENSG00000169855 | ROBO1 |
| ENSG00000169894 | MUC3A |
| ENSG00000169896 | ITGAM |
| ENSG00000169902 | TPST1 |
| ENSG00000169903 | TM4SF4 |
| ENSG00000169918 | OTUD7A |
| ENSG00000169926 | KLF13 |
| ENSG00000169994 | MYO7B |
| ENSG00000170035 | UBE2E3 |
| ENSG00000170088 | TMEM192 |
| ENSG00000170099 | SERPINA6 |
| ENSG00000170113 | NIPA1 |
| ENSG00000170175 | CHRNB1 |
| ENSG00000170231 | FABP6 |
| ENSG00000170324 | FRMPD2 |
| ENSG00000170390 | DCLK2 |
| ENSG00000170412 | GPRC5C |
| ENSG00000170421 | KRT8 |
| ENSG00000170442 | KRT86 |
| ENSG00000170464 | DNAJC18 |
| ENSG00000170485 | NPAS2 |
| ENSG00000170498 | KISS1 |
| ENSG00000170525 | PFKFB3 |
| ENSG00000170542 | SERPINB9 |
| ENSG00000170615 | SLC26A5 |
| ENSG00000170703 | TTLL6 |
| ENSG00000170927 | PKHD1 |
| ENSG00000170965 | PLAC1 |
| ENSG00000171004 | HS6ST2 |
| ENSG00000171055 | FEZ2 |
| ENSG00000171119 | NRTN |
| ENSG00000171121 | KCNMB3 |
| ENSG00000171132 | PRKCE |
| ENSG00000171234 | UGT2B7 |
| ENSG00000171303 | KCNK3 |
| ENSG00000171316 | CHD7 |
| ENSG00000171368 | TPPP |
| ENSG00000171488 | LRRC8C |
| ENSG00000171530 | TBCA |
| ENSG00000171533 | MAP6 |
| ENSG00000171540 | OTP |
| ENSG00000171574 | ZNF584 |
| ENSG00000171608 | PIK3CD |
| ENSG00000171631 | P2RY6 |

| | |
|---|---|
| ENSG00000171680 | PLEKHG5 |
| ENSG00000171703 | TCEA2 |
| ENSG00000171735 | CAMTA1 |
| ENSG00000171759 | PAH |
| ENSG00000171811 | CFAP46 |
| ENSG00000171812 | COL8A2 |
| ENSG00000171823 | FBXL14 |
| ENSG00000171840 | NINJ2 |
| ENSG00000171873 | ADRA1D |
| ENSG00000171940 | ZNF217 |
| ENSG00000171962 | DRC3 |
| ENSG00000171988 | JMJD1C |
| ENSG00000171992 | SYNPO |
| ENSG00000172116 | CD8B |
| ENSG00000172264 | MACROD2 |
| ENSG00000172379 | ARNT2 |
| ENSG00000172403 | SYNPO2 |
| ENSG00000172409 | CLP1 |
| ENSG00000172426 | RSPH9 |
| ENSG00000172482 | AGXT |
| ENSG00000172497 | ACOT12 |
| ENSG00000172531 | PPP1CA |
| ENSG00000172578 | KLHL6 |
| ENSG00000172613 | RAD9A |
| ENSG00000172738 | TMEM217 |
| ENSG00000172757 | CFL1 |
| ENSG00000172794 | RAB37 |
| ENSG00000172819 | RARG |
| ENSG00000172869 | DMXL1 |
| ENSG00000172893 | DHCR7 |
| ENSG00000172927 | MYEOV |
| ENSG00000172935 | MRGPRF |
| ENSG00000172943 | PHF8 |
| ENSG00000172985 | SH3RF3 |
| ENSG00000173068 | BNC2 |
| ENSG00000173175 | ADCY5 |
| ENSG00000173200 | PARP15 |
| ENSG00000173221 | GLRX |
| ENSG00000173227 | SYT12 |
| ENSG00000173269 | MMRN2 |
| ENSG00000173320 | STOX2 |
| ENSG00000173535 | TNFRSF10C |
| ENSG00000173546 | CSPG4 |
| ENSG00000173567 | ADGRF3 |
| ENSG00000173581 | CCDC106 |
| ENSG00000173597 | SULT1B1 |
| ENSG00000173809 | TDRD12 |
| ENSG00000173898 | SPTBN2 |
| ENSG00000173950 | XXYLT1 |
| ENSG00000174145 | NWD2 |

| | |
|---|---|
| ENSG00000174226 | SNX31 |
| ENSG00000174348 | PODN |
| ENSG00000174456 | C12orf76 |
| ENSG00000174469 | CNTNAP2 |
| ENSG00000174498 | IGDCC3 |
| ENSG00000174502 | SLC26A9 |
| ENSG00000174514 | MFSD4A |
| ENSG00000174547 | MRPL11 |
| ENSG00000174564 | IL20RB |
| ENSG00000174600 | CMKLR1 |
| ENSG00000174640 | SLCO2A1 |
| ENSG00000174844 | DNAH12 |
| ENSG00000175003 | SLC22A1 |
| ENSG00000175048 | ZDHHC14 |
| ENSG00000175084 | DES |
| ENSG00000175110 | MRPS22 |
| ENSG00000175164 | ABO |
| ENSG00000175170 | FAM182B |
| ENSG00000175267 | VWA3A |
| ENSG00000175274 | TP53I11 |
| ENSG00000175287 | PHYHD1 |
| ENSG00000175318 | GRAMD2A |
| ENSG00000175344 | CHRNA7 |
| ENSG00000175354 | PTPN2 |
| ENSG00000175356 | SCUBE2 |
| ENSG00000175390 | EIF3F |
| ENSG00000175482 | POLD4 |
| ENSG00000175513 | TSGA10IP |
| ENSG00000175564 | UCP3 |
| ENSG00000175643 | RMI2 |
| ENSG00000175662 | TOM1L2 |
| ENSG00000175699 | CCDC197 |
| ENSG00000175701 | LINC00116 |
| ENSG00000175782 | SLC35E3 |
| ENSG00000175792 | RUVBL1 |
| ENSG00000175832 | ETV4 |
| ENSG00000175866 | BAIAP2 |
| ENSG00000175894 | TSPEAR |
| ENSG00000175899 | A2M |
| ENSG00000176040 | TMPRSS7 |
| ENSG00000176049 | JAKMIP2 |
| ENSG00000176155 | CCDC57 |
| ENSG00000176204 | LRRTM4 |
| ENSG00000176244 | ACBD7 |
| ENSG00000176261 | ZBTB8OS |
| ENSG00000176438 | SYNE3 |
| ENSG00000176463 | SLCO3A1 |
| ENSG00000176533 | GNG7 |
| ENSG00000176624 | MEX3C |
| ENSG00000176697 | BDNF |

| | |
|---|---|
| ENSG00000176715 | ACSF3 |
| ENSG00000176720 | BOK |
| ENSG00000176771 | NCKAP5 |
| ENSG00000176788 | BASP1 |
| ENSG00000176834 | VSIG10 |
| ENSG00000176884 | GRIN1 |
| ENSG00000176909 | MAMSTR |
| ENSG00000177030 | DEAF1 |
| ENSG00000177058 | SLC38A9 |
| ENSG00000177098 | SCN4B |
| ENSG00000177103 | DSCAML1 |
| ENSG00000177106 | EPS8L2 |
| ENSG00000177301 | KCNA2 |
| ENSG00000177311 | ZBTB38 |
| ENSG00000177359 | AC024940.1 |
| ENSG00000177380 | PPFIA3 |
| ENSG00000177398 | UMODL1 |
| ENSG00000177426 | TGIF1 |
| ENSG00000177453 | NIM1K |
| ENSG00000177548 | RABEP2 |
| ENSG00000177556 | ATOX1 |
| ENSG00000177565 | TBL1XR1 |
| ENSG00000177675 | CD163L1 |
| ENSG00000177694 | NAALADL2 |
| ENSG00000177728 | TMEM94 |
| ENSG00000177830 | CHID1 |
| ENSG00000177981 | ASB8 |
| ENSG00000177992 | SPATA31E1 |
| ENSG00000178075 | GRAMD1C |
| ENSG00000178078 | STAP2 |
| ENSG00000178096 | BOLA1 |
| ENSG00000178104 | PDE4DIP |
| ENSG00000178209 | PLEC |
| ENSG00000178226 | PRSS36 |
| ENSG00000178297 | TMPRSS9 |
| ENSG00000178404 | CEP295NL |
| ENSG00000178662 | CSRNP3 |
| ENSG00000178685 | PARP10 |
| ENSG00000178772 | CPN2 |
| ENSG00000178796 | RIIAD1 |
| ENSG00000178826 | TMEM139 |
| ENSG00000178878 | APOLD1 |
| ENSG00000178882 | RFLNA |
| ENSG00000178935 | ZNF552 |
| ENSG00000179088 | C12orf42 |
| ENSG00000179165 | PXT1 |
| ENSG00000179222 | MAGED1 |
| ENSG00000179242 | CDH4 |
| ENSG00000179335 | CLK3 |
| ENSG00000179348 | GATA2 |

| | |
|---|---|
| ENSG00000179397 | CATSPERE |
| ENSG00000179477 | ALOX12B |
| ENSG00000179528 | LBX2 |
| ENSG00000179583 | CIITA |
| ENSG00000179588 | ZFPM1 |
| ENSG00000179715 | PCED1B |
| ENSG00000179846 | NKPD1 |
| ENSG00000179873 | NLRP11 |
| ENSG00000179913 | B3GNT3 |
| ENSG00000179954 | SSC5D |
| ENSG00000180035 | ZNF48 |
| ENSG00000180061 | TMEM150B |
| ENSG00000180316 | PNPLA1 |
| ENSG00000180432 | CYP8B1 |
| ENSG00000180448 | ARHGAP45 |
| ENSG00000180525 | PRR26 |
| ENSG00000180694 | TMEM64 |
| ENSG00000180745 | CLRN3 |
| ENSG00000180815 | MAP3K15 |
| ENSG00000180822 | PSMG4 |
| ENSG00000180891 | CUEDC1 |
| ENSG00000180999 | C1orf105 |
| ENSG00000181031 | RPH3AL |
| ENSG00000181035 | SLC25A42 |
| ENSG00000181291 | TMEM132E |
| ENSG00000181333 | HEPHL1 |
| ENSG00000181355 | OFCC1 |
| ENSG00000181378 | CFAP65 |
| ENSG00000181381 | DDX60L |
| ENSG00000181409 | AATK |
| ENSG00000181652 | ATG9B |
| ENSG00000181894 | ZNF329 |
| ENSG00000181938 | GINS3 |
| ENSG00000182022 | CHST15 |
| ENSG00000182149 | IST1 |
| ENSG00000182175 | RGMA |
| ENSG00000182185 | RAD51B |
| ENSG00000182218 | HHIPL1 |
| ENSG00000182253 | SYNM |
| ENSG00000182310 | SPACA6 |
| ENSG00000182326 | C1S |
| ENSG00000182329 | KIAA2012 |
| ENSG00000182372 | CLN8 |
| ENSG00000182472 | CAPN12 |
| ENSG00000182473 | EXOC7 |
| ENSG00000182557 | SPNS3 |
| ENSG00000182568 | SATB1 |
| ENSG00000182580 | EPHB3 |
| ENSG00000182606 | TRAK1 |
| ENSG00000182621 | PLCB1 |

| | |
|---|---|
| ENSG00000182718 | ANXA2 |
| ENSG00000182752 | PAPPA |
| ENSG00000182771 | GRID1 |
| ENSG00000182795 | C1orf116 |
| ENSG00000182870 | GALNT9 |
| ENSG00000182871 | COL18A1 |
| ENSG00000182885 | ADGRG3 |
| ENSG00000182903 | ZNF721 |
| ENSG00000182919 | C11orf54 |
| ENSG00000182957 | SPATA13 |
| ENSG00000183010 | PYCR1 |
| ENSG00000183023 | SLC8A1 |
| ENSG00000183049 | CAMK1D |
| ENSG00000183091 | NEB |
| ENSG00000183092 | BEGAIN |
| ENSG00000183111 | ARHGEF37 |
| ENSG00000183134 | PTGDR2 |
| ENSG00000183145 | RIPPLY3 |
| ENSG00000183230 | CTNNA3 |
| ENSG00000183248 | PRR36 |
| ENSG00000183317 | EPHA10 |
| ENSG00000183337 | BCOR |
| ENSG00000183454 | GRIN2A |
| ENSG00000183463 | URAD |
| ENSG00000183486 | MX2 |
| ENSG00000183520 | UTP11 |
| ENSG00000183682 | BMP8A |
| ENSG00000183778 | B3GALT5 |
| ENSG00000183844 | FAM3B |
| ENSG00000183876 | ARSI |
| ENSG00000183914 | DNAH2 |
| ENSG00000183921 | SDR42E2 |
| ENSG00000183963 | SMTN |
| ENSG00000184009 | ACTG1 |
| ENSG00000184012 | TMPRSS2 |
| ENSG00000184313 | MROH7 |
| ENSG00000184347 | SLIT3 |
| ENSG00000184381 | PLA2G6 |
| ENSG00000184384 | MAML2 |
| ENSG00000184428 | TOP1MT |
| ENSG00000184471 | C1QTNF8 |
| ENSG00000184497 | TMEM255B |
| ENSG00000184613 | NELL2 |
| ENSG00000184702 | sept-05 |
| ENSG00000184792 | OSBP2 |
| ENSG00000184828 | ZBTB7C |
| ENSG00000184922 | FMNL1 |
| ENSG00000184985 | SORCS2 |
| ENSG00000184988 | TMEM106A |
| ENSG00000185015 | CA13 |

| | |
|---|---|
| ENSG00000185033 | SEMA4B |
| ENSG00000185038 | MROH2A |
| ENSG00000185049 | NELFA |
| ENSG00000185306 | C12orf56 |
| ENSG00000185324 | CDK10 |
| ENSG00000185332 | TMEM105 |
| ENSG00000185345 | PRKN |
| ENSG00000185420 | SMYD3 |
| ENSG00000185442 | FAM174B |
| ENSG00000185482 | STAC3 |
| ENSG00000185513 | L3MBTL1 |
| ENSG00000185523 | SPATA45 |
| ENSG00000185527 | PDE6G |
| ENSG00000185532 | PRKG1 |
| ENSG00000185585 | OLFML2A |
| ENSG00000185630 | PBX1 |
| ENSG00000185640 | KRT79 |
| ENSG00000185651 | UBE2L3 |
| ENSG00000185658 | BRWD1 |
| ENSG00000185666 | SYN3 |
| ENSG00000185681 | MORN5 |
| ENSG00000185739 | SRL |
| ENSG00000185787 | MORF4L1 |
| ENSG00000185792 | NLRP9 |
| ENSG00000185842 | DNAH14 |
| ENSG00000185917 | SETD4 |
| ENSG00000185920 | PTCH1 |
| ENSG00000185973 | TMLHE |
| ENSG00000185974 | GRK1 |
| ENSG00000185989 | RASA3 |
| ENSG00000186007 | LEMD1 |
| ENSG00000186020 | ZNF529 |
| ENSG00000186074 | CD300LF |
| ENSG00000186153 | WWOX |
| ENSG00000186174 | BCL9L |
| ENSG00000186188 | FFAR4 |
| ENSG00000186197 | EDARADD |
| ENSG00000186350 | RXRA |
| ENSG00000186417 | GLDN |
| ENSG00000186448 | ZNF197 |
| ENSG00000186469 | GNG2 |
| ENSG00000186487 | MYT1L |
| ENSG00000186510 | CLCNKA |
| ENSG00000186517 | ARHGAP30 |
| ENSG00000186567 | CEACAM19 |
| ENSG00000186635 | ARAP1 |
| ENSG00000186642 | PDE2A |
| ENSG00000186732 | MPPED1 |
| ENSG00000186765 | FSCN2 |
| ENSG00000186812 | ZNF397 |

| | |
|---|---|
| ENSG00000186814 | ZSCAN30 |
| ENSG00000186868 | MAPT |
| ENSG00000186889 | TMEM17 |
| ENSG00000186897 | C1QL4 |
| ENSG00000186918 | ZNF395 |
| ENSG00000186998 | EMID1 |
| ENSG00000187021 | PNLIPRP1 |
| ENSG00000187024 | PTRH1 |
| ENSG00000187079 | TEAD1 |
| ENSG00000187091 | PLCD1 |
| ENSG00000187105 | HEATR4 |
| ENSG00000187122 | SLIT1 |
| ENSG00000187134 | AKR1C1 |
| ENSG00000187147 | RNF220 |
| ENSG00000187164 | SHTN1 |
| ENSG00000187187 | ZNF546 |
| ENSG00000187323 | DCC |
| ENSG00000187391 | MAGI2 |
| ENSG00000187498 | COL4A1 |
| ENSG00000187556 | NANOS3 |
| ENSG00000187595 | ZNF385C |
| ENSG00000187608 | ISG15 |
| ENSG00000187609 | EXD3 |
| ENSG00000187630 | DHRS4L2 |
| ENSG00000187634 | SAMD11 |
| ENSG00000187672 | ERC2 |
| ENSG00000187688 | TRPV2 |
| ENSG00000187699 | C2orf88 |
| ENSG00000187720 | THSD4 |
| ENSG00000187726 | DNAJB13 |
| ENSG00000187730 | GABRD |
| ENSG00000187772 | LIN28B |
| ENSG00000187775 | DNAH17 |
| ENSG00000187783 | TMEM72 |
| ENSG00000187800 | PEAR1 |
| ENSG00000187908 | DMBT1 |
| ENSG00000187955 | COL14A1 |
| ENSG00000188001 | TPRG1 |
| ENSG00000188037 | CLCN1 |
| ENSG00000188089 | PLA2G4E |
| ENSG00000188095 | MESP2 |
| ENSG00000188158 | NHS |
| ENSG00000188243 | COMMD6 |
| ENSG00000188282 | RUFY4 |
| ENSG00000188372 | ZP3 |
| ENSG00000188385 | JAKMIP3 |
| ENSG00000188522 | FAM83G |
| ENSG00000188549 | C15orf52 |
| ENSG00000188603 | CLN3 |
| ENSG00000188677 | PARVB |

| | |
|---|---|
| ENSG00000188687 | SLC4A5 |
| ENSG00000188735 | TMEM120B |
| ENSG00000188779 | SKOR1 |
| ENSG00000188878 | FBF1 |
| ENSG00000188897 | AC099489.1 |
| ENSG00000188937 | NYX |
| ENSG00000188981 | MSANTD1 |
| ENSG00000189001 | SBSN |
| ENSG00000189067 | LITAF |
| ENSG00000189120 | SP6 |
| ENSG00000189143 | CLDN4 |
| ENSG00000189144 | ZNF573 |
| ENSG00000189157 | FAM47E |
| ENSG00000189159 | JPT1 |
| ENSG00000189233 | NUGGC |
| ENSG00000189337 | KAZN |
| ENSG00000189350 | TOGARAM2 |
| ENSG00000189403 | HMGB1 |
| ENSG00000196092 | PAX5 |
| ENSG00000196123 | KIAA0895L |
| ENSG00000196132 | MYT1 |
| ENSG00000196139 | AKR1C3 |
| ENSG00000196154 | S100A4 |
| ENSG00000196208 | GREB1 |
| ENSG00000196218 | RYR1 |
| ENSG00000196220 | SRGAP3 |
| ENSG00000196235 | SUPT5H |
| ENSG00000196284 | SUPT3H |
| ENSG00000196313 | POM121 |
| ENSG00000196358 | NTNG2 |
| ENSG00000196405 | EVL |
| ENSG00000196417 | ZNF765 |
| ENSG00000196421 | C20orf204 |
| ENSG00000196431 | CRYBA4 |
| ENSG00000196470 | SIAH1 |
| ENSG00000196482 | ESRRG |
| ENSG00000196531 | NACA |
| ENSG00000196562 | SULF2 |
| ENSG00000196565 | HBG2 |
| ENSG00000196569 | LAMA2 |
| ENSG00000196576 | PLXNB2 |
| ENSG00000196591 | HDAC2 |
| ENSG00000196597 | ZNF782 |
| ENSG00000196605 | ZNF846 |
| ENSG00000196628 | TCF4 |
| ENSG00000196639 | HRH1 |
| ENSG00000196660 | SLC30A10 |
| ENSG00000196684 | HSH2D |
| ENSG00000196743 | GM2A |
| ENSG00000196872 | KIAA1211L |

| | |
|---|---|
| ENSG00000196946 | ZNF705A |
| ENSG00000196950 | SLC39A10 |
| ENSG00000196975 | ANXA4 |
| ENSG00000196998 | WDR45 |
| ENSG00000197046 | SIGLEC15 |
| ENSG00000197056 | ZMYM1 |
| ENSG00000197181 | PIWIL2 |
| ENSG00000197183 | NOL4L |
| ENSG00000197223 | C1D |
| ENSG00000197283 | SYNGAP1 |
| ENSG00000197361 | FBXL22 |
| ENSG00000197405 | C5AR1 |
| ENSG00000197415 | VEPH1 |
| ENSG00000197448 | GSTK1 |
| ENSG00000197471 | SPN |
| ENSG00000197536 | C5orf56 |
| ENSG00000197558 | SSPO |
| ENSG00000197580 | BCO2 |
| ENSG00000197653 | DNAH10 |
| ENSG00000197816 | CCDC180 |
| ENSG00000197852 | FAM212B |
| ENSG00000197872 | FAM49A |
| ENSG00000197893 | NRAP |
| ENSG00000197912 | SPG7 |
| ENSG00000197943 | PLCG2 |
| ENSG00000197959 | DNM3 |
| ENSG00000197971 | MBP |
| ENSG00000197977 | ELOVL2 |
| ENSG00000198039 | ZNF273 |
| ENSG00000198055 | GRK6 |
| ENSG00000198089 | SFI1 |
| ENSG00000198125 | MB |
| ENSG00000198133 | TMEM229B |
| ENSG00000198157 | HMGN5 |
| ENSG00000198216 | CACNA1E |
| ENSG00000198246 | SLC29A3 |
| ENSG00000198336 | MYL4 |
| ENSG00000198353 | HOXC4 |
| ENSG00000198373 | WWP2 |
| ENSG00000198431 | TXNRD1 |
| ENSG00000198517 | MAFK |
| ENSG00000198551 | ZNF627 |
| ENSG00000198586 | TLK1 |
| ENSG00000198598 | MMP17 |
| ENSG00000198626 | RYR2 |
| ENSG00000198646 | NCOA6 |
| ENSG00000198663 | C6orf89 |
| ENSG00000198673 | FAM19A2 |
| ENSG00000198691 | ABCA4 |
| ENSG00000198715 | GLMP |

| | |
|---|---|
| ENSG00000198719 | DLL1 |
| ENSG00000198720 | ANKRD13B |
| ENSG00000198723 | TEX45 |
| ENSG00000198728 | LDB1 |
| ENSG00000198729 | PPP1R14C |
| ENSG00000198732 | SMOC1 |
| ENSG00000198796 | ALPK2 |
| ENSG00000198807 | PAX9 |
| ENSG00000198821 | CD247 |
| ENSG00000198851 | CD3E |
| ENSG00000198873 | GRK5 |
| ENSG00000198898 | CAPZA2 |
| ENSG00000198910 | L1CAM |
| ENSG00000198915 | RASGEF1A |
| ENSG00000198945 | L3MBTL3 |
| ENSG00000198947 | DMD |
| ENSG00000198948 | MFAP3L |
| ENSG00000198959 | TGM2 |
| ENSG00000203499 | IQANK1 |
| ENSG00000203697 | CAPN8 |
| ENSG00000203814 | HIST2H2BF |
| ENSG00000203867 | RBM20 |
| ENSG00000203985 | LDLRAD1 |
| ENSG00000204060 | FOXO6 |
| ENSG00000204131 | NHSL2 |
| ENSG00000204140 | CLPSL1 |
| ENSG00000204209 | DAXX |
| ENSG00000204248 | COL11A2 |
| ENSG00000204262 | COL5A2 |
| ENSG00000204396 | VWA7 |
| ENSG00000204397 | CARD16 |
| ENSG00000204516 | MICB |
| ENSG00000204520 | MICA |
| ENSG00000204540 | PSORS1C1 |
| ENSG00000204580 | DDR1 |
| ENSG00000204616 | TRIM31 |
| ENSG00000204628 | RACK1 |
| ENSG00000204634 | TBC1D8 |
| ENSG00000204767 | FAM196B |
| ENSG00000204815 | TTC25 |
| ENSG00000204866 | IGFL2 |
| ENSG00000204882 | GPR20 |
| ENSG00000204920 | ZNF155 |
| ENSG00000204954 | C12orf73 |
| ENSG00000204960 | BLACE |
| ENSG00000204991 | SPIRE2 |
| ENSG00000205078 | SYCE1L |
| ENSG00000205086 | C2orf91 |
| ENSG00000205133 | TRIQK |
| ENSG00000205212 | CCDC144NL |

| | |
|---|---|
| ENSG00000205309 | NT5M |
| ENSG00000205336 | ADGRG1 |
| ENSG00000205639 | MFSD2B |
| ENSG00000205726 | ITSN1 |
| ENSG00000205744 | DENND1C |
| ENSG00000205795 | CYS1 |
| ENSG00000205832 | C16orf96 |
| ENSG00000206077 | ZDHHC11B |
| ENSG00000206190 | ATP10A |
| ENSG00000206561 | COLQ |
| ENSG00000211584 | SLC48A1 |
| ENSG00000212719 | C17orf51 |
| ENSG00000212901 | KRTAP3-1 |
| ENSG00000213160 | KLHL23 |
| ENSG00000213213 | CCDC183 |
| ENSG00000213397 | HAUS7 |
| ENSG00000213445 | SIPA1 |
| ENSG00000213780 | GTF2H4 |
| ENSG00000213889 | PPM1N |
| ENSG00000213903 | LTB4R |
| ENSG00000213949 | ITGA1 |
| ENSG00000213999 | MEF2B |
| ENSG00000214026 | MRPL23 |
| ENSG00000214050 | FBXO16 |
| ENSG00000214226 | C17orf67 |
| ENSG00000214357 | NEURL1B |
| ENSG00000214456 | PLIN5 |
| ENSG00000214491 | SEC14L6 |
| ENSG00000214688 | C10orf105 |
| ENSG00000214711 | CAPN14 |
| ENSG00000214814 | FER1L6 |
| ENSG00000214944 | ARHGEF28 |
| ENSG00000214954 | LRRC69 |
| ENSG00000215018 | COL28A1 |
| ENSG00000215045 | GRID2IP |
| ENSG00000215182 | MUC5AC |
| ENSG00000215252 | GOLGA8B |
| ENSG00000215271 | HOMEZ |
| ENSG00000215277 | RNF212B |
| ENSG00000215440 | NPEPL1 |
| ENSG00000215529 | EFCAB8 |
| ENSG00000215910 | C1orf167 |
| ENSG00000215912 | TTC34 |
| ENSG00000217930 | PAM16 |
| ENSG00000219626 | FAM228B |
| ENSG00000221823 | PPP3R1 |
| ENSG00000221866 | PLXNA4 |
| ENSG00000221909 | FAM200A |
| ENSG00000221926 | TRIM16 |
| ENSG00000224383 | PRR29 |

| | |
|---|---|
| ENSG00000224470 | ATXN1L |
| ENSG00000225526 | MKRN2OS |
| ENSG00000225940 | C5orf67 |
| ENSG00000225968 | ELFN1 |
| ENSG00000226174 | TEX22 |
| ENSG00000226321 | CROCC2 |
| ENSG00000226690 | AC013470.2 |
| ENSG00000227471 | AKR1B15 |
| ENSG00000231672 | DIRC3 |
| ENSG00000232119 | MCTS1 |
| ENSG00000234616 | JRK |
| ENSG00000234828 | IQCM |
| ENSG00000234965 | SHISA8 |
| ENSG00000235109 | ZSCAN31 |
| ENSG00000235162 | C12orf75 |
| ENSG00000235568 | NFAM1 |
| ENSG00000236383 | LINC00854 |
| ENSG00000236699 | ARHGEF38 |
| ENSG00000239605 | STPG4 |
| ENSG00000240720 | LRRD1 |
| ENSG00000241489 | AC244197.3 |
| ENSG00000241644 | INMT |
| ENSG00000241935 | HOGA1 |
| ENSG00000241962 | AC079447.1 |
| ENSG00000242612 | DECR2 |
| ENSG00000242852 | ZNF709 |
| ENSG00000243156 | MICAL3 |
| ENSG00000243696 | AC006254.1 |
| ENSG00000243709 | LEFTY1 |
| ENSG00000243710 | CFAP57 |
| ENSG00000243811 | APOBEC3D |
| ENSG00000243910 | TUBA4B |
| ENSG00000243978 | RTL9 |
| ENSG00000244122 | UGT1A7 |
| ENSG00000244405 | ETV5 |
| ENSG00000244486 | SCARF2 |
| ENSG00000244607 | CCDC13 |
| ENSG00000246922 | UBAP1L |
| ENSG00000248405 | PRR5-ARHGAP8 |
| ENSG00000248767 | AC187653.1 |
| ENSG00000249209 | AP000311.1 |
| ENSG00000249715 | FER1L5 |
| ENSG00000249884 | RNF103-CHMP3 |
| ENSG00000249961 | TERB1 |
| ENSG00000250317 | SMIM20 |
| ENSG00000250506 | CDK3 |
| ENSG00000250722 | SELENOP |
| ENSG00000251287 | ALG1L2 |
| ENSG00000251322 | SHANK3 |
| ENSG00000253309 | SERPINE3 |

| | |
|---|---|
| ENSG00000254402 | LRRC24 |
| ENSG00000254827 | SLC22A18AS |
| ENSG00000255346 | NOX5 |
| ENSG00000255508 | AP002990.1 |
| ENSG00000255872 | AL138752.2 |
| ENSG00000256061 | DNAAF4 |
| ENSG00000256087 | ZNF432 |
| ENSG00000257335 | MGAM |
| ENSG00000257743 | MGAM2 |
| ENSG00000257923 | CUX1 |
| ENSG00000257987 | TEX49 |
| ENSG00000258102 | MAP1LC3B2 |
| ENSG00000258461 | AC012651.1 |
| ENSG00000258472 | AC005726.2 |
| ENSG00000258659 | TRIM34 |
| ENSG00000259316 | AC087632.1 |
| ENSG00000259417 | CTXND1 |
| ENSG00000260027 | HOXB7 |
| ENSG00000260220 | CCDC187 |
| ENSG00000260230 | FRRS1L |
| ENSG00000260300 | AC009119.2 |
| ENSG00000260314 | MRC1 |
| ENSG00000260456 | C16orf95 |
| ENSG00000261341 | AC010325.1 |
| ENSG00000261371 | PECAM1 |
| ENSG00000261582 | AL121753.1 |
| ENSG00000263429 | LINC00675 |
| ENSG00000263639 | MSMB |
| ENSG00000263961 | C1orf186 |
| ENSG00000264230 | ANXA8L1 |
| ENSG00000264324 | AC006030.1 |
| ENSG00000265190 | ANXA8 |
| ENSG00000266074 | BAHCC1 |
| ENSG00000266076 | AC004805.1 |
| ENSG00000266714 | MYO15B |
| ENSG00000267561 | AC093155.3 |
| ENSG00000269113 | TRABD2B |
| ENSG00000269404 | SPIB |
| ENSG00000269693 | AC010422.6 |
| ENSG00000270106 | TSNAX-DISC1 |
| ENSG00000270647 | TAF15 |
| ENSG00000271447 | MMP28 |
| ENSG00000271605 | MILR1 |
| ENSG00000272305 | AC096887.1 |
| ENSG00000272636 | DOC2B |
| ENSG00000272899 | AC025594.2 |
| ENSG00000273167 | AL359736.1 |
| ENSG00000273259 | AL049839.2 |
| ENSG00000273398 | AC017083.4 |
| ENSG00000273899 | NOL12 |

| | |
|---|---|
| ENSG00000274070 | CASTOR2 |
| ENSG00000274322 | AL136531.2 |
| ENSG00000275718 | CCL15 |
| ENSG00000275832 | ARHGAP23 |
| ENSG00000276043 | UHRF1 |
| ENSG00000277363 | SRCIN1 |
| ENSG00000278500 | AC009336.2 |
| ENSG00000278540 | ACACA |
| ENSG00000278570 | NR2E3 |
| ENSG00000281106 | LINC00282 |
| ENSG00000283154 | IQCJ-SCHIP1 |
| ENSG00000283297 | AC005841.2 |
| ENSG00000283782 | AC116366.3 |
| ENSG00000283900 | Z98749.3 |
| ENSG00000284461 | RABGEF1 |
| ENSG00000284686 | AC119674.2 |
| ENSG00000284691 | AC073111.5 |

| gene_id | gene_name | **Z2Sc condition** |
|---|---|---|
| ENSG00000001084 | GCLC | |
| ENSG00000001626 | CFTR | |
| ENSG00000001630 | CYP51A1 | |
| ENSG00000002016 | RAD52 | |
| ENSG00000002726 | AOC1 | |
| ENSG00000002746 | HECW1 | |
| ENSG00000002919 | SNX11 | |
| ENSG00000003436 | TFPI | |
| ENSG00000004139 | SARM1 | |
| ENSG00000004399 | PLXND1 | |
| ENSG00000004846 | ABCB5 | |
| ENSG00000005020 | SKAP2 | |
| ENSG00000005102 | MEOX1 | |
| ENSG00000005187 | ACSM3 | |
| ENSG00000005243 | COPZ2 | |
| ENSG00000005379 | TSPOAP1 | |
| ENSG00000005471 | ABCB4 | |
| ENSG00000005486 | RHBDD2 | |
| ENSG00000005844 | ITGAL | |
| ENSG00000005882 | PDK2 | |
| ENSG00000005961 | ITGA2B | |
| ENSG00000006016 | CRLF1 | |
| ENSG00000006071 | ABCC8 | |
| ENSG00000006282 | SPATA20 | |
| ENSG00000006283 | CACNA1G | |
| ENSG00000006652 | IFRD1 | |
| ENSG00000006715 | VPS41 | |
| ENSG00000007047 | MARK4 | |
| ENSG00000007171 | NOS2 | |
| ENSG00000007216 | SLC13A2 | |
| ENSG00000007237 | GAS7 | |
| ENSG00000007402 | CACNA2D2 | |
| ENSG00000007516 | BAIAP3 | |
| ENSG00000008324 | SS18L2 | |
| ENSG00000008441 | NFIX | |
| ENSG00000008516 | MMP25 | |
| ENSG00000008710 | PKD1 | |
| ENSG00000008838 | MED24 | |
| ENSG00000008853 | RHOBTB2 | |
| ENSG00000009790 | TRAF3IP3 | |
| ENSG00000010295 | IFFO1 | |
| ENSG00000010310 | GIPR | |
| ENSG00000010327 | STAB1 | |
| ENSG00000010379 | SLC6A13 | |
| ENSG00000010404 | IDS | |
| ENSG00000010610 | CD4 | |
| ENSG00000010626 | LRRC23 | |
| ENSG00000010810 | FYN | |
| ENSG00000010818 | HIVEP2 | |

| | |
|---|---|
| ENSG00000010932 | FMO1 |
| ENSG00000011028 | MRC2 |
| ENSG00000011083 | SLC6A7 |
| ENSG00000011105 | TSPAN9 |
| ENSG00000011332 | DPF1 |
| ENSG00000011451 | WIZ |
| ENSG00000012061 | ERCC1 |
| ENSG00000012232 | EXTL3 |
| ENSG00000013364 | MVP |
| ENSG00000013523 | ANGEL1 |
| ENSG00000013561 | RNF14 |
| ENSG00000015413 | DPEP1 |
| ENSG00000017260 | ATP2C1 |
| ENSG00000018408 | WWTR1 |
| ENSG00000018625 | ATP1A2 |
| ENSG00000019102 | VSIG2 |
| ENSG00000019144 | PHLDB1 |
| ENSG00000019169 | MARCO |
| ENSG00000019485 | PRDM11 |
| ENSG00000020181 | ADGRA2 |
| ENSG00000020256 | ZFP64 |
| ENSG00000020633 | RUNX3 |
| ENSG00000021300 | PLEKHB1 |
| ENSG00000021645 | NRXN3 |
| ENSG00000022567 | SLC45A4 |
| ENSG00000023171 | GRAMD1B |
| ENSG00000023287 | RB1CC1 |
| ENSG00000023892 | DEF6 |
| ENSG00000023902 | PLEKHO1 |
| ENSG00000024422 | EHD2 |
| ENSG00000027075 | PRKCH |
| ENSG00000028116 | VRK2 |
| ENSG00000028137 | TNFRSF1B |
| ENSG00000028277 | POU2F2 |
| ENSG00000029534 | ANK1 |
| ENSG00000031003 | FAM13B |
| ENSG00000033011 | ALG1 |
| ENSG00000034677 | RNF19A |
| ENSG00000035403 | VCL |
| ENSG00000035664 | DAPK2 |
| ENSG00000036257 | CUL3 |
| ENSG00000036448 | MYOM2 |
| ENSG00000036672 | USP2 |
| ENSG00000039139 | DNAH5 |
| ENSG00000039523 | RIPOR1 |
| ENSG00000039600 | SOX30 |
| ENSG00000040608 | RTN4R |
| ENSG00000041353 | RAB27B |
| ENSG00000041515 | MYO16 |
| ENSG00000042062 | RIPOR3 |

| | |
|---|---|
| ENSG00000042781 | USH2A |
| ENSG00000042832 | TG |
| ENSG00000043093 | DCUN1D1 |
| ENSG00000044115 | CTNNA1 |
| ENSG00000047056 | WDR37 |
| ENSG00000047346 | FAM214A |
| ENSG00000047617 | ANO2 |
| ENSG00000047662 | FAM184B |
| ENSG00000047936 | ROS1 |
| ENSG00000048342 | CC2D2A |
| ENSG00000048405 | ZNF800 |
| ENSG00000048471 | SNX29 |
| ENSG00000049283 | EPN3 |
| ENSG00000049449 | RCN1 |
| ENSG00000049618 | ARID1B |
| ENSG00000050730 | TNIP3 |
| ENSG00000052749 | RRP12 |
| ENSG00000053108 | FSTL4 |
| ENSG00000053254 | FOXN3 |
| ENSG00000053524 | MCF2L2 |
| ENSG00000053918 | KCNQ1 |
| ENSG00000054116 | TRAPPC3 |
| ENSG00000054611 | TBC1D22A |
| ENSG00000054938 | CHRDL2 |
| ENSG00000055118 | KCNH2 |
| ENSG00000055163 | CYFIP2 |
| ENSG00000055208 | TAB2 |
| ENSG00000055955 | ITIH4 |
| ENSG00000057608 | GDI2 |
| ENSG00000057704 | TMCC3 |
| ENSG00000057935 | MTA3 |
| ENSG00000058091 | CDK14 |
| ENSG00000058335 | RASGRF1 |
| ENSG00000058404 | CAMK2B |
| ENSG00000058453 | CROCC |
| ENSG00000058866 | DGKG |
| ENSG00000059728 | MXD1 |
| ENSG00000061273 | HDAC7 |
| ENSG00000061337 | LZTS1 |
| ENSG00000061938 | TNK2 |
| ENSG00000062038 | CDH3 |
| ENSG00000062282 | DGAT2 |
| ENSG00000062370 | ZNF112 |
| ENSG00000062598 | ELMO2 |
| ENSG00000063245 | EPN1 |
| ENSG00000063438 | AHRR |
| ENSG00000063978 | RNF4 |
| ENSG00000064012 | CASP8 |
| ENSG00000064300 | NGFR |
| ENSG00000064886 | CHI3L2 |

| | |
|---|---|
| ENSG00000065150 | IPO5 |
| ENSG00000065320 | NTN1 |
| ENSG00000065357 | DGKA |
| ENSG00000065534 | MYLK |
| ENSG00000065618 | COL17A1 |
| ENSG00000065717 | TLE2 |
| ENSG00000065809 | FAM107B |
| ENSG00000065882 | TBC1D1 |
| ENSG00000065989 | PDE4A |
| ENSG00000066248 | NGEF |
| ENSG00000066405 | CLDN18 |
| ENSG00000066629 | EML1 |
| ENSG00000066735 | KIF26A |
| ENSG00000067208 | EVI5 |
| ENSG00000067840 | PDZD4 |
| ENSG00000067842 | ATP2B3 |
| ENSG00000068028 | RASSF1 |
| ENSG00000068383 | INPP5A |
| ENSG00000068615 | REEP1 |
| ENSG00000068724 | TTC7A |
| ENSG00000068745 | IP6K2 |
| ENSG00000068971 | PPP2R5B |
| ENSG00000069020 | MAST4 |
| ENSG00000069424 | KCNAB2 |
| ENSG00000069667 | RORA |
| ENSG00000069702 | TGFBR3 |
| ENSG00000069956 | MAPK6 |
| ENSG00000069966 | GNB5 |
| ENSG00000069974 | RAB27A |
| ENSG00000069998 | HDHD5 |
| ENSG00000070081 | NUCB2 |
| ENSG00000070087 | PFN2 |
| ENSG00000070182 | SPTB |
| ENSG00000070269 | TMEM260 |
| ENSG00000070526 | ST6GALNAC1 |
| ENSG00000070729 | CNGB1 |
| ENSG00000070808 | CAMK2A |
| ENSG00000070915 | SLC12A3 |
| ENSG00000071073 | MGAT4A |
| ENSG00000071242 | RPS6KA2 |
| ENSG00000071909 | MYO3B |
| ENSG00000072134 | EPN2 |
| ENSG00000072135 | PTPN18 |
| ENSG00000072163 | LIMS2 |
| ENSG00000072182 | ASIC4 |
| ENSG00000072195 | SPEG |
| ENSG00000072201 | LNX1 |
| ENSG00000072422 | RHOBTB1 |
| ENSG00000072609 | CHFR |
| ENSG00000072682 | P4HA2 |

| | |
|---|---|
| ENSG00000072864 | NDE1 |
| ENSG00000072952 | MRVI1 |
| ENSG00000073060 | SCARB1 |
| ENSG00000073146 | MOV10L1 |
| ENSG00000073350 | LLGL2 |
| ENSG00000073605 | GSDMB |
| ENSG00000073670 | ADAM11 |
| ENSG00000073754 | CD5L |
| ENSG00000074047 | GLI2 |
| ENSG00000074219 | TEAD2 |
| ENSG00000074416 | MGLL |
| ENSG00000074964 | ARHGEF10L |
| ENSG00000074966 | TXK |
| ENSG00000075073 | TACR2 |
| ENSG00000075131 | TIPIN |
| ENSG00000075142 | SRI |
| ENSG00000075213 | SEMA3A |
| ENSG00000075240 | GRAMD4 |
| ENSG00000075275 | CELSR1 |
| ENSG00000075290 | WNT8B |
| ENSG00000075292 | ZNF638 |
| ENSG00000075618 | FSCN1 |
| ENSG00000075624 | ACTB |
| ENSG00000075891 | PAX2 |
| ENSG00000075945 | KIFAP3 |
| ENSG00000076356 | PLXNA2 |
| ENSG00000076554 | TPD52 |
| ENSG00000076555 | ACACB |
| ENSG00000076706 | MCAM |
| ENSG00000077063 | CTTNBP2 |
| ENSG00000077092 | RARB |
| ENSG00000077514 | POLD3 |
| ENSG00000077942 | FBLN1 |
| ENSG00000078070 | MCCC1 |
| ENSG00000078237 | TIGAR |
| ENSG00000078246 | TULP3 |
| ENSG00000078399 | HOXA9 |
| ENSG00000078687 | TNRC6C |
| ENSG00000078804 | TP53INP2 |
| ENSG00000079112 | CDH17 |
| ENSG00000079308 | TNS1 |
| ENSG00000079335 | CDC14A |
| ENSG00000079337 | RAPGEF3 |
| ENSG00000079385 | CEACAM1 |
| ENSG00000079393 | DUSP13 |
| ENSG00000079435 | LIPE |
| ENSG00000079459 | FDFT1 |
| ENSG00000080503 | SMARCA2 |
| ENSG00000080824 | HSP90AA1 |
| ENSG00000080845 | DLGAP4 |

| | |
|---|---|
| ENSG00000080854 | IGSF9B |
| ENSG00000081087 | OSTM1 |
| ENSG00000081189 | MEF2C |
| ENSG00000081248 | CACNA1S |
| ENSG00000081320 | STK17B |
| ENSG00000082014 | SMARCD3 |
| ENSG00000082074 | FYB1 |
| ENSG00000082458 | DLG3 |
| ENSG00000082684 | SEMA5B |
| ENSG00000082781 | ITGB5 |
| ENSG00000083457 | ITGAE |
| ENSG00000083807 | SLC27A5 |
| ENSG00000083828 | ZNF586 |
| ENSG00000084070 | SMAP2 |
| ENSG00000084072 | PPIE |
| ENSG00000084444 | FAM234B |
| ENSG00000084628 | NKAIN1 |
| ENSG00000084693 | AGBL5 |
| ENSG00000084710 | EFR3B |
| ENSG00000085449 | WDFY1 |
| ENSG00000085465 | OVGP1 |
| ENSG00000085563 | ABCB1 |
| ENSG00000085644 | ZNF213 |
| ENSG00000085788 | DDHD2 |
| ENSG00000085831 | TTC39A |
| ENSG00000085978 | ATG16L1 |
| ENSG00000085998 | POMGNT1 |
| ENSG00000086015 | MAST2 |
| ENSG00000086200 | IPO11 |
| ENSG00000086289 | EPDR1 |
| ENSG00000086570 | FAT2 |
| ENSG00000086730 | LAT2 |
| ENSG00000086967 | MYBPC2 |
| ENSG00000086991 | NOX4 |
| ENSG00000087008 | ACOX3 |
| ENSG00000087237 | CETP |
| ENSG00000087258 | GNAO1 |
| ENSG00000087303 | NID2 |
| ENSG00000087460 | GNAS |
| ENSG00000087495 | PHACTR3 |
| ENSG00000087903 | RFX2 |
| ENSG00000088002 | SULT2B1 |
| ENSG00000088367 | EPB41L1 |
| ENSG00000088726 | TMEM40 |
| ENSG00000088826 | SMOX |
| ENSG00000088833 | NSFL1C |
| ENSG00000088876 | ZNF343 |
| ENSG00000088881 | EBF4 |
| ENSG00000088992 | TESC |
| ENSG00000089053 | ANAPC5 |

| | |
|---|---|
| ENSG00000089057 | SLC23A2 |
| ENSG00000089060 | SLC8B1 |
| ENSG00000089101 | CFAP61 |
| ENSG00000089159 | PXN |
| ENSG00000089351 | GRAMD1A |
| ENSG00000089356 | FXYD3 |
| ENSG00000089692 | LAG3 |
| ENSG00000089693 | MLF2 |
| ENSG00000089818 | NECAP1 |
| ENSG00000089820 | ARHGAP4 |
| ENSG00000089847 | ANKRD24 |
| ENSG00000090006 | LTBP4 |
| ENSG00000090020 | SLC9A1 |
| ENSG00000090097 | PCBP4 |
| ENSG00000090266 | NDUFB2 |
| ENSG00000090512 | FETUB |
| ENSG00000090857 | PDPR |
| ENSG00000090905 | TNRC6A |
| ENSG00000090975 | PITPNM2 |
| ENSG00000091128 | LAMB4 |
| ENSG00000091428 | RAPGEF4 |
| ENSG00000091482 | SMPX |
| ENSG00000091513 | TF |
| ENSG00000091536 | MYO15A |
| ENSG00000091583 | APOH |
| ENSG00000091947 | TMEM101 |
| ENSG00000091986 | CCDC80 |
| ENSG00000092200 | RPGRIP1 |
| ENSG00000092295 | TGM1 |
| ENSG00000092529 | CAPN3 |
| ENSG00000092607 | TBX15 |
| ENSG00000092621 | PHGDH |
| ENSG00000092758 | COL9A3 |
| ENSG00000092964 | DPYSL2 |
| ENSG00000093167 | LRRFIP2 |
| ENSG00000093183 | SEC22C |
| ENSG00000095066 | HOOK2 |
| ENSG00000095303 | PTGS1 |
| ENSG00000095370 | SH2D3C |
| ENSG00000095397 | WHRN |
| ENSG00000095585 | BLNK |
| ENSG00000095587 | TLL2 |
| ENSG00000095932 | SMIM24 |
| ENSG00000096060 | FKBP5 |
| ENSG00000096088 | PGC |
| ENSG00000096093 | EFHC1 |
| ENSG00000099204 | ABLIM1 |
| ENSG00000099284 | H2AFY2 |
| ENSG00000099308 | MAST3 |
| ENSG00000099385 | BCL7C |

| | |
|---|---|
| ENSG00000099622 | CIRBP |
| ENSG00000099797 | TECR |
| ENSG00000099917 | MED15 |
| ENSG00000099940 | SNAP29 |
| ENSG00000099953 | MMP11 |
| ENSG00000099954 | CECR2 |
| ENSG00000099957 | P2RX6 |
| ENSG00000099968 | BCL2L13 |
| ENSG00000099994 | SUSD2 |
| ENSG00000099999 | RNF215 |
| ENSG00000100031 | GGT1 |
| ENSG00000100033 | PRODH |
| ENSG00000100034 | PPM1F |
| ENSG00000100092 | SH3BP1 |
| ENSG00000100106 | TRIOBP |
| ENSG00000100154 | TTC28 |
| ENSG00000100170 | SLC5A1 |
| ENSG00000100207 | TCF20 |
| ENSG00000100218 | RSPH14 |
| ENSG00000100228 | RAB36 |
| ENSG00000100242 | SUN2 |
| ENSG00000100276 | RASL10A |
| ENSG00000100280 | AP1B1 |
| ENSG00000100299 | ARSA |
| ENSG00000100307 | CBX7 |
| ENSG00000100346 | CACNA1I |
| ENSG00000100385 | IL2RB |
| ENSG00000100416 | TRMU |
| ENSG00000100433 | KCNK10 |
| ENSG00000100490 | CDKL1 |
| ENSG00000100612 | DHRS7 |
| ENSG00000100628 | ASB2 |
| ENSG00000100629 | CEP128 |
| ENSG00000100650 | SRSF5 |
| ENSG00000100902 | PSMA6 |
| ENSG00000100934 | SEC23A |
| ENSG00000101000 | PROCR |
| ENSG00000101004 | NINL |
| ENSG00000101096 | NFATC2 |
| ENSG00000101098 | RIMS4 |
| ENSG00000101144 | BMP7 |
| ENSG00000101188 | NTSR1 |
| ENSG00000101198 | NKAIN4 |
| ENSG00000101203 | COL20A1 |
| ENSG00000101204 | CHRNA4 |
| ENSG00000101224 | CDC25B |
| ENSG00000101255 | TRIB3 |
| ENSG00000101276 | SLC52A3 |
| ENSG00000101294 | HM13 |
| ENSG00000101306 | MYLK2 |

| | |
|---|---|
| ENSG00000101331 | CCM2L |
| ENSG00000101342 | TLDC2 |
| ENSG00000101464 | PIGU |
| ENSG00000101489 | CELF4 |
| ENSG00000101544 | ADNP2 |
| ENSG00000101605 | MYOM1 |
| ENSG00000101680 | LAMA1 |
| ENSG00000101773 | RBBP8 |
| ENSG00000101871 | MID1 |
| ENSG00000101901 | ALG13 |
| ENSG00000101935 | AMMECR1 |
| ENSG00000101940 | WDR13 |
| ENSG00000101974 | ATP11C |
| ENSG00000102003 | SYP |
| ENSG00000102057 | KCND1 |
| ENSG00000102144 | PGK1 |
| ENSG00000102174 | PHEX |
| ENSG00000102359 | SRPX2 |
| ENSG00000102445 | RUBCNL |
| ENSG00000102554 | KLF5 |
| ENSG00000102606 | ARHGEF7 |
| ENSG00000102755 | FLT1 |
| ENSG00000102780 | DGKH |
| ENSG00000102901 | CENPT |
| ENSG00000102921 | N4BP1 |
| ENSG00000102962 | CCL22 |
| ENSG00000102984 | ZNF821 |
| ENSG00000103021 | CCDC113 |
| ENSG00000103034 | NDRG4 |
| ENSG00000103043 | VAC14 |
| ENSG00000103056 | SMPD3 |
| ENSG00000103091 | WDR59 |
| ENSG00000103196 | CRISPLD2 |
| ENSG00000103227 | LMF1 |
| ENSG00000103264 | FBXO31 |
| ENSG00000103365 | GGA2 |
| ENSG00000103375 | AQP8 |
| ENSG00000103544 | C16orf62 |
| ENSG00000103549 | RNF40 |
| ENSG00000103647 | CORO2B |
| ENSG00000103710 | RASL12 |
| ENSG00000103740 | ACSBG1 |
| ENSG00000103769 | RAB11A |
| ENSG00000103811 | CTSH |
| ENSG00000104044 | OCA2 |
| ENSG00000104059 | FAM189A1 |
| ENSG00000104067 | TJP1 |
| ENSG00000104368 | PLAT |
| ENSG00000104381 | GDAP1 |
| ENSG00000104408 | EIF3E |

| | |
|---|---|
| ENSG00000104415 | WISP1 |
| ENSG00000104419 | NDRG1 |
| ENSG00000104447 | TRPS1 |
| ENSG00000104490 | NCALD |
| ENSG00000104626 | ERI1 |
| ENSG00000104679 | R3HCC1 |
| ENSG00000104691 | UBXN8 |
| ENSG00000104714 | ERICH1 |
| ENSG00000104728 | ARHGEF10 |
| ENSG00000104765 | BNIP3L |
| ENSG00000104856 | RELB |
| ENSG00000104866 | PPP1R37 |
| ENSG00000104870 | FCGRT |
| ENSG00000104883 | PEX11G |
| ENSG00000104889 | RNASEH2A |
| ENSG00000104892 | KLC3 |
| ENSG00000104936 | DMPK |
| ENSG00000104941 | RSPH6A |
| ENSG00000104957 | CCDC130 |
| ENSG00000105088 | OLFM2 |
| ENSG00000105176 | URI1 |
| ENSG00000105220 | GPI |
| ENSG00000105227 | PRX |
| ENSG00000105251 | SHD |
| ENSG00000105327 | BBC3 |
| ENSG00000105339 | DENND3 |
| ENSG00000105357 | MYH14 |
| ENSG00000105401 | CDC37 |
| ENSG00000105409 | ATP1A3 |
| ENSG00000105464 | GRIN2D |
| ENSG00000105514 | RAB3D |
| ENSG00000105523 | FAM83E |
| ENSG00000105538 | RASIP1 |
| ENSG00000105552 | BCAT2 |
| ENSG00000105613 | MAST1 |
| ENSG00000105642 | KCNN1 |
| ENSG00000105650 | PDE4C |
| ENSG00000105668 | UPK1A |
| ENSG00000105696 | TMEM59L |
| ENSG00000105732 | ZNF574 |
| ENSG00000105793 | GTPBP10 |
| ENSG00000105889 | STEAP1B |
| ENSG00000105929 | ATP6V0A4 |
| ENSG00000105963 | ADAP1 |
| ENSG00000105971 | CAV2 |
| ENSG00000105997 | HOXA3 |
| ENSG00000106003 | LFNG |
| ENSG00000106066 | CPVL |
| ENSG00000106069 | CHN2 |
| ENSG00000106125 | MINDY4 |

| | |
|---|---|
| ENSG00000106302 | HYAL4 |
| ENSG00000106327 | TFR2 |
| ENSG00000106355 | LSM5 |
| ENSG00000106366 | SERPINE1 |
| ENSG00000106392 | C1GALT1 |
| ENSG00000106404 | CLDN15 |
| ENSG00000106541 | AGR2 |
| ENSG00000106617 | PRKAG2 |
| ENSG00000106689 | LHX2 |
| ENSG00000106852 | LHX6 |
| ENSG00000106948 | AKNA |
| ENSG00000106976 | DNM1 |
| ENSG00000106991 | ENG |
| ENSG00000107077 | KDM4C |
| ENSG00000107104 | KANK1 |
| ENSG00000107130 | NCS1 |
| ENSG00000107187 | LHX3 |
| ENSG00000107249 | GLIS3 |
| ENSG00000107263 | RAPGEF1 |
| ENSG00000107371 | EXOSC3 |
| ENSG00000107551 | RASSF4 |
| ENSG00000107593 | PKD2L1 |
| ENSG00000107719 | PALD1 |
| ENSG00000107731 | UNC5B |
| ENSG00000107736 | CDH23 |
| ENSG00000107738 | VSIR |
| ENSG00000107742 | SPOCK2 |
| ENSG00000107798 | LIPA |
| ENSG00000107807 | TLX1 |
| ENSG00000107821 | KAZALD1 |
| ENSG00000107859 | PITX3 |
| ENSG00000107902 | LHPP |
| ENSG00000107951 | MTPAP |
| ENSG00000107954 | NEURL1 |
| ENSG00000107957 | SH3PXD2A |
| ENSG00000108100 | CCNY |
| ENSG00000108176 | DNAJC12 |
| ENSG00000108262 | GIT1 |
| ENSG00000108370 | RGS9 |
| ENSG00000108379 | WNT3 |
| ENSG00000108387 | sept-04 |
| ENSG00000108389 | MTMR4 |
| ENSG00000108465 | CDK5RAP3 |
| ENSG00000108511 | HOXB6 |
| ENSG00000108557 | RAI1 |
| ENSG00000108561 | C1QBP |
| ENSG00000108576 | SLC6A4 |
| ENSG00000108641 | B9D1 |
| ENSG00000108666 | C17orf75 |
| ENSG00000108786 | HSD17B1 |

| | |
|---|---|
| ENSG00000108823 | SGCA |
| ENSG00000108846 | ABCC3 |
| ENSG00000108852 | MPP2 |
| ENSG00000108878 | CACNG1 |
| ENSG00000109016 | DHRS7B |
| ENSG00000109101 | FOXN1 |
| ENSG00000109180 | OCIAD1 |
| ENSG00000109339 | MAPK10 |
| ENSG00000109390 | NDUFC1 |
| ENSG00000109472 | CPE |
| ENSG00000109572 | CLCN3 |
| ENSG00000109610 | SOD3 |
| ENSG00000109667 | SLC2A9 |
| ENSG00000109680 | TBC1D19 |
| ENSG00000109743 | BST1 |
| ENSG00000109846 | CRYAB |
| ENSG00000109906 | ZBTB16 |
| ENSG00000110013 | SIAE |
| ENSG00000110047 | EHD1 |
| ENSG00000110076 | NRXN2 |
| ENSG00000110375 | UPK2 |
| ENSG00000110660 | SLC35F2 |
| ENSG00000110693 | SOX6 |
| ENSG00000110696 | C11orf58 |
| ENSG00000110777 | POU2AF1 |
| ENSG00000110799 | VWF |
| ENSG00000110811 | P3H3 |
| ENSG00000110844 | PRPF40B |
| ENSG00000110871 | COQ5 |
| ENSG00000110876 | SELPLG |
| ENSG00000110934 | BIN2 |
| ENSG00000110987 | BCL7A |
| ENSG00000111144 | LTA4H |
| ENSG00000111145 | ELK3 |
| ENSG00000111181 | SLC6A12 |
| ENSG00000111186 | WNT5B |
| ENSG00000111319 | SCNN1A |
| ENSG00000111321 | LTBR |
| ENSG00000111405 | ENDOU |
| ENSG00000111424 | VDR |
| ENSG00000111452 | ADGRD1 |
| ENSG00000111664 | GNB3 |
| ENSG00000111671 | SPSB2 |
| ENSG00000111676 | ATN1 |
| ENSG00000111716 | LDHB |
| ENSG00000111846 | GCNT2 |
| ENSG00000111859 | NEDD9 |
| ENSG00000111860 | CEP85L |
| ENSG00000111863 | ADTRP |
| ENSG00000111879 | FAM184A |

| | |
|---|---|
| ENSG00000111886 | GABRR2 |
| ENSG00000111907 | TPD52L1 |
| ENSG00000111912 | NCOA7 |
| ENSG00000111913 | RIPOR2 |
| ENSG00000111961 | SASH1 |
| ENSG00000112053 | SLC26A8 |
| ENSG00000112096 | SOD2 |
| ENSG00000112137 | PHACTR1 |
| ENSG00000112146 | FBXO9 |
| ENSG00000112182 | BACH2 |
| ENSG00000112245 | PTP4A1 |
| ENSG00000112297 | CRYBG1 |
| ENSG00000112339 | HBS1L |
| ENSG00000112486 | CCR6 |
| ENSG00000112499 | SLC22A2 |
| ENSG00000112541 | PDE10A |
| ENSG00000112562 | SMOC2 |
| ENSG00000112576 | CCND3 |
| ENSG00000112584 | FAM120B |
| ENSG00000112695 | COX7A2 |
| ENSG00000112699 | GMDS |
| ENSG00000112782 | CLIC5 |
| ENSG00000112787 | FBRSL1 |
| ENSG00000112812 | PRSS16 |
| ENSG00000112981 | NME5 |
| ENSG00000113231 | PDE8B |
| ENSG00000113240 | CLK4 |
| ENSG00000113319 | RASGRF2 |
| ENSG00000113387 | SUB1 |
| ENSG00000113389 | NPR3 |
| ENSG00000113391 | FAM172A |
| ENSG00000113448 | PDE4D |
| ENSG00000113504 | SLC12A7 |
| ENSG00000113580 | NR3C1 |
| ENSG00000113594 | LIFR |
| ENSG00000113645 | WWC1 |
| ENSG00000113657 | DPYSL3 |
| ENSG00000113721 | PDGFRB |
| ENSG00000113722 | CDX1 |
| ENSG00000113749 | HRH2 |
| ENSG00000113758 | DBN1 |
| ENSG00000113763 | UNC5A |
| ENSG00000113790 | EHHADH |
| ENSG00000113889 | KNG1 |
| ENSG00000114113 | RBP2 |
| ENSG00000114115 | RBP1 |
| ENSG00000114124 | GRK7 |
| ENSG00000114268 | PFKFB4 |
| ENSG00000114349 | GNAT1 |
| ENSG00000114378 | HYAL1 |

| | |
|---|---|
| ENSG00000114416 | FXR1 |
| ENSG00000114541 | FRMD4B |
| ENSG00000114544 | SLC41A3 |
| ENSG00000114654 | EFCC1 |
| ENSG00000114735 | HEMK1 |
| ENSG00000114738 | MAPKAPK3 |
| ENSG00000114812 | VIPR1 |
| ENSG00000114982 | KANSL3 |
| ENSG00000115041 | KCNIP3 |
| ENSG00000115053 | NCL |
| ENSG00000115155 | OTOF |
| ENSG00000115165 | CYTIP |
| ENSG00000115194 | SLC30A3 |
| ENSG00000115239 | ASB3 |
| ENSG00000115266 | APC2 |
| ENSG00000115310 | RTN4 |
| ENSG00000115457 | IGFBP2 |
| ENSG00000115468 | EFHD1 |
| ENSG00000115504 | EHBP1 |
| ENSG00000115525 | ST3GAL5 |
| ENSG00000115556 | PLCD4 |
| ENSG00000115590 | IL1R2 |
| ENSG00000115594 | IL1R1 |
| ENSG00000115604 | IL18R1 |
| ENSG00000115641 | FHL2 |
| ENSG00000115648 | MLPH |
| ENSG00000115850 | LCT |
| ENSG00000115944 | COX7A2L |
| ENSG00000115963 | RND3 |
| ENSG00000115977 | AAK1 |
| ENSG00000115998 | C2orf42 |
| ENSG00000116016 | EPAS1 |
| ENSG00000116039 | ATP6V1B1 |
| ENSG00000116044 | NFE2L2 |
| ENSG00000116062 | MSH6 |
| ENSG00000116128 | BCL9 |
| ENSG00000116183 | PAPPA2 |
| ENSG00000116251 | RPL22 |
| ENSG00000116254 | CHD5 |
| ENSG00000116299 | KIAA1324 |
| ENSG00000116473 | RAP1A |
| ENSG00000116584 | ARHGEF2 |
| ENSG00000116670 | MAD2L2 |
| ENSG00000116675 | DNAJC6 |
| ENSG00000116701 | NCF2 |
| ENSG00000116833 | NR5A2 |
| ENSG00000116852 | KIF21B |
| ENSG00000116857 | TMEM9 |
| ENSG00000116977 | LGALS8 |
| ENSG00000116991 | SIPA1L2 |

| | |
|---|---|
| ENSG00000117115 | PADI2 |
| ENSG00000117245 | KIF17 |
| ENSG00000117298 | ECE1 |
| ENSG00000117305 | HMGCL |
| ENSG00000117400 | MPL |
| ENSG00000117425 | PTCH2 |
| ENSG00000117472 | TSPAN1 |
| ENSG00000117616 | RSRP1 |
| ENSG00000117632 | STMN1 |
| ENSG00000117643 | MAN1C1 |
| ENSG00000117859 | OSBPL9 |
| ENSG00000117971 | CHRNB4 |
| ENSG00000117983 | MUC5B |
| ENSG00000118004 | COLEC11 |
| ENSG00000118046 | STK11 |
| ENSG00000118094 | TREH |
| ENSG00000118160 | SLC8A2 |
| ENSG00000118242 | MREG |
| ENSG00000118257 | NRP2 |
| ENSG00000118308 | LRMP |
| ENSG00000118515 | SGK1 |
| ENSG00000118557 | PMFBP1 |
| ENSG00000118640 | VAMP8 |
| ENSG00000118690 | ARMC2 |
| ENSG00000118777 | ABCG2 |
| ENSG00000118855 | MFSD1 |
| ENSG00000118898 | PPL |
| ENSG00000118922 | KLF12 |
| ENSG00000118960 | HS1BP3 |
| ENSG00000119042 | SATB2 |
| ENSG00000119125 | GDA |
| ENSG00000119283 | TRIM67 |
| ENSG00000119396 | RAB14 |
| ENSG00000119403 | PHF19 |
| ENSG00000119574 | ZBTB45 |
| ENSG00000119630 | PGF |
| ENSG00000119650 | IFT43 |
| ENSG00000119681 | LTBP2 |
| ENSG00000119685 | TTLL5 |
| ENSG00000119698 | PPP4R4 |
| ENSG00000119771 | KLHL29 |
| ENSG00000119772 | DNMT3A |
| ENSG00000119888 | EPCAM |
| ENSG00000119913 | TECTB |
| ENSG00000119917 | IFIT3 |
| ENSG00000119922 | IFIT2 |
| ENSG00000119943 | PYROXD2 |
| ENSG00000119946 | CNNM1 |
| ENSG00000119950 | MXI1 |
| ENSG00000119969 | HELLS |

| | |
|---|---|
| ENSG00000120049 | KCNIP2 |
| ENSG00000120071 | KANSL1 |
| ENSG00000120093 | HOXB3 |
| ENSG00000120332 | TNN |
| ENSG00000120341 | SEC16B |
| ENSG00000120440 | TTLL2 |
| ENSG00000120549 | KIAA1217 |
| ENSG00000120616 | EPC1 |
| ENSG00000120645 | IQSEC3 |
| ENSG00000120647 | CCDC77 |
| ENSG00000120693 | SMAD9 |
| ENSG00000120725 | SIL1 |
| ENSG00000120784 | ZFP30 |
| ENSG00000120896 | SORBS3 |
| ENSG00000120903 | CHRNA2 |
| ENSG00000120913 | PDLIM2 |
| ENSG00000121075 | TBX4 |
| ENSG00000121101 | TEX14 |
| ENSG00000121270 | ABCC11 |
| ENSG00000121380 | BCL2L14 |
| ENSG00000121410 | A1BG |
| ENSG00000121454 | LHX4 |
| ENSG00000121577 | POPDC2 |
| ENSG00000121653 | MAPK8IP1 |
| ENSG00000121753 | ADGRB2 |
| ENSG00000121898 | CPXM2 |
| ENSG00000121933 | TMIGD3 |
| ENSG00000122012 | SV2C |
| ENSG00000122025 | FLT3 |
| ENSG00000122188 | LAX1 |
| ENSG00000122367 | LDB3 |
| ENSG00000122390 | NAA60 |
| ENSG00000122547 | EEPD1 |
| ENSG00000122711 | SPINK4 |
| ENSG00000122786 | CALD1 |
| ENSG00000122787 | AKR1D1 |
| ENSG00000122877 | EGR2 |
| ENSG00000123080 | CDKN2C |
| ENSG00000123096 | SSPN |
| ENSG00000123106 | CCDC91 |
| ENSG00000123130 | ACOT9 |
| ENSG00000123358 | NR4A1 |
| ENSG00000123384 | LRP1 |
| ENSG00000123415 | SMUG1 |
| ENSG00000123453 | SARDH |
| ENSG00000123472 | ATPAF1 |
| ENSG00000123643 | SLC36A1 |
| ENSG00000123992 | DNPEP |
| ENSG00000124003 | MOGAT1 |
| ENSG00000124006 | OBSL1 |

| | |
|---|---|
| ENSG00000124126 | PREX1 |
| ENSG00000124134 | KCNS1 |
| ENSG00000124159 | MATN4 |
| ENSG00000124203 | ZNF831 |
| ENSG00000124205 | EDN3 |
| ENSG00000124212 | PTGIS |
| ENSG00000124215 | CDH26 |
| ENSG00000124225 | PMEPA1 |
| ENSG00000124313 | IQSEC2 |
| ENSG00000124343 | XG |
| ENSG00000124357 | NAGK |
| ENSG00000124440 | HIF3A |
| ENSG00000124496 | TRERF1 |
| ENSG00000124507 | PACSIN1 |
| ENSG00000124570 | SERPINB6 |
| ENSG00000124588 | NQO2 |
| ENSG00000124596 | OARD1 |
| ENSG00000124613 | ZNF391 |
| ENSG00000124743 | KLHL31 |
| ENSG00000124749 | COL21A1 |
| ENSG00000124772 | CPNE5 |
| ENSG00000124783 | SSR1 |
| ENSG00000124813 | RUNX2 |
| ENSG00000124831 | LRRFIP1 |
| ENSG00000124839 | RAB17 |
| ENSG00000124942 | AHNAK |
| ENSG00000125037 | EMC3 |
| ENSG00000125046 | SSUH2 |
| ENSG00000125089 | SH3TC1 |
| ENSG00000125246 | CLYBL |
| ENSG00000125337 | KIF25 |
| ENSG00000125462 | C1orf61 |
| ENSG00000125618 | PAX8 |
| ENSG00000125637 | PSD4 |
| ENSG00000125648 | SLC25A23 |
| ENSG00000125726 | CD70 |
| ENSG00000125730 | C3 |
| ENSG00000125735 | TNFSF14 |
| ENSG00000125741 | OPA3 |
| ENSG00000125744 | RTN2 |
| ENSG00000125746 | EML2 |
| ENSG00000125775 | SDCBP2 |
| ENSG00000125780 | TGM3 |
| ENSG00000125818 | PSMF1 |
| ENSG00000125821 | DTD1 |
| ENSG00000125864 | BFSP1 |
| ENSG00000125871 | MGME1 |
| ENSG00000125888 | BANF2 |
| ENSG00000125966 | MMP24 |
| ENSG00000126005 | MMP24-AS1 |

| | |
|---|---|
| ENSG00000126214 | KLC1 |
| ENSG00000126217 | MCF2L |
| ENSG00000126262 | FFAR2 |
| ENSG00000126351 | THRA |
| ENSG00000126583 | PRKCG |
| ENSG00000126705 | AHDC1 |
| ENSG00000126759 | CFP |
| ENSG00000126777 | KTN1 |
| ENSG00000126803 | HSPA2 |
| ENSG00000126856 | PRDM7 |
| ENSG00000126970 | ZC4H2 |
| ENSG00000127124 | HIVEP3 |
| ENSG00000127191 | TRAF2 |
| ENSG00000127220 | ABHD8 |
| ENSG00000127325 | BEST3 |
| ENSG00000127329 | PTPRB |
| ENSG00000127377 | CRYGN |
| ENSG00000127415 | IDUA |
| ENSG00000127452 | FBXL12 |
| ENSG00000127585 | FBXL16 |
| ENSG00000127603 | MACF1 |
| ENSG00000127838 | PNKD |
| ENSG00000127863 | TNFRSF19 |
| ENSG00000127946 | HIP1 |
| ENSG00000127948 | POR |
| ENSG00000128011 | LRFN1 |
| ENSG00000128266 | GNAZ |
| ENSG00000128268 | MGAT3 |
| ENSG00000128284 | APOL3 |
| ENSG00000128294 | TPST2 |
| ENSG00000128487 | SPECC1 |
| ENSG00000128563 | PRKRIP1 |
| ENSG00000128683 | GAD1 |
| ENSG00000128805 | ARHGAP22 |
| ENSG00000128849 | CGNL1 |
| ENSG00000128918 | ALDH1A2 |
| ENSG00000128928 | IVD |
| ENSG00000128973 | CLN6 |
| ENSG00000129116 | PALLD |
| ENSG00000129159 | KCNC1 |
| ENSG00000129214 | SHBG |
| ENSG00000129244 | ATP1B2 |
| ENSG00000129353 | SLC44A2 |
| ENSG00000129521 | EGLN3 |
| ENSG00000129595 | EPB41L4A |
| ENSG00000129646 | QRICH2 |
| ENSG00000129657 | SEC14L1 |
| ENSG00000129673 | AANAT |
| ENSG00000129680 | MAP7D3 |
| ENSG00000129682 | FGF13 |

| | |
|---|---|
| ENSG00000130035 | GALNT8 |
| ENSG00000130038 | CRACR2A |
| ENSG00000130052 | STARD8 |
| ENSG00000130055 | GDPD2 |
| ENSG00000130255 | RPL36 |
| ENSG00000130287 | NCAN |
| ENSG00000130307 | USHBP1 |
| ENSG00000130312 | MRPL34 |
| ENSG00000130340 | SNX9 |
| ENSG00000130475 | FCHO1 |
| ENSG00000130477 | UNC13A |
| ENSG00000130513 | GDF15 |
| ENSG00000130528 | HRC |
| ENSG00000130561 | SAG |
| ENSG00000130584 | ZBTB46 |
| ENSG00000130643 | CALY |
| ENSG00000130700 | GATA5 |
| ENSG00000130701 | RBBP8NL |
| ENSG00000130703 | OSBPL2 |
| ENSG00000130707 | ASS1 |
| ENSG00000130711 | PRDM12 |
| ENSG00000130751 | NPAS1 |
| ENSG00000130812 | ANGPTL6 |
| ENSG00000130816 | DNMT1 |
| ENSG00000130830 | MPP1 |
| ENSG00000130844 | ZNF331 |
| ENSG00000130940 | CASZ1 |
| ENSG00000131016 | AKAP12 |
| ENSG00000131018 | SYNE1 |
| ENSG00000131037 | EPS8L1 |
| ENSG00000131044 | TTLL9 |
| ENSG00000131050 | BPIFA2 |
| ENSG00000131089 | ARHGEF9 |
| ENSG00000131115 | ZNF227 |
| ENSG00000131126 | TEX101 |
| ENSG00000131149 | GSE1 |
| ENSG00000131183 | SLC34A1 |
| ENSG00000131188 | PRR7 |
| ENSG00000131196 | NFATC1 |
| ENSG00000131242 | RAB11FIP4 |
| ENSG00000131370 | SH3BP5 |
| ENSG00000131374 | TBC1D5 |
| ENSG00000131379 | C3orf20 |
| ENSG00000131398 | KCNC3 |
| ENSG00000131408 | NR1H2 |
| ENSG00000131409 | LRRC4B |
| ENSG00000131435 | PDLIM4 |
| ENSG00000131446 | MGAT1 |
| ENSG00000131459 | GFPT2 |
| ENSG00000131473 | ACLY |

| | |
|---|---|
| ENSG00000131477 | RAMP2 |
| ENSG00000131508 | UBE2D2 |
| ENSG00000131584 | ACAP3 |
| ENSG00000131620 | ANO1 |
| ENSG00000131759 | RARA |
| ENSG00000131781 | FMO5 |
| ENSG00000131848 | ZSCAN5A |
| ENSG00000131914 | LIN28A |
| ENSG00000131981 | LGALS3 |
| ENSG00000132002 | DNAJB1 |
| ENSG00000132003 | ZSWIM4 |
| ENSG00000132005 | RFX1 |
| ENSG00000132026 | RTBDN |
| ENSG00000132205 | EMILIN2 |
| ENSG00000132256 | TRIM5 |
| ENSG00000132359 | RAP1GAP2 |
| ENSG00000132405 | TBC1D14 |
| ENSG00000132436 | FIGNL1 |
| ENSG00000132510 | KDM6B |
| ENSG00000132518 | GUCY2D |
| ENSG00000132563 | REEP2 |
| ENSG00000132604 | TERF2 |
| ENSG00000132623 | ANKEF1 |
| ENSG00000132669 | RIN2 |
| ENSG00000132670 | PTPRA |
| ENSG00000132702 | HAPLN2 |
| ENSG00000132821 | VSTM2L |
| ENSG00000132874 | SLC14A2 |
| ENSG00000132915 | PDE6A |
| ENSG00000132932 | ATP8A2 |
| ENSG00000132938 | MTUS2 |
| ENSG00000132965 | ALOX5AP |
| ENSG00000133027 | PEMT |
| ENSG00000133048 | CHI3L1 |
| ENSG00000133063 | CHIT1 |
| ENSG00000133121 | STARD13 |
| ENSG00000133131 | MORC4 |
| ENSG00000133135 | RNF128 |
| ENSG00000133142 | TCEAL4 |
| ENSG00000133195 | SLC39A11 |
| ENSG00000133226 | SRRM1 |
| ENSG00000133243 | BTBD2 |
| ENSG00000133256 | PDE6B |
| ENSG00000133328 | HRASLS2 |
| ENSG00000133424 | LARGE1 |
| ENSG00000133454 | MYO18B |
| ENSG00000133597 | ADCK2 |
| ENSG00000133657 | ATP13A3 |
| ENSG00000133687 | TMTC1 |
| ENSG00000133808 | MICALCL |

| | |
|---|---|
| ENSG00000133812 | SBF2 |
| ENSG00000133816 | MICAL2 |
| ENSG00000133818 | RRAS2 |
| ENSG00000133943 | C14orf159 |
| ENSG00000133961 | NUMB |
| ENSG00000133962 | CATSPERB |
| ENSG00000133980 | VRTN |
| ENSG00000134042 | MRO |
| ENSG00000134193 | REG4 |
| ENSG00000134201 | GSTM5 |
| ENSG00000134207 | SYT6 |
| ENSG00000134245 | WNT2B |
| ENSG00000134247 | PTGFRN |
| ENSG00000134318 | ROCK2 |
| ENSG00000134324 | LPIN1 |
| ENSG00000134369 | NAV1 |
| ENSG00000134504 | KCTD1 |
| ENSG00000134516 | DOCK2 |
| ENSG00000134686 | PHC2 |
| ENSG00000134769 | DTNA |
| ENSG00000134871 | COL4A2 |
| ENSG00000134954 | ETS1 |
| ENSG00000134955 | SLC37A2 |
| ENSG00000134986 | NREP |
| ENSG00000135048 | TMEM2 |
| ENSG00000135077 | HAVCR2 |
| ENSG00000135083 | CCNJL |
| ENSG00000135093 | USP30 |
| ENSG00000135114 | OASL |
| ENSG00000135119 | RNFT2 |
| ENSG00000135127 | BICDL1 |
| ENSG00000135205 | CCDC146 |
| ENSG00000135218 | CD36 |
| ENSG00000135250 | SRPK2 |
| ENSG00000135253 | KCP |
| ENSG00000135362 | PRR5L |
| ENSG00000135373 | EHF |
| ENSG00000135406 | PRPH |
| ENSG00000135439 | AGAP2 |
| ENSG00000135443 | KRT85 |
| ENSG00000135446 | CDK4 |
| ENSG00000135452 | TSPAN31 |
| ENSG00000135472 | FAIM2 |
| ENSG00000135549 | PKIB |
| ENSG00000135596 | MICAL1 |
| ENSG00000135631 | RAB11FIP5 |
| ENSG00000135636 | DYSF |
| ENSG00000135678 | CPM |
| ENSG00000135723 | FHOD1 |
| ENSG00000135740 | SLC9A5 |

| | |
|---|---|
| ENSG00000135766 | EGLN1 |
| ENSG00000135773 | CAPN9 |
| ENSG00000135835 | KIAA1614 |
| ENSG00000135842 | FAM129A |
| ENSG00000135845 | PIGC |
| ENSG00000135899 | SP110 |
| ENSG00000135919 | SERPINE2 |
| ENSG00000135925 | WNT10A |
| ENSG00000135931 | ARMC9 |
| ENSG00000135951 | TSGA10 |
| ENSG00000136014 | USP44 |
| ENSG00000136059 | VILL |
| ENSG00000136153 | LMO7 |
| ENSG00000136235 | GPNMB |
| ENSG00000136240 | KDELR2 |
| ENSG00000136250 | AOAH |
| ENSG00000136267 | DGKB |
| ENSG00000136273 | HUS1 |
| ENSG00000136286 | MYO1G |
| ENSG00000136379 | ABHD17C |
| ENSG00000136425 | CIB2 |
| ENSG00000136449 | MYCBPAP |
| ENSG00000136560 | TANK |
| ENSG00000136574 | GATA4 |
| ENSG00000136689 | IL1RN |
| ENSG00000136830 | FAM129B |
| ENSG00000136848 | DAB2IP |
| ENSG00000136854 | STXBP1 |
| ENSG00000136877 | FPGS |
| ENSG00000136895 | GARNL3 |
| ENSG00000136928 | GABBR2 |
| ENSG00000136935 | GOLGA1 |
| ENSG00000136944 | LMX1B |
| ENSG00000137038 | DMAC1 |
| ENSG00000137074 | APTX |
| ENSG00000137090 | DMRT1 |
| ENSG00000137135 | ARHGEF39 |
| ENSG00000137198 | GMPR |
| ENSG00000137203 | TFAP2A |
| ENSG00000137270 | GCM1 |
| ENSG00000137275 | RIPK1 |
| ENSG00000137411 | VARS2 |
| ENSG00000137463 | MGARP |
| ENSG00000137491 | SLCO2B1 |
| ENSG00000137504 | CREBZF |
| ENSG00000137507 | LRRC32 |
| ENSG00000137509 | PRCP |
| ENSG00000137672 | TRPC6 |
| ENSG00000137674 | MMP20 |
| ENSG00000137699 | TRIM29 |

| | |
|---|---|
| ENSG00000137709 | POU2F3 |
| ENSG00000137726 | FXYD6 |
| ENSG00000137767 | SQOR |
| ENSG00000137809 | ITGA11 |
| ENSG00000137815 | RTF1 |
| ENSG00000137819 | PAQR5 |
| ENSG00000137834 | SMAD6 |
| ENSG00000137841 | PLCB2 |
| ENSG00000137842 | TMEM62 |
| ENSG00000137843 | PAK6 |
| ENSG00000137857 | DUOX1 |
| ENSG00000137869 | CYP19A1 |
| ENSG00000137871 | ZNF280D |
| ENSG00000137877 | SPTBN5 |
| ENSG00000137960 | GIPC2 |
| ENSG00000137962 | ARHGAP29 |
| ENSG00000137968 | SLC44A5 |
| ENSG00000138018 | SELENOI |
| ENSG00000138030 | KHK |
| ENSG00000138061 | CYP1B1 |
| ENSG00000138079 | SLC3A1 |
| ENSG00000138080 | EMILIN1 |
| ENSG00000138100 | TRIM54 |
| ENSG00000138101 | DTNB |
| ENSG00000138119 | MYOF |
| ENSG00000138138 | ATAD1 |
| ENSG00000138152 | BTBD16 |
| ENSG00000138162 | TACC2 |
| ENSG00000138190 | EXOC6 |
| ENSG00000138315 | OIT3 |
| ENSG00000138316 | ADAMTS14 |
| ENSG00000138356 | AOX1 |
| ENSG00000138385 | SSB |
| ENSG00000138411 | HECW2 |
| ENSG00000138413 | IDH1 |
| ENSG00000138435 | CHRNA1 |
| ENSG00000138617 | PARP16 |
| ENSG00000138621 | PPCDC |
| ENSG00000138622 | HCN4 |
| ENSG00000138641 | HERC3 |
| ENSG00000138669 | PRKG2 |
| ENSG00000138696 | BMPR1B |
| ENSG00000138757 | G3BP2 |
| ENSG00000138760 | SCARB2 |
| ENSG00000138771 | SHROOM3 |
| ENSG00000138792 | ENPEP |
| ENSG00000138821 | SLC39A8 |
| ENSG00000138835 | RGS3 |
| ENSG00000139083 | ETV6 |
| ENSG00000139154 | AEBP2 |

| | |
|---|---|
| ENSG00000139160 | ETFBKMT |
| ENSG00000139174 | PRICKLE1 |
| ENSG00000139209 | SLC38A4 |
| ENSG00000139269 | INHBE |
| ENSG00000139364 | TMEM132B |
| ENSG00000139438 | FAM222A |
| ENSG00000139549 | DHH |
| ENSG00000139567 | ACVRL1 |
| ENSG00000139644 | TMBIM6 |
| ENSG00000139725 | RHOF |
| ENSG00000139865 | TTC6 |
| ENSG00000139890 | REM2 |
| ENSG00000139926 | FRMD6 |
| ENSG00000139946 | PELI2 |
| ENSG00000139988 | RDH12 |
| ENSG00000140057 | AK7 |
| ENSG00000140107 | SLC25A47 |
| ENSG00000140254 | DUOXA1 |
| ENSG00000140264 | SERF2 |
| ENSG00000140274 | DUOXA2 |
| ENSG00000140280 | LYSMD2 |
| ENSG00000140391 | TSPAN3 |
| ENSG00000140443 | IGF1R |
| ENSG00000140470 | ADAMTS17 |
| ENSG00000140479 | PCSK6 |
| ENSG00000140481 | CCDC33 |
| ENSG00000140564 | FURIN |
| ENSG00000140675 | SLC5A2 |
| ENSG00000140678 | ITGAX |
| ENSG00000140682 | TGFB1I1 |
| ENSG00000140743 | CDR2 |
| ENSG00000140795 | MYLK3 |
| ENSG00000140836 | ZFHX3 |
| ENSG00000140859 | KIFC3 |
| ENSG00000140939 | NOL3 |
| ENSG00000140941 | MAP1LC3B |
| ENSG00000140950 | TLDC1 |
| ENSG00000140961 | OSGIN1 |
| ENSG00000141052 | MYOCD |
| ENSG00000141068 | KSR1 |
| ENSG00000141127 | PRPSAP2 |
| ENSG00000141252 | VPS53 |
| ENSG00000141279 | NPEPPS |
| ENSG00000141293 | SKAP1 |
| ENSG00000141314 | RHBDL3 |
| ENSG00000141316 | SPACA3 |
| ENSG00000141337 | ARSG |
| ENSG00000141391 | PRELID3A |
| ENSG00000141404 | GNAL |
| ENSG00000141485 | SLC13A5 |

| | |
|---|---|
| ENSG00000141524 | TMC6 |
| ENSG00000141526 | SLC16A3 |
| ENSG00000141527 | CARD14 |
| ENSG00000141556 | TBCD |
| ENSG00000141564 | RPTOR |
| ENSG00000141574 | SECTM1 |
| ENSG00000141639 | MAPK4 |
| ENSG00000141736 | ERBB2 |
| ENSG00000141837 | CACNA1A |
| ENSG00000141905 | NFIC |
| ENSG00000141971 | MVB12A |
| ENSG00000142046 | TMEM91 |
| ENSG00000142089 | IFITM3 |
| ENSG00000142185 | TRPM2 |
| ENSG00000142227 | EMP3 |
| ENSG00000142303 | ADAMTS10 |
| ENSG00000142319 | SLC6A3 |
| ENSG00000142347 | MYO1F |
| ENSG00000142552 | RCN3 |
| ENSG00000142583 | SLC2A5 |
| ENSG00000142599 | RERE |
| ENSG00000142609 | CFAP74 |
| ENSG00000142611 | PRDM16 |
| ENSG00000142619 | PADI3 |
| ENSG00000142621 | FHAD1 |
| ENSG00000142623 | PADI1 |
| ENSG00000142632 | ARHGEF19 |
| ENSG00000142661 | MYOM3 |
| ENSG00000142765 | SYTL1 |
| ENSG00000142949 | PTPRF |
| ENSG00000142973 | CYP4B1 |
| ENSG00000143126 | CELSR2 |
| ENSG00000143127 | ITGA10 |
| ENSG00000143167 | GPA33 |
| ENSG00000143199 | ADCY10 |
| ENSG00000143217 | NECTIN4 |
| ENSG00000143321 | HDGF |
| ENSG00000143382 | ADAMTSL4 |
| ENSG00000143434 | SEMA6C |
| ENSG00000143479 | DYRK3 |
| ENSG00000143515 | ATP8B2 |
| ENSG00000143549 | TPM3 |
| ENSG00000143552 | NUP210L |
| ENSG00000143590 | EFNA3 |
| ENSG00000143603 | KCNN3 |
| ENSG00000143641 | GALNT2 |
| ENSG00000143669 | LYST |
| ENSG00000143753 | DEGS1 |
| ENSG00000143772 | ITPKB |
| ENSG00000143786 | CNIH3 |

| | |
|---|---|
| ENSG00000143850 | PLEKHA6 |
| ENSG00000143858 | SYT2 |
| ENSG00000143870 | PDIA6 |
| ENSG00000143882 | ATP6V1C2 |
| ENSG00000143951 | WDPCP |
| ENSG00000144029 | MRPS5 |
| ENSG00000144040 | SFXN5 |
| ENSG00000144118 | RALB |
| ENSG00000144130 | NT5DC4 |
| ENSG00000144229 | THSD7B |
| ENSG00000144407 | PTH2R |
| ENSG00000144488 | ESPNL |
| ENSG00000144560 | VGLL4 |
| ENSG00000144596 | GRIP2 |
| ENSG00000144648 | ACKR2 |
| ENSG00000144668 | ITGA9 |
| ENSG00000144711 | IQSEC1 |
| ENSG00000144712 | CAND2 |
| ENSG00000144736 | SHQ1 |
| ENSG00000144741 | SLC25A26 |
| ENSG00000144791 | LIMD1 |
| ENSG00000144824 | PHLDB2 |
| ENSG00000144847 | IGSF11 |
| ENSG00000144852 | NR1I2 |
| ENSG00000144867 | SRPRB |
| ENSG00000145016 | RUBCN |
| ENSG00000145214 | DGKQ |
| ENSG00000145287 | PLAC8 |
| ENSG00000145332 | KLHL8 |
| ENSG00000145348 | TBCK |
| ENSG00000145362 | ANK2 |
| ENSG00000145391 | SETD7 |
| ENSG00000145642 | FAM159B |
| ENSG00000145685 | LHFPL2 |
| ENSG00000145901 | TNIP1 |
| ENSG00000145949 | MYLK4 |
| ENSG00000146090 | RASGEF1C |
| ENSG00000146122 | DAAM2 |
| ENSG00000146205 | ANO7 |
| ENSG00000146216 | TTBK1 |
| ENSG00000146285 | SCML4 |
| ENSG00000146426 | TIAM2 |
| ENSG00000146433 | TMEM181 |
| ENSG00000146535 | GNA12 |
| ENSG00000146592 | CREB5 |
| ENSG00000146648 | EGFR |
| ENSG00000146700 | SSC4D |
| ENSG00000146729 | NIPSNAP2 |
| ENSG00000146733 | PSPH |
| ENSG00000146828 | SLC12A9 |

| | |
|---|---|
| ENSG00000146872 | TLK2 |
| ENSG00000146966 | DENND2A |
| ENSG00000147133 | TAF1 |
| ENSG00000147164 | SNX12 |
| ENSG00000147394 | ZNF185 |
| ENSG00000147403 | RPL10 |
| ENSG00000147485 | PXDNL |
| ENSG00000147488 | ST18 |
| ENSG00000147509 | RGS20 |
| ENSG00000147526 | TACC1 |
| ENSG00000147642 | SYBU |
| ENSG00000147676 | MAL2 |
| ENSG00000147689 | FAM83A |
| ENSG00000147872 | PLIN2 |
| ENSG00000148120 | C9orf3 |
| ENSG00000148180 | GSN |
| ENSG00000148204 | CRB2 |
| ENSG00000148219 | ASTN2 |
| ENSG00000148344 | PTGES |
| ENSG00000148357 | HMCN2 |
| ENSG00000148400 | NOTCH1 |
| ENSG00000148408 | CACNA1B |
| ENSG00000148671 | ADIRF |
| ENSG00000148700 | ADD3 |
| ENSG00000148702 | HABP2 |
| ENSG00000148795 | CYP17A1 |
| ENSG00000148814 | LRRC27 |
| ENSG00000148834 | GSTO1 |
| ENSG00000148848 | ADAM12 |
| ENSG00000148985 | PGAP2 |
| ENSG00000149016 | TUT1 |
| ENSG00000149090 | PAMR1 |
| ENSG00000149091 | DGKZ |
| ENSG00000149115 | TNKS1BP1 |
| ENSG00000149256 | TENM4 |
| ENSG00000149403 | GRIK4 |
| ENSG00000149485 | FADS1 |
| ENSG00000149527 | PLCH2 |
| ENSG00000149557 | FEZ1 |
| ENSG00000149564 | ESAM |
| ENSG00000149575 | SCN2B |
| ENSG00000149596 | JPH2 |
| ENSG00000149599 | DUSP15 |
| ENSG00000149633 | KIAA1755 |
| ENSG00000149646 | CNBD2 |
| ENSG00000149823 | VPS51 |
| ENSG00000149925 | ALDOA |
| ENSG00000149932 | TMEM219 |
| ENSG00000149972 | CNTN5 |
| ENSG00000150054 | MPP7 |

| | |
|---|---|
| ENSG00000150093 | ITGB1 |
| ENSG00000150471 | ADGRL3 |
| ENSG00000150527 | CTAGE5 |
| ENSG00000150656 | CNDP1 |
| ENSG00000150667 | FSIP1 |
| ENSG00000150687 | PRSS23 |
| ENSG00000150722 | PPP1R1C |
| ENSG00000150750 | C11orf53 |
| ENSG00000150764 | DIXDC1 |
| ENSG00000150967 | ABCB9 |
| ENSG00000151062 | CACNA2D4 |
| ENSG00000151065 | DCP1B |
| ENSG00000151067 | CACNA1C |
| ENSG00000151117 | TMEM86A |
| ENSG00000151240 | DIP2C |
| ENSG00000151320 | AKAP6 |
| ENSG00000151364 | KCTD14 |
| ENSG00000151388 | ADAMTS12 |
| ENSG00000151413 | NUBPL |
| ENSG00000151468 | CCDC3 |
| ENSG00000151474 | FRMD4A |
| ENSG00000151491 | EPS8 |
| ENSG00000151572 | ANO4 |
| ENSG00000151617 | EDNRA |
| ENSG00000151632 | AKR1C2 |
| ENSG00000151655 | ITIH2 |
| ENSG00000151726 | ACSL1 |
| ENSG00000151834 | GABRA2 |
| ENSG00000151835 | SACS |
| ENSG00000151882 | CCL28 |
| ENSG00000151914 | DST |
| ENSG00000152128 | TMEM163 |
| ENSG00000152147 | GEMIN6 |
| ENSG00000152253 | SPC25 |
| ENSG00000152284 | TCF7L1 |
| ENSG00000152503 | TRIM36 |
| ENSG00000152558 | TMEM123 |
| ENSG00000152601 | MBNL1 |
| ENSG00000152684 | PELO |
| ENSG00000152689 | RASGRP3 |
| ENSG00000152700 | SAR1B |
| ENSG00000152936 | LMNTD1 |
| ENSG00000153029 | MR1 |
| ENSG00000153046 | CDYL |
| ENSG00000153060 | TEKT5 |
| ENSG00000153064 | BANK1 |
| ENSG00000153093 | ACOXL |
| ENSG00000153113 | CAST |
| ENSG00000153130 | SCOC |
| ENSG00000153179 | RASSF3 |

| | |
|---|---|
| ENSG00000153233 | PTPRR |
| ENSG00000153234 | NR4A2 |
| ENSG00000153283 | CD96 |
| ENSG00000153294 | ADGRF4 |
| ENSG00000153303 | FRMD1 |
| ENSG00000153339 | TRAPPC8 |
| ENSG00000153531 | ADPRHL1 |
| ENSG00000153707 | PTPRD |
| ENSG00000153814 | JAZF1 |
| ENSG00000153902 | LGI4 |
| ENSG00000153914 | SREK1 |
| ENSG00000153930 | ANKFN1 |
| ENSG00000154016 | GRAP |
| ENSG00000154025 | SLC5A10 |
| ENSG00000154118 | JPH3 |
| ENSG00000154134 | ROBO3 |
| ENSG00000154146 | NRGN |
| ENSG00000154153 | RETREG1 |
| ENSG00000154227 | CERS3 |
| ENSG00000154262 | ABCA6 |
| ENSG00000154305 | MIA3 |
| ENSG00000154319 | FAM167A |
| ENSG00000154358 | OBSCN |
| ENSG00000154556 | SORBS2 |
| ENSG00000154678 | PDE1C |
| ENSG00000154710 | RABGEF1 |
| ENSG00000154783 | FGD5 |
| ENSG00000154928 | EPHB1 |
| ENSG00000155093 | PTPRN2 |
| ENSG00000155096 | AZIN1 |
| ENSG00000155265 | GOLGA7B |
| ENSG00000155324 | GRAMD2B |
| ENSG00000155465 | SLC7A7 |
| ENSG00000155506 | LARP1 |
| ENSG00000155545 | MIER3 |
| ENSG00000155629 | PIK3AP1 |
| ENSG00000155749 | ALS2CR12 |
| ENSG00000155754 | C2CD6 |
| ENSG00000155849 | ELMO1 |
| ENSG00000155893 | PXYLP1 |
| ENSG00000155926 | SLA |
| ENSG00000155959 | VBP1 |
| ENSG00000155980 | KIF5A |
| ENSG00000156011 | PSD3 |
| ENSG00000156030 | ELMSAN1 |
| ENSG00000156113 | KCNMA1 |
| ENSG00000156127 | BATF |
| ENSG00000156170 | NDUFAF6 |
| ENSG00000156206 | CFAP161 |
| ENSG00000156219 | ART3 |

| | |
|---|---|
| ENSG00000156222 | SLC28A1 |
| ENSG00000156273 | BACH1 |
| ENSG00000156411 | C14orf2 |
| ENSG00000156475 | PPP2R2B |
| ENSG00000156500 | FAM122C |
| ENSG00000156515 | HK1 |
| ENSG00000156574 | NODAL |
| ENSG00000156671 | SAMD8 |
| ENSG00000156711 | MAPK13 |
| ENSG00000156802 | ATAD2 |
| ENSG00000156853 | ZNF689 |
| ENSG00000157064 | NMNAT2 |
| ENSG00000157152 | SYN2 |
| ENSG00000157214 | STEAP2 |
| ENSG00000157216 | SSBP3 |
| ENSG00000157224 | CLDN12 |
| ENSG00000157349 | DDX19B |
| ENSG00000157350 | ST3GAL2 |
| ENSG00000157368 | IL34 |
| ENSG00000157388 | CACNA1D |
| ENSG00000157399 | ARSE |
| ENSG00000157429 | ZNF19 |
| ENSG00000157450 | RNF111 |
| ENSG00000157470 | FAM81A |
| ENSG00000157510 | AFAP1L1 |
| ENSG00000157538 | DSCR3 |
| ENSG00000157601 | MX1 |
| ENSG00000157625 | TAB3 |
| ENSG00000157657 | ZNF618 |
| ENSG00000157703 | SVOPL |
| ENSG00000157800 | SLC37A3 |
| ENSG00000157833 | GAREM2 |
| ENSG00000157851 | DPYSL5 |
| ENSG00000157856 | DRC1 |
| ENSG00000157890 | MEGF11 |
| ENSG00000158008 | EXTL1 |
| ENSG00000158019 | BABAM2 |
| ENSG00000158022 | TRIM63 |
| ENSG00000158023 | WDR66 |
| ENSG00000158062 | UBXN11 |
| ENSG00000158104 | HPD |
| ENSG00000158113 | LRRC43 |
| ENSG00000158169 | FANCC |
| ENSG00000158270 | COLEC12 |
| ENSG00000158290 | CUL4B |
| ENSG00000158321 | AUTS2 |
| ENSG00000158445 | KCNB1 |
| ENSG00000158458 | NRG2 |
| ENSG00000158525 | CPA5 |
| ENSG00000158552 | ZFAND2B |

| | |
|---|---|
| ENSG00000158555 | GDPD5 |
| ENSG00000158560 | DYNC1I1 |
| ENSG00000158683 | PKD1L1 |
| ENSG00000158691 | ZSCAN12 |
| ENSG00000158747 | NBL1 |
| ENSG00000158806 | NPM2 |
| ENSG00000158856 | DMTN |
| ENSG00000158859 | ADAMTS4 |
| ENSG00000158955 | WNT9B |
| ENSG00000158985 | CDC42SE2 |
| ENSG00000159164 | SV2A |
| ENSG00000159173 | TNNI1 |
| ENSG00000159176 | CSRP1 |
| ENSG00000159200 | RCAN1 |
| ENSG00000159216 | RUNX1 |
| ENSG00000159261 | CLDN14 |
| ENSG00000159307 | SCUBE1 |
| ENSG00000159314 | ARHGAP27 |
| ENSG00000159337 | PLA2G4D |
| ENSG00000159403 | C1R |
| ENSG00000159450 | TCHH |
| ENSG00000159618 | ADGRG5 |
| ENSG00000159625 | DRC7 |
| ENSG00000159650 | UROC1 |
| ENSG00000159674 | SPON2 |
| ENSG00000159685 | CHCHD6 |
| ENSG00000159714 | ZDHHC1 |
| ENSG00000159733 | ZFYVE28 |
| ENSG00000159842 | ABR |
| ENSG00000159871 | LYPD5 |
| ENSG00000159899 | NPR2 |
| ENSG00000159921 | GNE |
| ENSG00000160111 | CPAMD8 |
| ENSG00000160145 | KALRN |
| ENSG00000160179 | ABCG1 |
| ENSG00000160183 | TMPRSS3 |
| ENSG00000160190 | SLC37A1 |
| ENSG00000160191 | PDE9A |
| ENSG00000160216 | AGPAT3 |
| ENSG00000160255 | ITGB2 |
| ENSG00000160323 | ADAMTS13 |
| ENSG00000160408 | ST6GALNAC6 |
| ENSG00000160447 | PKN3 |
| ENSG00000160460 | SPTBN4 |
| ENSG00000160593 | JAML |
| ENSG00000160746 | ANO10 |
| ENSG00000160789 | LMNA |
| ENSG00000160799 | CCDC12 |
| ENSG00000160808 | MYL3 |
| ENSG00000160838 | LRRC71 |

| | |
|---|---|
| ENSG00000160959 | LRRC14 |
| ENSG00000161010 | MRNIP |
| ENSG00000161011 | SQSTM1 |
| ENSG00000161031 | PGLYRP2 |
| ENSG00000161091 | MFSD12 |
| ENSG00000161217 | PCYT1A |
| ENSG00000161249 | DMKN |
| ENSG00000161267 | BDH1 |
| ENSG00000161270 | NPHS1 |
| ENSG00000161395 | PGAP3 |
| ENSG00000161509 | GRIN2C |
| ENSG00000161542 | PRPSAP1 |
| ENSG00000161544 | CYGB |
| ENSG00000161558 | TMEM143 |
| ENSG00000161609 | CCDC155 |
| ENSG00000161642 | ZNF385A |
| ENSG00000161800 | RACGAP1 |
| ENSG00000161813 | LARP4 |
| ENSG00000161888 | SPC24 |
| ENSG00000161914 | ZNF653 |
| ENSG00000162066 | AMDHD2 |
| ENSG00000162078 | ZG16B |
| ENSG00000162104 | ADCY9 |
| ENSG00000162148 | PPP1R32 |
| ENSG00000162341 | TPCN2 |
| ENSG00000162383 | SLC1A7 |
| ENSG00000162398 | LEXM |
| ENSG00000162413 | KLHL21 |
| ENSG00000162415 | ZSWIM5 |
| ENSG00000162438 | CTRC |
| ENSG00000162458 | FBLIM1 |
| ENSG00000162490 | DRAXIN |
| ENSG00000162510 | MATN1 |
| ENSG00000162511 | LAPTM5 |
| ENSG00000162576 | MXRA8 |
| ENSG00000162585 | FAAP20 |
| ENSG00000162599 | NFIA |
| ENSG00000162669 | HFM1 |
| ENSG00000162733 | DDR2 |
| ENSG00000162746 | FCRLB |
| ENSG00000162761 | LMX1A |
| ENSG00000162772 | ATF3 |
| ENSG00000162849 | KIF26B |
| ENSG00000162896 | PIGR |
| ENSG00000162909 | CAPN2 |
| ENSG00000162946 | DISC1 |
| ENSG00000162949 | CAPN13 |
| ENSG00000162972 | MAIP1 |
| ENSG00000163002 | NUP35 |
| ENSG00000163017 | ACTG2 |

| | |
|---|---|
| ENSG00000163050 | COQ8A |
| ENSG00000163072 | NOSTRIN |
| ENSG00000163131 | CTSS |
| ENSG00000163138 | PACRGL |
| ENSG00000163171 | CDC42EP3 |
| ENSG00000163191 | S100A11 |
| ENSG00000163218 | PGLYRP4 |
| ENSG00000163263 | C1orf189 |
| ENSG00000163288 | GABRB1 |
| ENSG00000163297 | ANTXR2 |
| ENSG00000163322 | ABRAXAS1 |
| ENSG00000163346 | PBXIP1 |
| ENSG00000163357 | DCST1 |
| ENSG00000163395 | IGFN1 |
| ENSG00000163406 | SLC15A2 |
| ENSG00000163412 | EIF4E3 |
| ENSG00000163430 | FSTL1 |
| ENSG00000163453 | IGFBP7 |
| ENSG00000163485 | ADORA1 |
| ENSG00000163520 | FBLN2 |
| ENSG00000163531 | NFASC |
| ENSG00000163535 | SGO2 |
| ENSG00000163635 | ATXN7 |
| ENSG00000163637 | PRICKLE2 |
| ENSG00000163704 | PRRT3 |
| ENSG00000163714 | U2SURP |
| ENSG00000163738 | MTHFD2L |
| ENSG00000163785 | RYK |
| ENSG00000163803 | PLB1 |
| ENSG00000163806 | SPDYA |
| ENSG00000163818 | LZTFL1 |
| ENSG00000163827 | LRRC2 |
| ENSG00000163866 | SMIM12 |
| ENSG00000163870 | TPRA1 |
| ENSG00000163885 | CFAP100 |
| ENSG00000163898 | LIPH |
| ENSG00000163904 | SENP2 |
| ENSG00000163923 | RPL39L |
| ENSG00000163959 | SLC51A |
| ENSG00000163964 | PIGX |
| ENSG00000163975 | MELTF |
| ENSG00000163995 | ABLIM2 |
| ENSG00000164007 | CLDN19 |
| ENSG00000164038 | SLC9B2 |
| ENSG00000164054 | SHISA5 |
| ENSG00000164066 | INTU |
| ENSG00000164081 | TEX264 |
| ENSG00000164093 | PITX2 |
| ENSG00000164096 | C4orf3 |
| ENSG00000164124 | TMEM144 |

| | |
|---|---|
| ENSG00000164175 | SLC45A2 |
| ENSG00000164253 | WDR41 |
| ENSG00000164292 | RHOBTB3 |
| ENSG00000164303 | ENPP6 |
| ENSG00000164309 | CMYA5 |
| ENSG00000164363 | SLC6A18 |
| ENSG00000164398 | ACSL6 |
| ENSG00000164402 | sept-08 |
| ENSG00000164465 | DCBLD1 |
| ENSG00000164488 | DACT2 |
| ENSG00000164512 | ANKRD55 |
| ENSG00000164530 | PI16 |
| ENSG00000164535 | DAGLB |
| ENSG00000164574 | GALNT10 |
| ENSG00000164638 | SLC29A4 |
| ENSG00000164674 | SYTL3 |
| ENSG00000164690 | SHH |
| ENSG00000164713 | BRI3 |
| ENSG00000164741 | DLC1 |
| ENSG00000164742 | ADCY1 |
| ENSG00000164808 | SPIDR |
| ENSG00000164830 | OXR1 |
| ENSG00000164855 | TMEM184A |
| ENSG00000164867 | NOS3 |
| ENSG00000164941 | INTS8 |
| ENSG00000164951 | PDP1 |
| ENSG00000164972 | C9orf24 |
| ENSG00000165124 | SVEP1 |
| ENSG00000165140 | FBP1 |
| ENSG00000165238 | WNK2 |
| ENSG00000165424 | ZCCHC24 |
| ENSG00000165449 | SLC16A9 |
| ENSG00000165490 | DDIAS |
| ENSG00000165507 | C10orf10 |
| ENSG00000165548 | TMEM63C |
| ENSG00000165568 | AKR1E2 |
| ENSG00000165617 | DACT1 |
| ENSG00000165626 | BEND7 |
| ENSG00000165695 | AK8 |
| ENSG00000165702 | GFI1B |
| ENSG00000165752 | STK32C |
| ENSG00000165757 | JCAD |
| ENSG00000165775 | FUNDC2 |
| ENSG00000165810 | BTNL9 |
| ENSG00000165821 | SALL2 |
| ENSG00000165868 | HSPA12A |
| ENSG00000165914 | TTC7B |
| ENSG00000165917 | RAPSN |
| ENSG00000165929 | TC2N |
| ENSG00000165959 | CLMN |

| | |
|---|---|
| ENSG00000165973 | NELL1 |
| ENSG00000165995 | CACNB2 |
| ENSG00000166016 | ABTB2 |
| ENSG00000166025 | AMOTL1 |
| ENSG00000166033 | HTRA1 |
| ENSG00000166035 | LIPC |
| ENSG00000166111 | SVOP |
| ENSG00000166159 | LRTM2 |
| ENSG00000166220 | TBATA |
| ENSG00000166268 | MYRFL |
| ENSG00000166278 | C2 |
| ENSG00000166317 | SYNPO2L |
| ENSG00000166341 | DCHS1 |
| ENSG00000166359 | WDR88 |
| ENSG00000166391 | MOGAT2 |
| ENSG00000166394 | CYB5R2 |
| ENSG00000166402 | TUB |
| ENSG00000166411 | IDH3A |
| ENSG00000166415 | WDR72 |
| ENSG00000166444 | ST5 |
| ENSG00000166509 | CLEC3A |
| ENSG00000166532 | RIMKLB |
| ENSG00000166579 | NDEL1 |
| ENSG00000166669 | ATF7IP2 |
| ENSG00000166682 | TMPRSS5 |
| ENSG00000166743 | ACSM1 |
| ENSG00000166793 | YPEL4 |
| ENSG00000166819 | PLIN1 |
| ENSG00000166825 | ANPEP |
| ENSG00000166833 | NAV2 |
| ENSG00000166840 | GLYATL1 |
| ENSG00000166863 | TAC3 |
| ENSG00000166888 | STAT6 |
| ENSG00000166900 | STX3 |
| ENSG00000166922 | SCG5 |
| ENSG00000167074 | TEF |
| ENSG00000167077 | MEI1 |
| ENSG00000167100 | SAMD14 |
| ENSG00000167123 | CERCAM |
| ENSG00000167165 | UGT1A6 |
| ENSG00000167173 | C15orf39 |
| ENSG00000167193 | CRK |
| ENSG00000167207 | NOD2 |
| ENSG00000167244 | IGF2 |
| ENSG00000167264 | DUS2 |
| ENSG00000167291 | TBC1D16 |
| ENSG00000167434 | CA4 |
| ENSG00000167460 | TPM4 |
| ENSG00000167483 | FAM129C |
| ENSG00000167549 | CORO6 |

| | |
|---|---|
| ENSG00000167553 | TUBA1C |
| ENSG00000167562 | ZNF701 |
| ENSG00000167601 | AXL |
| ENSG00000167608 | TMC4 |
| ENSG00000167614 | TTYH1 |
| ENSG00000167632 | TRAPPC9 |
| ENSG00000167641 | PPP1R14A |
| ENSG00000167642 | SPINT2 |
| ENSG00000167685 | ZNF444 |
| ENSG00000167723 | TRPV3 |
| ENSG00000167733 | HSD11B1L |
| ENSG00000167766 | ZNF83 |
| ENSG00000167767 | KRT80 |
| ENSG00000167800 | TBX10 |
| ENSG00000167815 | PRDX2 |
| ENSG00000167840 | ZNF232 |
| ENSG00000167861 | HID1 |
| ENSG00000167880 | EVPL |
| ENSG00000167895 | TMC8 |
| ENSG00000167986 | DDB1 |
| ENSG00000168000 | BSCL2 |
| ENSG00000168036 | CTNNB1 |
| ENSG00000168071 | CCDC88B |
| ENSG00000168079 | SCARA5 |
| ENSG00000168214 | RBPJ |
| ENSG00000168234 | TTC39C |
| ENSG00000168256 | NKIRAS2 |
| ENSG00000168259 | DNAJC7 |
| ENSG00000168280 | KIF5C |
| ENSG00000168306 | ACOX2 |
| ENSG00000168314 | MOBP |
| ENSG00000168350 | DEGS2 |
| ENSG00000168386 | FILIP1L |
| ENSG00000168395 | ING5 |
| ENSG00000168398 | BDKRB2 |
| ENSG00000168421 | RHOH |
| ENSG00000168461 | RAB31 |
| ENSG00000168477 | TNXB |
| ENSG00000168490 | PHYHIP |
| ENSG00000168522 | FNTA |
| ENSG00000168528 | SERINC2 |
| ENSG00000168575 | SLC20A2 |
| ENSG00000168631 | DPCR1 |
| ENSG00000168646 | AXIN2 |
| ENSG00000168675 | LDLRAD4 |
| ENSG00000168874 | ATOH8 |
| ENSG00000168916 | ZNF608 |
| ENSG00000168961 | LGALS9 |
| ENSG00000168993 | CPLX1 |
| ENSG00000169045 | HNRNPH1 |

| | |
|---|---|
| ENSG00000169057 | MECP2 |
| ENSG00000169084 | DHRSX |
| ENSG00000169085 | C8orf46 |
| ENSG00000169118 | CSNK1G1 |
| ENSG00000169129 | AFAP1L2 |
| ENSG00000169169 | CPT1C |
| ENSG00000169184 | MN1 |
| ENSG00000169213 | RAB3B |
| ENSG00000169220 | RGS14 |
| ENSG00000169251 | NMD3 |
| ENSG00000169306 | IL1RAPL1 |
| ENSG00000169403 | PTAFR |
| ENSG00000169435 | RASSF6 |
| ENSG00000169499 | PLEKHA2 |
| ENSG00000169641 | LUZP1 |
| ENSG00000169758 | TMEM266 |
| ENSG00000169855 | ROBO1 |
| ENSG00000169894 | MUC3A |
| ENSG00000169896 | ITGAM |
| ENSG00000169902 | TPST1 |
| ENSG00000169903 | TM4SF4 |
| ENSG00000169918 | OTUD7A |
| ENSG00000169926 | KLF13 |
| ENSG00000169994 | MYO7B |
| ENSG00000170035 | UBE2E3 |
| ENSG00000170049 | KCNAB3 |
| ENSG00000170088 | TMEM192 |
| ENSG00000170099 | SERPINA6 |
| ENSG00000170113 | NIPA1 |
| ENSG00000170175 | CHRNB1 |
| ENSG00000170231 | FABP6 |
| ENSG00000170324 | FRMPD2 |
| ENSG00000170390 | DCLK2 |
| ENSG00000170412 | GPRC5C |
| ENSG00000170421 | KRT8 |
| ENSG00000170442 | KRT86 |
| ENSG00000170464 | DNAJC18 |
| ENSG00000170485 | NPAS2 |
| ENSG00000170498 | KISS1 |
| ENSG00000170525 | PFKFB3 |
| ENSG00000170542 | SERPINB9 |
| ENSG00000170615 | SLC26A5 |
| ENSG00000170634 | ACYP2 |
| ENSG00000170703 | TTLL6 |
| ENSG00000170927 | PKHD1 |
| ENSG00000170965 | PLAC1 |
| ENSG00000171004 | HS6ST2 |
| ENSG00000171055 | FEZ2 |
| ENSG00000171119 | NRTN |
| ENSG00000171121 | KCNMB3 |

| | |
|---|---|
| ENSG00000171132 | PRKCE |
| ENSG00000171234 | UGT2B7 |
| ENSG00000171303 | KCNK3 |
| ENSG00000171316 | CHD7 |
| ENSG00000171368 | TPPP |
| ENSG00000171488 | LRRC8C |
| ENSG00000171530 | TBCA |
| ENSG00000171533 | MAP6 |
| ENSG00000171540 | OTP |
| ENSG00000171574 | ZNF584 |
| ENSG00000171608 | PIK3CD |
| ENSG00000171631 | P2RY6 |
| ENSG00000171680 | PLEKHG5 |
| ENSG00000171703 | TCEA2 |
| ENSG00000171735 | CAMTA1 |
| ENSG00000171759 | PAH |
| ENSG00000171811 | CFAP46 |
| ENSG00000171812 | COL8A2 |
| ENSG00000171823 | FBXL14 |
| ENSG00000171840 | NINJ2 |
| ENSG00000171873 | ADRA1D |
| ENSG00000171940 | ZNF217 |
| ENSG00000171962 | DRC3 |
| ENSG00000171988 | JMJD1C |
| ENSG00000171992 | SYNPO |
| ENSG00000172116 | CD8B |
| ENSG00000172264 | MACROD2 |
| ENSG00000172379 | ARNT2 |
| ENSG00000172403 | SYNPO2 |
| ENSG00000172409 | CLP1 |
| ENSG00000172426 | RSPH9 |
| ENSG00000172482 | AGXT |
| ENSG00000172497 | ACOT12 |
| ENSG00000172531 | PPP1CA |
| ENSG00000172578 | KLHL6 |
| ENSG00000172613 | RAD9A |
| ENSG00000172738 | TMEM217 |
| ENSG00000172757 | CFL1 |
| ENSG00000172771 | EFCAB12 |
| ENSG00000172794 | RAB37 |
| ENSG00000172819 | RARG |
| ENSG00000172869 | DMXL1 |
| ENSG00000172893 | DHCR7 |
| ENSG00000172927 | MYEOV |
| ENSG00000172935 | MRGPRF |
| ENSG00000172943 | PHF8 |
| ENSG00000172985 | SH3RF3 |
| ENSG00000173068 | BNC2 |
| ENSG00000173175 | ADCY5 |
| ENSG00000173200 | PARP15 |

| | |
|---|---|
| ENSG00000173221 | GLRX |
| ENSG00000173227 | SYT12 |
| ENSG00000173269 | MMRN2 |
| ENSG00000173320 | STOX2 |
| ENSG00000173535 | TNFRSF10C |
| ENSG00000173546 | CSPG4 |
| ENSG00000173567 | ADGRF3 |
| ENSG00000173581 | CCDC106 |
| ENSG00000173597 | SULT1B1 |
| ENSG00000173809 | TDRD12 |
| ENSG00000173898 | SPTBN2 |
| ENSG00000173950 | XXYLT1 |
| ENSG00000174136 | RGMB |
| ENSG00000174145 | NWD2 |
| ENSG00000174226 | SNX31 |
| ENSG00000174348 | PODN |
| ENSG00000174456 | C12orf76 |
| ENSG00000174469 | CNTNAP2 |
| ENSG00000174498 | IGDCC3 |
| ENSG00000174502 | SLC26A9 |
| ENSG00000174514 | MFSD4A |
| ENSG00000174547 | MRPL11 |
| ENSG00000174564 | IL20RB |
| ENSG00000174600 | CMKLR1 |
| ENSG00000174640 | SLCO2A1 |
| ENSG00000174844 | DNAH12 |
| ENSG00000175003 | SLC22A1 |
| ENSG00000175048 | ZDHHC14 |
| ENSG00000175084 | DES |
| ENSG00000175110 | MRPS22 |
| ENSG00000175164 | ABO |
| ENSG00000175170 | FAM182B |
| ENSG00000175267 | VWA3A |
| ENSG00000175274 | TP53I11 |
| ENSG00000175287 | PHYHD1 |
| ENSG00000175318 | GRAMD2A |
| ENSG00000175344 | CHRNA7 |
| ENSG00000175354 | PTPN2 |
| ENSG00000175356 | SCUBE2 |
| ENSG00000175390 | EIF3F |
| ENSG00000175482 | POLD4 |
| ENSG00000175513 | TSGA10IP |
| ENSG00000175564 | UCP3 |
| ENSG00000175643 | RMI2 |
| ENSG00000175662 | TOM1L2 |
| ENSG00000175699 | CCDC197 |
| ENSG00000175701 | LINC00116 |
| ENSG00000175782 | SLC35E3 |
| ENSG00000175832 | ETV4 |
| ENSG00000175866 | BAIAP2 |

| | |
|---|---|
| ENSG00000175894 | TSPEAR |
| ENSG00000175899 | A2M |
| ENSG00000176040 | TMPRSS7 |
| ENSG00000176049 | JAKMIP2 |
| ENSG00000176155 | CCDC57 |
| ENSG00000176204 | LRRTM4 |
| ENSG00000176244 | ACBD7 |
| ENSG00000176261 | ZBTB8OS |
| ENSG00000176438 | SYNE3 |
| ENSG00000176444 | CLK2 |
| ENSG00000176463 | SLCO3A1 |
| ENSG00000176533 | GNG7 |
| ENSG00000176624 | MEX3C |
| ENSG00000176697 | BDNF |
| ENSG00000176715 | ACSF3 |
| ENSG00000176720 | BOK |
| ENSG00000176771 | NCKAP5 |
| ENSG00000176788 | BASP1 |
| ENSG00000176834 | VSIG10 |
| ENSG00000176884 | GRIN1 |
| ENSG00000176909 | MAMSTR |
| ENSG00000177030 | DEAF1 |
| ENSG00000177058 | SLC38A9 |
| ENSG00000177098 | SCN4B |
| ENSG00000177103 | DSCAML1 |
| ENSG00000177106 | EPS8L2 |
| ENSG00000177301 | KCNA2 |
| ENSG00000177311 | ZBTB38 |
| ENSG00000177359 | AC024940.1 |
| ENSG00000177380 | PPFIA3 |
| ENSG00000177398 | UMODL1 |
| ENSG00000177426 | TGIF1 |
| ENSG00000177453 | NIM1K |
| ENSG00000177548 | RABEP2 |
| ENSG00000177565 | TBL1XR1 |
| ENSG00000177675 | CD163L1 |
| ENSG00000177694 | NAALADL2 |
| ENSG00000177728 | TMEM94 |
| ENSG00000177830 | CHID1 |
| ENSG00000177981 | ASB8 |
| ENSG00000177992 | SPATA31E1 |
| ENSG00000178075 | GRAMD1C |
| ENSG00000178078 | STAP2 |
| ENSG00000178096 | BOLA1 |
| ENSG00000178104 | PDE4DIP |
| ENSG00000178188 | SH2B1 |
| ENSG00000178209 | PLEC |
| ENSG00000178226 | PRSS36 |
| ENSG00000178297 | TMPRSS9 |
| ENSG00000178404 | CEP295NL |

| | |
|---|---|
| ENSG00000178662 | CSRNP3 |
| ENSG00000178685 | PARP10 |
| ENSG00000178772 | CPN2 |
| ENSG00000178796 | RIIAD1 |
| ENSG00000178826 | TMEM139 |
| ENSG00000178878 | APOLD1 |
| ENSG00000178882 | RFLNA |
| ENSG00000178935 | ZNF552 |
| ENSG00000179088 | C12orf42 |
| ENSG00000179165 | PXT1 |
| ENSG00000179222 | MAGED1 |
| ENSG00000179242 | CDH4 |
| ENSG00000179335 | CLK3 |
| ENSG00000179348 | GATA2 |
| ENSG00000179397 | CATSPERE |
| ENSG00000179477 | ALOX12B |
| ENSG00000179528 | LBX2 |
| ENSG00000179583 | CIITA |
| ENSG00000179588 | ZFPM1 |
| ENSG00000179715 | PCED1B |
| ENSG00000179846 | NKPD1 |
| ENSG00000179873 | NLRP11 |
| ENSG00000179913 | B3GNT3 |
| ENSG00000179954 | SSC5D |
| ENSG00000180035 | ZNF48 |
| ENSG00000180061 | TMEM150B |
| ENSG00000180316 | PNPLA1 |
| ENSG00000180432 | CYP8B1 |
| ENSG00000180448 | ARHGAP45 |
| ENSG00000180525 | PRR26 |
| ENSG00000180628 | PCGF5 |
| ENSG00000180694 | TMEM64 |
| ENSG00000180745 | CLRN3 |
| ENSG00000180815 | MAP3K15 |
| ENSG00000180822 | PSMG4 |
| ENSG00000180891 | CUEDC1 |
| ENSG00000180999 | C1orf105 |
| ENSG00000181031 | RPH3AL |
| ENSG00000181035 | SLC25A42 |
| ENSG00000181291 | TMEM132E |
| ENSG00000181333 | HEPHL1 |
| ENSG00000181355 | OFCC1 |
| ENSG00000181378 | CFAP65 |
| ENSG00000181381 | DDX60L |
| ENSG00000181409 | AATK |
| ENSG00000181652 | ATG9B |
| ENSG00000181894 | ZNF329 |
| ENSG00000181938 | GINS3 |
| ENSG00000182022 | CHST15 |
| ENSG00000182149 | IST1 |

| | |
|---|---|
| ENSG00000182175 | RGMA |
| ENSG00000182185 | RAD51B |
| ENSG00000182218 | HHIPL1 |
| ENSG00000182253 | SYNM |
| ENSG00000182310 | SPACA6 |
| ENSG00000182326 | C1S |
| ENSG00000182329 | KIAA2012 |
| ENSG00000182372 | CLN8 |
| ENSG00000182472 | CAPN12 |
| ENSG00000182473 | EXOC7 |
| ENSG00000182557 | SPNS3 |
| ENSG00000182568 | SATB1 |
| ENSG00000182578 | CSF1R |
| ENSG00000182580 | EPHB3 |
| ENSG00000182621 | PLCB1 |
| ENSG00000182718 | ANXA2 |
| ENSG00000182752 | PAPPA |
| ENSG00000182771 | GRID1 |
| ENSG00000182795 | C1orf116 |
| ENSG00000182870 | GALNT9 |
| ENSG00000182871 | COL18A1 |
| ENSG00000182885 | ADGRG3 |
| ENSG00000182919 | C11orf54 |
| ENSG00000182957 | SPATA13 |
| ENSG00000183010 | PYCR1 |
| ENSG00000183023 | SLC8A1 |
| ENSG00000183049 | CAMK1D |
| ENSG00000183091 | NEB |
| ENSG00000183092 | BEGAIN |
| ENSG00000183111 | ARHGEF37 |
| ENSG00000183134 | PTGDR2 |
| ENSG00000183145 | RIPPLY3 |
| ENSG00000183230 | CTNNA3 |
| ENSG00000183248 | PRR36 |
| ENSG00000183317 | EPHA10 |
| ENSG00000183337 | BCOR |
| ENSG00000183454 | GRIN2A |
| ENSG00000183463 | URAD |
| ENSG00000183486 | MX2 |
| ENSG00000183520 | UTP11 |
| ENSG00000183682 | BMP8A |
| ENSG00000183778 | B3GALT5 |
| ENSG00000183844 | FAM3B |
| ENSG00000183876 | ARSI |
| ENSG00000183914 | DNAH2 |
| ENSG00000183921 | SDR42E2 |
| ENSG00000183963 | SMTN |
| ENSG00000184009 | ACTG1 |
| ENSG00000184012 | TMPRSS2 |
| ENSG00000184313 | MROH7 |

| | |
|---|---|
| ENSG00000184347 | SLIT3 |
| ENSG00000184368 | MAP7D2 |
| ENSG00000184381 | PLA2G6 |
| ENSG00000184384 | MAML2 |
| ENSG00000184428 | TOP1MT |
| ENSG00000184471 | C1QTNF8 |
| ENSG00000184497 | TMEM255B |
| ENSG00000184613 | NELL2 |
| ENSG00000184702 | sept-05 |
| ENSG00000184792 | OSBP2 |
| ENSG00000184828 | ZBTB7C |
| ENSG00000184922 | FMNL1 |
| ENSG00000184985 | SORCS2 |
| ENSG00000184988 | TMEM106A |
| ENSG00000185015 | CA13 |
| ENSG00000185033 | SEMA4B |
| ENSG00000185038 | MROH2A |
| ENSG00000185049 | NELFA |
| ENSG00000185215 | TNFAIP2 |
| ENSG00000185306 | C12orf56 |
| ENSG00000185324 | CDK10 |
| ENSG00000185332 | TMEM105 |
| ENSG00000185345 | PRKN |
| ENSG00000185420 | SMYD3 |
| ENSG00000185442 | FAM174B |
| ENSG00000185482 | STAC3 |
| ENSG00000185513 | L3MBTL1 |
| ENSG00000185523 | SPATA45 |
| ENSG00000185527 | PDE6G |
| ENSG00000185585 | OLFML2A |
| ENSG00000185630 | PBX1 |
| ENSG00000185640 | KRT79 |
| ENSG00000185651 | UBE2L3 |
| ENSG00000185658 | BRWD1 |
| ENSG00000185666 | SYN3 |
| ENSG00000185681 | MORN5 |
| ENSG00000185739 | SRL |
| ENSG00000185787 | MORF4L1 |
| ENSG00000185792 | NLRP9 |
| ENSG00000185842 | DNAH14 |
| ENSG00000185917 | SETD4 |
| ENSG00000185920 | PTCH1 |
| ENSG00000185973 | TMLHE |
| ENSG00000185974 | GRK1 |
| ENSG00000185989 | RASA3 |
| ENSG00000186007 | LEMD1 |
| ENSG00000186020 | ZNF529 |
| ENSG00000186074 | CD300LF |
| ENSG00000186153 | WWOX |
| ENSG00000186174 | BCL9L |

| | |
|---|---|
| ENSG00000186188 | FFAR4 |
| ENSG00000186197 | EDARADD |
| ENSG00000186350 | RXRA |
| ENSG00000186417 | GLDN |
| ENSG00000186448 | ZNF197 |
| ENSG00000186469 | GNG2 |
| ENSG00000186487 | MYT1L |
| ENSG00000186510 | CLCNKA |
| ENSG00000186517 | ARHGAP30 |
| ENSG00000186567 | CEACAM19 |
| ENSG00000186635 | ARAP1 |
| ENSG00000186642 | PDE2A |
| ENSG00000186648 | CARMIL3 |
| ENSG00000186654 | PRR5 |
| ENSG00000186732 | MPPED1 |
| ENSG00000186765 | FSCN2 |
| ENSG00000186812 | ZNF397 |
| ENSG00000186814 | ZSCAN30 |
| ENSG00000186868 | MAPT |
| ENSG00000186889 | TMEM17 |
| ENSG00000186897 | C1QL4 |
| ENSG00000186918 | ZNF395 |
| ENSG00000186998 | EMID1 |
| ENSG00000187021 | PNLIPRP1 |
| ENSG00000187024 | PTRH1 |
| ENSG00000187079 | TEAD1 |
| ENSG00000187091 | PLCD1 |
| ENSG00000187105 | HEATR4 |
| ENSG00000187122 | SLIT1 |
| ENSG00000187134 | AKR1C1 |
| ENSG00000187147 | RNF220 |
| ENSG00000187164 | SHTN1 |
| ENSG00000187187 | ZNF546 |
| ENSG00000187323 | DCC |
| ENSG00000187391 | MAGI2 |
| ENSG00000187498 | COL4A1 |
| ENSG00000187556 | NANOS3 |
| ENSG00000187595 | ZNF385C |
| ENSG00000187608 | ISG15 |
| ENSG00000187609 | EXD3 |
| ENSG00000187624 | C17orf97 |
| ENSG00000187630 | DHRS4L2 |
| ENSG00000187634 | SAMD11 |
| ENSG00000187672 | ERC2 |
| ENSG00000187688 | TRPV2 |
| ENSG00000187699 | C2orf88 |
| ENSG00000187720 | THSD4 |
| ENSG00000187726 | DNAJB13 |
| ENSG00000187730 | GABRD |
| ENSG00000187772 | LIN28B |

| | |
|---|---|
| ENSG00000187775 | DNAH17 |
| ENSG00000187783 | TMEM72 |
| ENSG00000187800 | PEAR1 |
| ENSG00000187908 | DMBT1 |
| ENSG00000187955 | COL14A1 |
| ENSG00000188001 | TPRG1 |
| ENSG00000188037 | CLCN1 |
| ENSG00000188089 | PLA2G4E |
| ENSG00000188095 | MESP2 |
| ENSG00000188158 | NHS |
| ENSG00000188243 | COMMD6 |
| ENSG00000188282 | RUFY4 |
| ENSG00000188372 | ZP3 |
| ENSG00000188385 | JAKMIP3 |
| ENSG00000188522 | FAM83G |
| ENSG00000188549 | C15orf52 |
| ENSG00000188603 | CLN3 |
| ENSG00000188677 | PARVB |
| ENSG00000188687 | SLC4A5 |
| ENSG00000188735 | TMEM120B |
| ENSG00000188779 | SKOR1 |
| ENSG00000188878 | FBF1 |
| ENSG00000188897 | AC099489.1 |
| ENSG00000188937 | NYX |
| ENSG00000188981 | MSANTD1 |
| ENSG00000189001 | SBSN |
| ENSG00000189045 | ANKDD1B |
| ENSG00000189067 | LITAF |
| ENSG00000189120 | SP6 |
| ENSG00000189143 | CLDN4 |
| ENSG00000189144 | ZNF573 |
| ENSG00000189157 | FAM47E |
| ENSG00000189159 | JPT1 |
| ENSG00000189233 | NUGGC |
| ENSG00000189337 | KAZN |
| ENSG00000189350 | TOGARAM2 |
| ENSG00000189403 | HMGB1 |
| ENSG00000196092 | PAX5 |
| ENSG00000196123 | KIAA0895L |
| ENSG00000196132 | MYT1 |
| ENSG00000196139 | AKR1C3 |
| ENSG00000196154 | S100A4 |
| ENSG00000196208 | GREB1 |
| ENSG00000196218 | RYR1 |
| ENSG00000196220 | SRGAP3 |
| ENSG00000196235 | SUPT5H |
| ENSG00000196284 | SUPT3H |
| ENSG00000196313 | POM121 |
| ENSG00000196358 | NTNG2 |
| ENSG00000196405 | EVL |

| | |
|---|---|
| ENSG00000196417 | ZNF765 |
| ENSG00000196421 | C20orf204 |
| ENSG00000196431 | CRYBA4 |
| ENSG00000196470 | SIAH1 |
| ENSG00000196482 | ESRRG |
| ENSG00000196531 | NACA |
| ENSG00000196562 | SULF2 |
| ENSG00000196565 | HBG2 |
| ENSG00000196569 | LAMA2 |
| ENSG00000196576 | PLXNB2 |
| ENSG00000196591 | HDAC2 |
| ENSG00000196597 | ZNF782 |
| ENSG00000196605 | ZNF846 |
| ENSG00000196628 | TCF4 |
| ENSG00000196639 | HRH1 |
| ENSG00000196660 | SLC30A10 |
| ENSG00000196684 | HSH2D |
| ENSG00000196743 | GM2A |
| ENSG00000196872 | KIAA1211L |
| ENSG00000196946 | ZNF705A |
| ENSG00000196950 | SLC39A10 |
| ENSG00000196975 | ANXA4 |
| ENSG00000196998 | WDR45 |
| ENSG00000197046 | SIGLEC15 |
| ENSG00000197056 | ZMYM1 |
| ENSG00000197181 | PIWIL2 |
| ENSG00000197183 | NOL4L |
| ENSG00000197223 | C1D |
| ENSG00000197283 | SYNGAP1 |
| ENSG00000197361 | FBXL22 |
| ENSG00000197405 | C5AR1 |
| ENSG00000197415 | VEPH1 |
| ENSG00000197444 | OGDHL |
| ENSG00000197448 | GSTK1 |
| ENSG00000197471 | SPN |
| ENSG00000197536 | C5orf56 |
| ENSG00000197558 | SSPO |
| ENSG00000197580 | BCO2 |
| ENSG00000197653 | DNAH10 |
| ENSG00000197702 | PARVA |
| ENSG00000197757 | HOXC6 |
| ENSG00000197816 | CCDC180 |
| ENSG00000197852 | FAM212B |
| ENSG00000197872 | FAM49A |
| ENSG00000197879 | MYO1C |
| ENSG00000197893 | NRAP |
| ENSG00000197912 | SPG7 |
| ENSG00000197943 | PLCG2 |
| ENSG00000197959 | DNM3 |
| ENSG00000197971 | MBP |

| | |
|---|---|
| ENSG00000197977 | ELOVL2 |
| ENSG00000198039 | ZNF273 |
| ENSG00000198055 | GRK6 |
| ENSG00000198089 | SFI1 |
| ENSG00000198125 | MB |
| ENSG00000198130 | HIBCH |
| ENSG00000198133 | TMEM229B |
| ENSG00000198157 | HMGN5 |
| ENSG00000198216 | CACNA1E |
| ENSG00000198336 | MYL4 |
| ENSG00000198353 | HOXC4 |
| ENSG00000198373 | WWP2 |
| ENSG00000198431 | TXNRD1 |
| ENSG00000198517 | MAFK |
| ENSG00000198551 | ZNF627 |
| ENSG00000198586 | TLK1 |
| ENSG00000198598 | MMP17 |
| ENSG00000198626 | RYR2 |
| ENSG00000198646 | NCOA6 |
| ENSG00000198663 | C6orf89 |
| ENSG00000198673 | FAM19A2 |
| ENSG00000198691 | ABCA4 |
| ENSG00000198715 | GLMP |
| ENSG00000198719 | DLL1 |
| ENSG00000198720 | ANKRD13B |
| ENSG00000198723 | TEX45 |
| ENSG00000198728 | LDB1 |
| ENSG00000198729 | PPP1R14C |
| ENSG00000198732 | SMOC1 |
| ENSG00000198796 | ALPK2 |
| ENSG00000198807 | PAX9 |
| ENSG00000198821 | CD247 |
| ENSG00000198851 | CD3E |
| ENSG00000198873 | GRK5 |
| ENSG00000198898 | CAPZA2 |
| ENSG00000198910 | L1CAM |
| ENSG00000198915 | RASGEF1A |
| ENSG00000198945 | L3MBTL3 |
| ENSG00000198947 | DMD |
| ENSG00000198948 | MFAP3L |
| ENSG00000198959 | TGM2 |
| ENSG00000203499 | IQANK1 |
| ENSG00000203697 | CAPN8 |
| ENSG00000203814 | HIST2H2BF |
| ENSG00000203867 | RBM20 |
| ENSG00000203985 | LDLRAD1 |
| ENSG00000204060 | FOXO6 |
| ENSG00000204131 | NHSL2 |
| ENSG00000204140 | CLPSL1 |
| ENSG00000204209 | DAXX |

| | |
|---|---|
| ENSG00000204248 | COL11A2 |
| ENSG00000204262 | COL5A2 |
| ENSG00000204396 | VWA7 |
| ENSG00000204397 | CARD16 |
| ENSG00000204516 | MICB |
| ENSG00000204520 | MICA |
| ENSG00000204540 | PSORS1C1 |
| ENSG00000204580 | DDR1 |
| ENSG00000204616 | TRIM31 |
| ENSG00000204628 | RACK1 |
| ENSG00000204634 | TBC1D8 |
| ENSG00000204767 | FAM196B |
| ENSG00000204815 | TTC25 |
| ENSG00000204866 | IGFL2 |
| ENSG00000204882 | GPR20 |
| ENSG00000204920 | ZNF155 |
| ENSG00000204954 | C12orf73 |
| ENSG00000204960 | BLACE |
| ENSG00000204991 | SPIRE2 |
| ENSG00000205078 | SYCE1L |
| ENSG00000205086 | C2orf91 |
| ENSG00000205133 | TRIQK |
| ENSG00000205212 | CCDC144NL |
| ENSG00000205309 | NT5M |
| ENSG00000205336 | ADGRG1 |
| ENSG00000205639 | MFSD2B |
| ENSG00000205726 | ITSN1 |
| ENSG00000205744 | DENND1C |
| ENSG00000205795 | CYS1 |
| ENSG00000205832 | C16orf96 |
| ENSG00000206077 | ZDHHC11B |
| ENSG00000206190 | ATP10A |
| ENSG00000206561 | COLQ |
| ENSG00000211584 | SLC48A1 |
| ENSG00000212719 | C17orf51 |
| ENSG00000212901 | KRTAP3-1 |
| ENSG00000213160 | KLHL23 |
| ENSG00000213213 | CCDC183 |
| ENSG00000213397 | HAUS7 |
| ENSG00000213445 | SIPA1 |
| ENSG00000213780 | GTF2H4 |
| ENSG00000213889 | PPM1N |
| ENSG00000213903 | LTB4R |
| ENSG00000213949 | ITGA1 |
| ENSG00000213999 | MEF2B |
| ENSG00000214026 | MRPL23 |
| ENSG00000214050 | FBXO16 |
| ENSG00000214226 | C17orf67 |
| ENSG00000214357 | NEURL1B |
| ENSG00000214456 | PLIN5 |

| | |
|---|---|
| ENSG00000214491 | SEC14L6 |
| ENSG00000214688 | C10orf105 |
| ENSG00000214711 | CAPN14 |
| ENSG00000214814 | FER1L6 |
| ENSG00000214944 | ARHGEF28 |
| ENSG00000214954 | LRRC69 |
| ENSG00000215018 | COL28A1 |
| ENSG00000215045 | GRID2IP |
| ENSG00000215182 | MUC5AC |
| ENSG00000215252 | GOLGA8B |
| ENSG00000215271 | HOMEZ |
| ENSG00000215440 | NPEPL1 |
| ENSG00000215529 | EFCAB8 |
| ENSG00000215910 | C1orf167 |
| ENSG00000215912 | TTC34 |
| ENSG00000217930 | PAM16 |
| ENSG00000219626 | FAM228B |
| ENSG00000221823 | PPP3R1 |
| ENSG00000221866 | PLXNA4 |
| ENSG00000221909 | FAM200A |
| ENSG00000221986 | MYBPHL |
| ENSG00000224383 | PRR29 |
| ENSG00000224470 | ATXN1L |
| ENSG00000225526 | MKRN2OS |
| ENSG00000225940 | C5orf67 |
| ENSG00000225968 | ELFN1 |
| ENSG00000226174 | TEX22 |
| ENSG00000226321 | CROCC2 |
| ENSG00000226690 | AC013470.2 |
| ENSG00000227471 | AKR1B15 |
| ENSG00000231672 | DIRC3 |
| ENSG00000232119 | MCTS1 |
| ENSG00000234616 | JRK |
| ENSG00000234828 | IQCM |
| ENSG00000234965 | SHISA8 |
| ENSG00000235109 | ZSCAN31 |
| ENSG00000235162 | C12orf75 |
| ENSG00000235568 | NFAM1 |
| ENSG00000236383 | LINC00854 |
| ENSG00000236699 | ARHGEF38 |
| ENSG00000239605 | STPG4 |
| ENSG00000240720 | LRRD1 |
| ENSG00000241489 | AC244197.3 |
| ENSG00000241644 | INMT |
| ENSG00000241697 | TMEFF1 |
| ENSG00000241935 | HOGA1 |
| ENSG00000241962 | AC079447.1 |
| ENSG00000242612 | DECR2 |
| ENSG00000242852 | ZNF709 |
| ENSG00000243156 | MICAL3 |

| | |
|---|---|
| ENSG00000243660 | ZNF487 |
| ENSG00000243696 | AC006254.1 |
| ENSG00000243709 | LEFTY1 |
| ENSG00000243710 | CFAP57 |
| ENSG00000243811 | APOBEC3D |
| ENSG00000243910 | TUBA4B |
| ENSG00000243978 | RTL9 |
| ENSG00000244122 | UGT1A7 |
| ENSG00000244405 | ETV5 |
| ENSG00000244486 | SCARF2 |
| ENSG00000244607 | CCDC13 |
| ENSG00000246922 | UBAP1L |
| ENSG00000248405 | PRR5-ARHGAP8 |
| ENSG00000248767 | AC187653.1 |
| ENSG00000249209 | AP000311.1 |
| ENSG00000249715 | FER1L5 |
| ENSG00000249884 | RNF103-CHMP3 |
| ENSG00000249961 | TERB1 |
| ENSG00000250317 | SMIM20 |
| ENSG00000250506 | CDK3 |
| ENSG00000250722 | SELENOP |
| ENSG00000251287 | ALG1L2 |
| ENSG00000251322 | SHANK3 |
| ENSG00000253309 | SERPINE3 |
| ENSG00000254402 | LRRC24 |
| ENSG00000254827 | SLC22A18AS |
| ENSG00000255346 | NOX5 |
| ENSG00000255508 | AP002990.1 |
| ENSG00000255872 | AL138752.2 |
| ENSG00000256061 | DNAAF4 |
| ENSG00000256087 | ZNF432 |
| ENSG00000256806 | C17orf100 |
| ENSG00000257335 | MGAM |
| ENSG00000257743 | MGAM2 |
| ENSG00000257923 | CUX1 |
| ENSG00000257987 | TEX49 |
| ENSG00000258102 | MAP1LC3B2 |
| ENSG00000258461 | AC012651.1 |
| ENSG00000258472 | AC005726.2 |
| ENSG00000258659 | TRIM34 |
| ENSG00000259316 | AC087632.1 |
| ENSG00000259417 | CTXND1 |
| ENSG00000260027 | HOXB7 |
| ENSG00000260220 | CCDC187 |
| ENSG00000260230 | FRRS1L |
| ENSG00000260300 | AC009119.2 |
| ENSG00000260314 | MRC1 |
| ENSG00000260456 | C16orf95 |
| ENSG00000261341 | AC010325.1 |
| ENSG00000261371 | PECAM1 |

| | |
|---|---|
| ENSG00000261582 | AL121753.1 |
| ENSG00000263429 | LINC00675 |
| ENSG00000263639 | MSMB |
| ENSG00000263961 | C1orf186 |
| ENSG00000264230 | ANXA8L1 |
| ENSG00000264324 | AC006030.1 |
| ENSG00000265190 | ANXA8 |
| ENSG00000266074 | BAHCC1 |
| ENSG00000266076 | AC004805.1 |
| ENSG00000266094 | RASSF5 |
| ENSG00000266714 | MYO15B |
| ENSG00000267561 | AC093155.3 |
| ENSG00000269113 | TRABD2B |
| ENSG00000269404 | SPIB |
| ENSG00000269693 | AC010422.6 |
| ENSG00000270106 | TSNAX-DISC1 |
| ENSG00000270647 | TAF15 |
| ENSG00000271447 | MMP28 |
| ENSG00000271605 | MILR1 |
| ENSG00000272305 | AC096887.1 |
| ENSG00000272636 | DOC2B |
| ENSG00000272899 | AC025594.2 |
| ENSG00000273167 | AL359736.1 |
| ENSG00000273259 | AL049839.2 |
| ENSG00000273398 | AC017083.4 |
| ENSG00000273899 | NOL12 |
| ENSG00000274070 | CASTOR2 |
| ENSG00000274322 | AL136531.2 |
| ENSG00000275832 | ARHGAP23 |
| ENSG00000276043 | UHRF1 |
| ENSG00000276600 | RAB7B |
| ENSG00000277363 | SRCIN1 |
| ENSG00000278500 | AC009336.2 |
| ENSG00000278540 | ACACA |
| ENSG00000278570 | NR2E3 |
| ENSG00000281106 | LINC00282 |
| ENSG00000283154 | IQCJ-SCHIP1 |
| ENSG00000283297 | AC005841.2 |
| ENSG00000283782 | AC116366.3 |
| ENSG00000283900 | Z98749.3 |
| ENSG00000284461 | RABGEF1 |
| ENSG00000284686 | AC119674.2 |
| ENSG00000284691 | AC073111.5 |

#### A Genotyping primers

| Gene | Forward primers | Reverse primers |
|---|---|---|
| **Tail samples** | | |
| *wild-type or floxed H2afz* | CGCCTTGGTAATTCTATCTTCTCC | CGCCAGTTAACACACATGTGATC |
| *wild-type or floxed H2afv* | GCCTCAGATCATCCAGTC | GGCTCTGAATTCCCAATGTAG |
| wild-type Lgr5 | ATACCCCATCCCTTTTGAGC | CTGCTCTCTGCTCCCAGTCT |
| Lgr5-CRE insert | GAACTTCAGGGTCAGCTTGC | CTGCTCTCTGCTCCCAGTCT |
| **Intestinal epithelium** | | |
| *wild-type and (native or recombined) floxed H2afz* | CGCCTTGGTAATTCTATCTTCTCC | AAGCCTCCAACTTGCTCAAA |
| *wild-type and (native or recombined) floxed H2afv* | GCCTCCAACTACAAATGCTC | GGCTCTGAATTCCCAATGTAG |

#### B siRNA sequence

| siRNA | Sequence |
|---|---|
| Ctrl | ACUCAAACUCACGAAGGAA |
| H2A.Z.1 | GUAGUGGGUUUUGAUUGAG |
| H2A.Z.2 | GGAAAAGCAUAGACAAUUA |
| P400 | UGAAGAAGGUUCCCAAGAA |
| SRCAP | GGAAACGAUUGAAGUUGAA |

#### C ChIP primers

| Gene promoter | Forward primers | Reverse primers |
|---|---|---|
| SI | AATAGTTCACAGCTTTGAGAAATCA | GATCAAGGAAAGCTGCTTAGG |
| *LPH* | AAAATTAGCCAGGCATCGTG | TTCAGACATTTTCCGGGTTC |
| *CDKN1A*$^{p21}$ | GTGGCTCTGATTGGCTTTCTG | CTGAAAACAGGCAGCCCAAGG |
| *LGR5* | AGAGAGCGCTGGGACACTTA | ACTCCACTGCTGCCTTCCTA |
| *MMP7* | ATTTCCACATTCGAGGCTGA | GCCTGTTCCCACTGTAGCTC |
| *ATOH1* | GCAACCGGAGAAGCATAGTT | TGGTCTCCCAACTCCTTCAC |
| *RPLP0* | GGCGACCTGGAAGTCCAACT | CCATCAGCACCACAGCCTTC |
| *GAPDH intron 8* | TATGTGGCTGGGGCCAGAGA | CAGGGCCCTTTTTCTGAGCC |
| *KCNH6* | CGGAAGTTCCTGATTGCCAA | GGTAGTAGAGGATGTCCACC |

### D CRISPR-Cas9 tools

| | |
|---|---|
| ***H2AFZ*** | |
| Primer guides | ACACCGCCTGAGATAACAAGGAATCCG<br>AAAACGGATTCCTTGTTATCTCAGGCG |
| Screening primers | AGCGTATTACCCCTCGTCAC<br>TGTGCTTAGTTATTGCTGCTAGA |
| ***H2AFV*** | |
| Primer guides | ACACCGCCAGTTACAGTACAATGACGG<br>AAAACCGTCATTGTACTGTAACTGGCG |
| Screening primers | CCTTCATTTGATATGCTAATCATGG<br>CACCACATGAAAAGGTACTTTTATC |

### E RT-qPCR primers

#### Human primers

| Gene | Forward primers | Reverse primers |
|---|---|---|
| *β2M* | AAAGATGAGTATGCCTGCCG | CCTCCATGATGCTGCTTACA |
| *RPLP0* | GGCGACCTGGAAGTCCAACT | CCATCAGCACCACAGCCTTC |
| *H2A.Z.1* | CCTTTTCTCTGCCTTGCTTG | CGGTGAGGTACTCCAGGATG |
| *H2A.Z.2* | TCCCTCACATCCACAAATCTC | AGTACAATGACGGGGAGGAA |
| *SI* | ATGTGAAGGTTGCCCAAAAC | AAAATTGGCCATGTTTTCCA |
| *LPH* | CCAGGAGATATGTTCAGTTC | CACTCTCTGTAAATTCTGGC |
| *MUCDHL L* | CTCCCACCAACCAACCAC | CATATCCACCACCGAGAAGC |
| *MUCDHL M* | TGGAGGGAGAGGTTGTGCT | GGCCGCCACCTGTGGAGG |
| *CDKN1A*$^{p21}$ | GTCAGAACCGGCTGGGGATG | TGAGCGAGGCACAAGGGTAC |
| *LGR5* | TGCTCTTCACCAACTGCATC | CTCAGGCTCACCAGATCCTC |

#### Murine primers

| Gene | Forward primers | Reverse primers |
|---|---|---|
| *B2m* | CCTGGTCTTTCTGGTGCTTG | TATGTTCGGCTTCCCATTCT |
| *Rplp0* | GGCGACCTGGAAGTCCAACT | CCATCAGCACCACAGCCTTC |
| *H2a.z.1* | CCGTATTCATCGACACCTGA | AAGTGACGAGGGGTGATACG |
| *H2a.z.2* | CGCATCCACAGACACTTGAA | TCAGCTGTGAGGTACTCCAGAA |
| *Si* | GCTGGTCGATGGGGAGGA | CCAACGAGCACAGAGGTGGTAT |
| *Lph* | CCTTGAGCCCAAAGTGAAAG | GGACGTACAGCTCAGGAAGG |
| *Muc2* | TGATGGCCATTGAGGTGGAG | CTGGCCCTTTGTGTTGTTGC |
| *Math1* | GCTTCCTCTGGGGGTTACTC | CTGTGGGATCTGGGAGATGT |
| *Hes1* | CCAGCCAGTGTCAACACGA | AATGCCGGGAGCTATCTTTCT |
| *Lgr5* | AGGCTGCCAAAAACTTCAGA | TAACCCAGTCACAGGGAAGG |
| *Olfm4* | GCACCTGCCAGTGTTCTGTT | GACCTCTACTCGGACCGTCA |

__Supplementary Table 3__ : Technical tools and sequences

A) Genotyping primer sequences. B) Sequences of siRNAs. C) Sequences of primers used in ChIP experiments. D) Tools used in Caco-2/15 cells for CRISPR-Cas9 strategy to tag and screen *H2AFZ* or *H2AFV* coding sequences on their respective genes. E) RT-qPCR primer sequences.
